# 3’-5’ crosstalk contributes to transcriptional bursting

**DOI:** 10.1101/514174

**Authors:** Massimo Cavallaro, Mark D. Walsh, Matt Jones, James Teahan, Simone Tiberi, Bärbel Finkenstädt, Daniel Hebenstreit

**Affiliations:** School of Life Sciences, University of Warwick, Coventry, UK; Mathematics Institute and Zeeman Institute for Systems Biology and Infectious Disease Epidemiology Research, University of Warwick, Coventry, UK; Department of Statistics, University of Warwick, Coventry, UK; Department of Chemistry, University of Warwick, Coventry, UK; Institute of Molecular Life Sciences and Swiss Institute of Bioinformatics, University of Zurich, Zurich, Switzerland

**Keywords:** gene expression, parameter inference, mathematical modelling, gene looping, biological noise, liquid-liquid phase separation

## Abstract

**Background:** Transcription in mammalian cells is a complex stochastic process involving shuttling of polymerase between genes and phase-separated liquid condensates. It occurs in bursts, which results in vastly different numbers of an mRNA species in isogenic cell populations. Several factors contributing to transcriptional bursting have been identified, usually classified as intrinsic, in other words local to single genes, or extrinsic, relating to the macroscopic state of the cell. However, some possible contributors have not been explored yet. Here, we focus on processes at the 3’ and 5’ ends of a gene that enable reinitiation of transcription upon termination.

**Results:** Using Bayesian methodology, we measure the transcriptional bursting in inducible transgenes, showing that perturbation of polymerase shuttling typically reduces burst size, increases burst frequency, and thus limits transcriptional noise. Analysis based on paired-end tag sequencing (PolII ChIA-PET) suggests that this effect is genome wide. The observed noise patterns are also reproduced by a generative model that captures major characteristics of the polymerase flux between the ends of a gene and a phase-separated compartment.

**Conclusions:** Interactions between the 3’ and 5’ ends of a gene, which facilitate polymerase recycling, are major contributors to transcriptional noise.

## Introduction

In many cellular systems, mRNAs appear to be produced in burst-like fashion. This is directly observed in real-time experimental studies [1, 2, 3] and also agrees with theoretical analyses of steady-state mRNA distributions among single cells [4, 5]. Such bursty dynamics are thought to be the signature of gene regulation and are often described in terms of transcriptional “noise” [6, 5]. Due to the central role of transcription in cellular functions, it is important to understand the mechanisms from which the bursting originates [7].

The microscopic dynamics underlying transcription are not yet well understood. Various factors have been found to influence transcriptional dynamics, mostly by modulating bursting parameters such as the size or frequency of bursts [3, 5]. These factors are often classified as either intrinsic or extrinsic, although this distinction is blurred in many cases. This classification originally derives from the observation that fluctuations in expression levels are partially correlated across multiple genes [8], thus suggesting common, extrinsic causes, while the remaining, independent fluctuations are intrinsic to each gene. Typical major extrinsic noise sources are the cell cycle [9, 10, 11] and cell-size fluctuations [12], the latter partially due to the former. Numerous additional factors such as neighbouring cells, cell morphology and others have been found to affect transcription to varying degrees [13]. Intrinsic factors include non-linear transcription factor interactions [4, 5, 8], changing chromatin status [14, 15], promoter architecture [3], transcription factor diffusion [16], and several others [17, 18, 19].

It is unclear how these phenomena relate to the local environment at transcribing genes. These are associated to clusters of RNA polymerase II (PolII), which have been interpreted as “transcription factories” [20] and suggested to modulate the temporal patterns of transcription [21, 22]. More recently, it has been found that, in proximity to active genes, the PolIIs are incorporated in membrane-less droplets, maintained by liquid-liquid phase separation (LLPS) from the rest of the nucleus, with the net effect of locally increasing the population of factors involved in initiation; when PolII is liberated from this domain, transcription can be initiated [23, 24, 25, 26, 27, 28]. LLPS also provides an explanation for the hitherto enigmatic action-at-adistance type of gene regulation by distal enhancers, as the nuclear condensates are indeed able to restructure the genome, albeit results on LLPS are relatively preliminary at this stage [29, 30].

While a comprehensive description of the interactions between PolIIs, other factors, and the chromatin within these niches is missing, several observations suggest that termination is linked to reinitiation; these include the presence of the same factor species at both ends of a gene, the reduction of initiation upon perturbation of 3’ processes, and protein interactions that have been suggested to juxtapose the promoter and the terminator DNA, forming a structure that has been referred to as a “gene loop” [31, 32]. Importantly, it has been demonstrated that 3’-end processing favours transcription initiation; the presence of such 3’-5’ crosstalk in a gene increases its mean expression level [33]. The concept of LLPS appears highly important in this regard, as PolII undergoes a sequence of post-translational modifications on its C-terminal domain during transcription, while integration into phase separated domains and reinitiation requires it to be unmodified [24]. In line with this, recent studies suggest that LLPS is also involved in 3’-end transcriptional processes [34]. We generically refer to the shuttling of PolIIs from 5’ to 3’, potentially passing through the LLPS compartment, as the recycling. It has been suggested that a repetitive cycle of reinitiation and termination due to these mechanisms is likely to produce a rapid succession of mRNA creation events, thus potentially contributing to the transcriptional bursts [35] but, to the best of our knowledge, an experimental verification is as yet lacking.

In this paper, we investigate the interplay between bursty expression and 3’-5’ interactions using an inter-disciplinary approach. We first consider two integrated genes that permit studying transcription upon perturbation of their 3’-5’ processes at different induction levels; we demonstrate that these interactions strikingly influence the transcription kinetics and typically elicit the transcriptional noise, by decreasing burst frequency and increasing burst size. We then focus on genome-wide 3’-5’ interactions involved in transcription by means of PolII ChIA-PET sequencing data, showing that they are related to the gene-expression parameters similarly to the transgenes’ results. This scenario is well described by a microscopic stochastic model of gene expression, where tuning a single parameter—corresponding to the probability of local polymerase recycling—naturally yields the observed expression patterns, without involving extrinsic-noise contributors or alternative intrinsic mechanisms.

## Results

### Cell lines as model systems for PolII recycling

We utilized two HEK293 cell lines which contain on their genomes copies of the genes *β*-*globin* (HBB) [33] and a modified version of HIV-1-*env* [36], respectively, driven by inducible CMV promoters (Figure 1 A and B). This transgene approach allowed us to exploit very well characterized model systems for recycling perturbation, which achieve mono-allelic expression and, most importantly, allow precise control of expression levels with inducers [33]. The first gene, HBB, is an example for long-range chromosomal interactions in its native genomic neighbourhood. Its expression involves spatial proximity between the promoter and a locus control region (LCR) over 50 Kbs away [37]. The LCR has been studied extensively in murine and human cells (see, e.g., references [38, 39]) and jointly regulates expression of several *β*-*globin* like genes at the locus, likely involving LLPS [40]. A recent study demonstrates burst-like expression of murine HBB and suggests that interactions between the LCR and the HBB promoter modulate the bursting parameters [9]. Our cell line features an ectopic insertion of human HBB under control of a tetracycline (Tet) responsive promoter. A previous study of this system has provided a substantial number of results suggesting that 3’ mRNA processing contributes to reinitiation of transcription [33]. This notion is based on several findings relating to the introduction of a single point mutation in the SV40 late poly-adenylation (pA) site (Figure 1 C). This includes decreased average mRNA expression levels, while “read-through” transcription downstream of the pA site is increased. Furthermore, the mutation leads to a decrease of PolII, TBP and TFIIB levels at the promoter shortly after gene induction, and to an accumulation at the “read-through” region instead. Reduced transcription initiation compared to wild-type (WT) cells was also supported by nuclear run on assays and by a changed profile of post-translational modifications of PolII. Noticeably, TFIIB has been demonstrated to be functionally involved in linking 3’ and 5’ transcriptional activities [41], while post-translational modifications of PolII are in part carried out by Ssu72, which is associated with gene-loop formation in yeast [42] and appears to have similar roles in vertebrates [43]. A further recent study that utilized the ectopic HBB system reports direct detection of gene loops based on a 3C assay in the WT cell line, but not the mutant [44].

**Figure 1.**
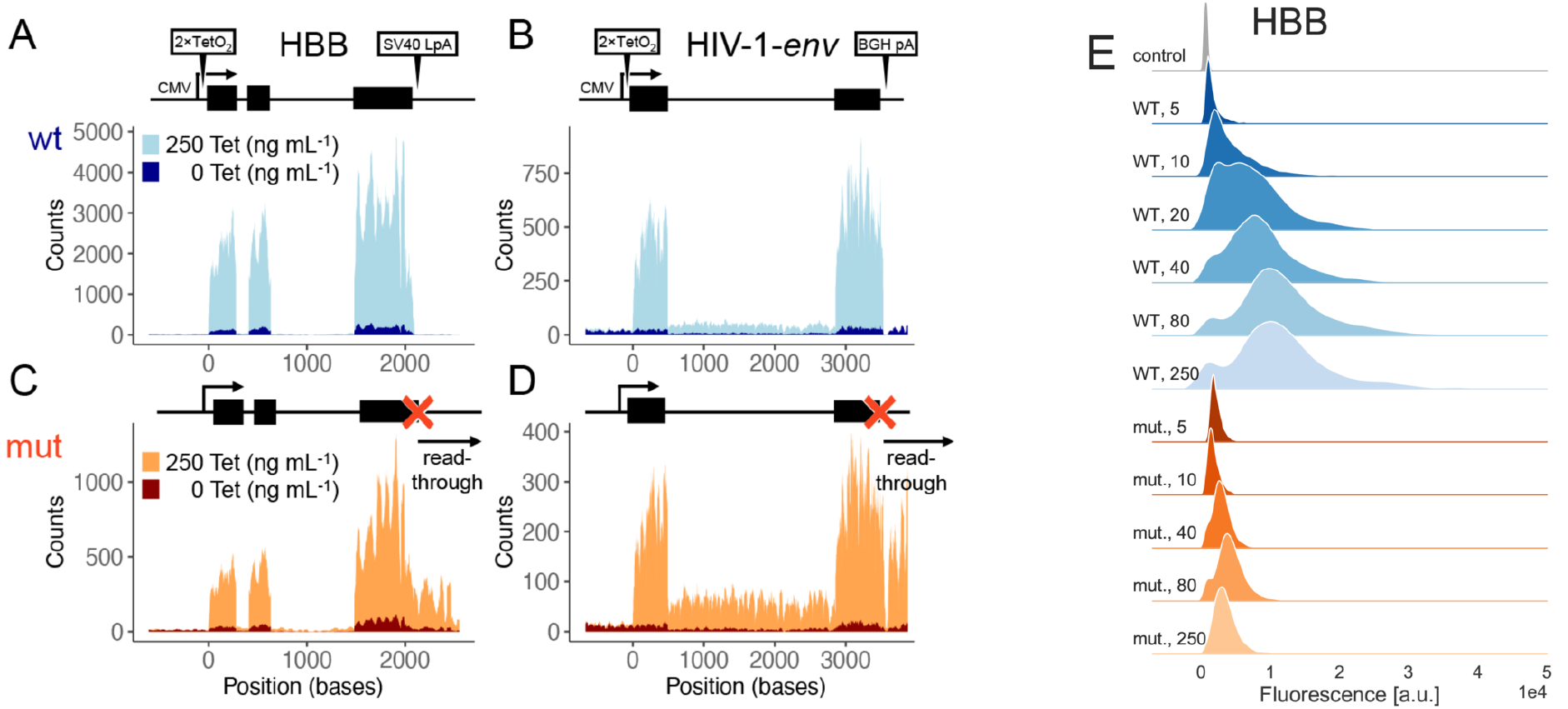
Characteristics of transgenes used in this study. (A) schematic gene structure (top) of WT HBB including CMV promoter, Tet operator, pA signal, and exons (black blocks) as indicated. Total RNA-seq confirms Tet-inducible expression (bottom). (B) as (A), for *env*. (C)-(D) mutant versions of HBB and *env*, respectively. Point mutations in pA sites (‘x’) and read-through transcription are indicated. Positions are relative to TSS, blue and orange shades correspond to WT and mutant versions, respectively, and light and dark shades correspond to 250 and 0 ng mL^−1^ Tet, respectively. Coverages by sequencing reads are shown. (E) Kernel density estimates of the flow-FISH single-cell readings corresponding to the abundances of HBB transcripts, WT (blue), mutant (orange) variants, and control (gray) cells, from replicate *k* = 1, at different induction levels (Tet concentrations in unit of ng mL^−1^, shades of colors, as indicated on the left). Gene expression increases and saturates upon increasing Tet concentration, mutant-cell expression is lower than the WT; a.u., arbitrary units; *y*-axes not to scale.

The second cell line, containing a Tet-inducible version of HIV-1-*env*, was previously studied in similar fashion to the HBB constructs. Results using a mutated version of the pA site (Figure 1 D) mirrored those obtained with HBB, suggesting extensive 3’-5’ crosstalk and recycling of factors including polymerase [33, 45]. The *env* construct uses a BGH, not an SV40 pA site, which suggests that the findings are independent of the type of pA site. Notably, expression of the HIV-1 gene using its native long terminal repeat (LTR) promoter exhibits bursting dynamics [6].

We used these cell lines and their mutant versions as a model system for mammalian gene expression in presence and absence of 3’-5’ crosstalk. We confirmed by total RNA-seq that HBB and *env* mRNAs are expressed inducibly in all cell lines (Figure 1 A-D). At high Tet concentration (250 ng mL^−1^), the fold changes over the un-induced samples were ≈ 16 and ≈ 26 for HBB and *env*, respectively. The mutants were expressed at lower levels and featured read-through transcription as described, with intact transcript sequences, i.e., not subject to splicing defects (Figure 1 C-D). This indicated specificity of the pA site mutations.

In order to detect transcripts at the single molecule level, we designed probes for single molecule RNA-FISH (smFISH) and confirmed detection of large transcript numbers upon Tet stimulation of the cells, while the expression of a control gene, AKT1, remained constant (Additional file 1: section S1 and Fig. S3). Microscopy-based smFISH is not ideal for HEK293 cells, since they tend to overlap and form aggregates when growing. We therefore decided to record the sm-FISH signal by adapting a flow-FISH technique based on flow cytometry [46]; this also resolves extrinsic-noise contributors such as cell size, morphology, and cycle, and, thanks to its high throughput, permits recording vast numbers of cells to analyse overall population structures (Additional file 1: sections S1 and S7).

While the flow-cytometer fluorescence signal from stained cells serves as a proxy for the mRNA abundance, it is returned in arbitrary units (a.u.) rather than in absolute counts. We thus used microscope imaging and nCounter® data to calibrate the flow FISH fluorescence readings of HBB and *env* cells, respectively. Applying the clustering algorithm of [47] to the flow-FISH recordings allowed us to select single-cell readings against those from cell clumps, doublets, and debris (Additional file 1: section S1 and Fig. S1).

Flow-FISH data demonstrate Tet-dose dependent expression of HBB and *env*, indicating specific detection of transcripts above background noise. The stationary expression levels appeared to reach saturation at 80 ng mL^−1^ Tet (Figure 1 E and Additional file 1: section S1 and Fig. S2). Staining for the DNA content demonstrates a mild increase of HBB and *env* expression with increasing cell cycle stage. We found that the contribution to the total variability, measured as the squared coefficient of variation (CV^2^) of the mRNA population, due to the cell cycle and size was minor (Additional file 1: section S6) and therefore focused on local genic mechanisms to investigate the observed noise pattern. The measured signal includes a background of unspecific staining and auto-fluorescence of the cells, which is subtracted from the total signal [48]. To gauge this background we deleted the *env* gene from its host cell line with Cas9 and performed the staining procedure as before. The resulting control cells had low fluorescence intensity that remained virtually unchanged upon maximal Tet stimulation, thus confirming specificity of our system and validating the use of this control to estimate the background (Additional file 1: section S1 Table S1). Nuclear RNA export was largely unaltered by the mutations (Wilcoxon rank sum test on nuclear/ cytoplasmic ratios from 83 HBB cells at 250ng mL^−1^ Tet, P=0.85; for 203 *env* cells P=6 · 10^−9^, but the a ratios differed only by 10%). Note that FlowFISH and their analysis/interpretation are unaffected by nuclear export issues.

### Increased transcriptional bursting upon 3’-5’ crosstalk

In order to gain insights into the transcriptional dynamics driving WT and mutant expression of HBB and *env*, we employed a Markov chain Monte Carlo (MCMC) sampling approach to fit statistical models to the flow-FISH data (Figure 2). Importantly, Bayesian modelling permitted using microscope and nCounter® data to estimate informative prior distributions that calibrate the absolute mRNA quantification, whilst retaining flexibility in this respect. We further incorporated the background signal in the Bayesian framework based on the estimates from the Tet-stimulated control cells (*Material and Methods* and Additional file 1: file 1: section S2-S3).

**Figure 2.**
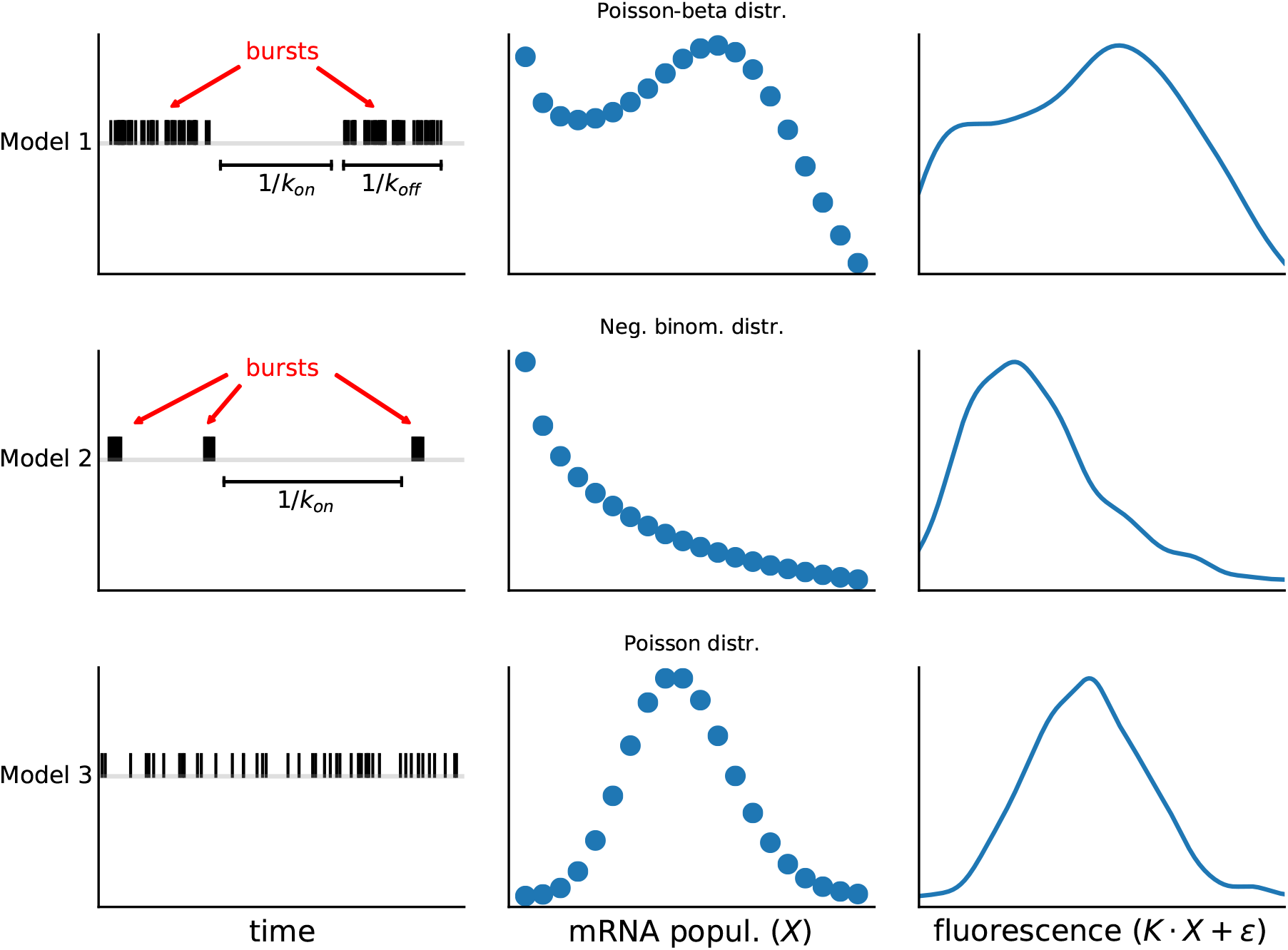
Relations between transcriptional mechanisms, mRNA distributions, model fitting, and parameter estimates. The timing of transcription events at a gene is important. If a population of isogenic cells is induced to express a certain gene, then the resulting mRNA numbers in each cell will reflect the dynamic properties of the process of transcription. While higher frequencies of transcription events unsurprisingly will boost mRNA numbers, more complex patterns can be identified; if transcription occurs clustered in time, forming “bursts” of mRNA production, then the variability of mRNA numbers among individual cells will increase. The precise nature of these relations depends on the mRNA half-life and other dynamical parameters. In fact, these parameters in general shape the distribution of mRNAs among cells in characteristic ways. We can exploit this by testing which parameter values and models are compatible with experimentally derived mRNA distributions and thus infer the dynamics of the underlying transcription process. In combination with experimental perturbations, this produces mechanistic insights into transcription. We applied this approach to our datasets and test their agreement with three models of nested complexity, in line with Occam’s razor. The most complex model corresponds to the transcriptional bursting described above and predicts intricate mRNA distributions, subject to several parameters. Estimating *k*_off_ and *k*_on_ for instance allows us to determine the average times (as their inverse) the genes spend transcribing and non-transcribing, respectively (Model 1). The second model restricts the duration of the bursts and has fewer parameters (Model 2), while the third assumes that the transcriptional events are homogeneous over time (Model 3). These models generate mRNA counts *X* and, in turn, fluorescence intensity, which also depends on the scaling factor *κ* and the measurement noise *E*. To determine which models and parameters best explains our data, we used a Bayesian approach. Broadly speaking, this makes use of the so-called “Bayes’ theorem” to determine the probability of a hypothesis conditional to experimental data. The power of this approach is that it allows the construction of very complex settings from conditional and prior probabilities, which can be computationally explored by means of Markov chain Monte Carlo (MCMC) sampling and produce results which again are probabilities. In general, prior probabilities refer to general assumptions that are taken into account independently of the experimental data, while posterior probabilities result as informed output of the Bayesian inference procedure. The latter correspond to probability distributions of the model parameters, which thus permit excellent assessment of the uncertainties associated with the results. For our study, Bayesian inference was ideal; it allowed us to embed in a single probabilistic framework the data for multiple independent replicates, the measurement precision of our calibration experiments, and the data transformation introduced by the flow cytometer. At the same time, it produced posterior distributions that are highly informative.

Our strategy requires flexible models to represent the absolute mRNA abundance. We considered three stochastic models of gene expression to capture the phenomenology of the transcription process (Figure 2 and *Materials and Methods*). According to the first model, the gene can stay in an “on” state, in which transcription occurs at rate 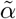, or in an “off” state, in which no transcription occurs. The gene switches from “off” to “on” and “on” to “off” at rates 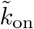 and 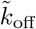, respectively. Assuming that the mRNA degrades at constant rate 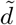, this model corresponds to a Poisson-beta mixture distribution for the stationary per-cell mRNA population, which can be expressed in terms of the dimensionless rates *α*, *k*_on_, and *k*_off_ (Additional file 1: section S2) [4, 49]. The second model is a simplified version of the former two-state model, where *α* and *k*_off_ approach infinity, whilst the ratio *α/k*_off_, which is referred to as the average burst size [50] and incorporated as a single parameter, is held finite; this model gives rise to a negative binomial stationary mRNA distribution and allows much more efficient MCMC sampling than the Poisson-beta model (Additional file 1: sections S3-S4). The third model is the most naïve as it assumes that transcription events of individual mRNAs occur independently at constant rate 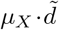, where *μ_X_* is the mean mRNA population, thus yielding a Poisson distributed mRNA population at equilibrium which is thought to characterise genes with unregulated expression [5]. Noise levels consistent with the Poisson model [51, 52] or higher [4, 13] have both been reported in the literature. Estimates of the degradation rates 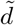 for both mutant and WT transgenes are listed in Additional file 1: section S5.

We obtained better fits for the Poisson-beta and the negative-binomial models than the Poisson model (Additional file 1: sections S4 and S6) for all the replicates. In the Poisson-beta case, the MCMC traces of the rates *k*_off_ and *α* had a strong correlation; this revealed that most of the information about these two parameters is encoded in the ratio *α/k*_off_ (Additional file 1: section S6 and Fig. S10), which is more straightforwardly inferred by means of the negative-binomial model. In fact, for our data, these two models give consistent results in terms of CV^2^, average burst size *α/k*_off_, and burst frequency 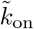. To study the transcriptional noise, we obtained the CV^2^ of the mRNA abundance (which we refer to as 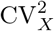 from the estimated parameters (Additional file 1: sections S2 and S6), and plotted it against the estimated mean expression levels *μ_X_* (Figure 3 A-C). These reveal a trend observed before in other systems [6, 53, 54, 55], i.e., the transcriptional noise decreases as *μ*_X_ increases, with the data of each experiment well fitted by a curve of the form 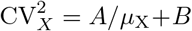, and seems to approach a lower limit beyond which it does not further decrease. Such a limit is known as the noise floor [56, 57, 58, 59, 60]. Strikingly, the presence of the mutation alters the noise trends, thus suggesting that PolII recycling indeed contributes to the noise. The transcriptional noise at intermediate expression levels is significantly higher in WT than mutant cells. For the HBB gene, this pattern extends throughout the range of all induction levels. *Env* shows less pronounced differences between WT and mutant cells for the highest expression levels but resembles HBB otherwise. In all these cases, the noise clearly appears higher than postulated by the Poisson prediction curve 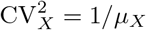 (solid lines in Figure 3 (Additional A-C).

**Figure 3.**
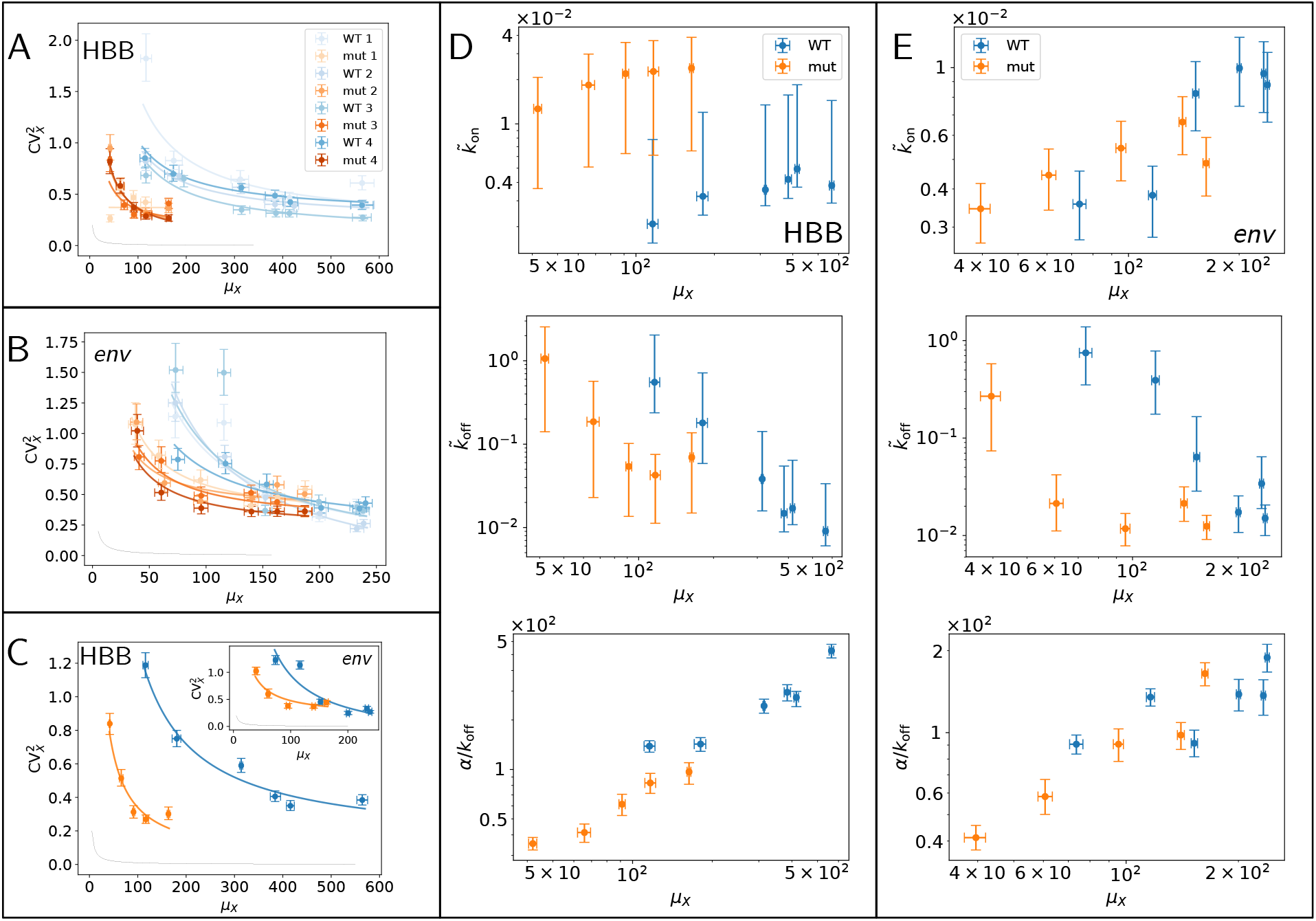
Bayesian parameter estimates. Noise plots of HBB (A) and HIV (B) gene expressions, obtained from the Poisson-beta model for both WT (blue) and mutant (orange) gene variants. Different color intensities correspond to replicates. Mutation changes the balance between noise and average expression level. (C) Results from replicates are aggregated into consensus estimates (Additional file 1: section S4) for HBB and HIV (inset). Solid lines are orthogonal-distance regression curves 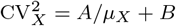. (D-E) Consensus estimates of Poisson-beta model parameters *μ_X_*, 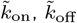, and *α/k*_off_ for HBB (D) and HIV (E). WT (blue) and mutant (orange) show different patterns, with WT genes having highest average burst size and lower burst frequency than mutant at intermediate expression levels. Single-replicate estimates, and negative binomial and Poisson model results are in Additional file 1: section S6. Points and error bars correspond to medians and 90% HPD CIs of the posterior distributions.

Using the DNA content and the forward scatter signal (FSC-A) as proxies of the cell-cycle progression and the cell size, respectively, we heuristically selected populations corresponding to G1, S, and G2 phases of three different sizes each from 40 ng mL^−1^ Tet-induced cells (Figure 4 A-C); we fitted the negative-binomial model to their mRNA-expression reads, and estimated kinetic parameters and noise for each population, separately. Based on this, we found that the cell cycle and size, which typically are major extrinsic-noise contributors, only account for less than 20% of total mRNA variability for the transgenes (Figure 4 D-F), in contrast with [9, 10]; for further details see Additional file 1: section S7.

**Figure 4.**
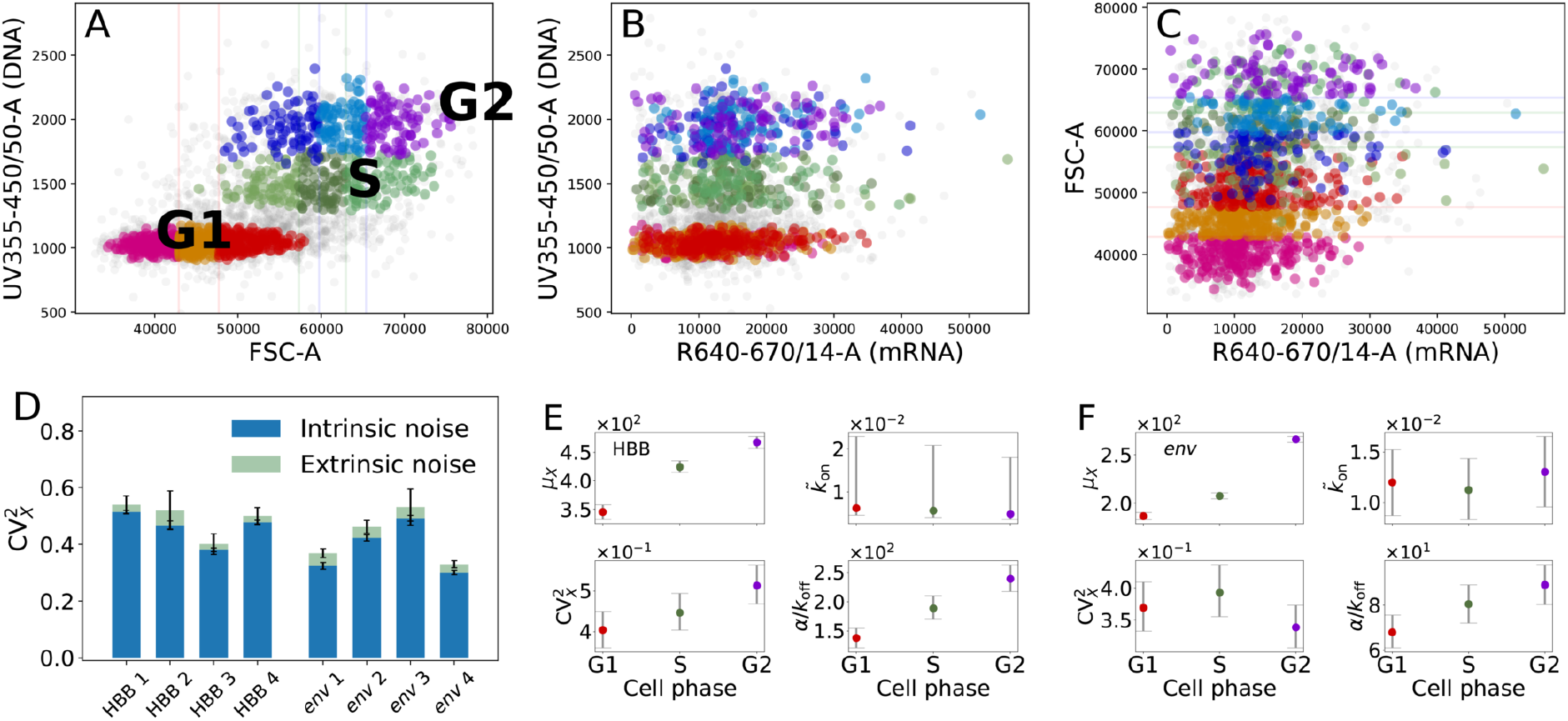
Extrinsic and intrinsic noise. (A), (B), and (C) Scatter plots from flow-FISH signals for the HBB gene, replicate *k* = 3; cells from G1, S, and G2 phase highlighted with red-, green-, and blue-scale colors, respectively; each cell-phase cluster is split into three subsets of different average size with breakpoints at 0.33th and 0.99th quantiles of their FSC-A signals; cell-phase and size are extrinsic variables. (D) Extrinsic and intrinsic contributions to WT HBB and *env* genes’ expression noise, SE error bars obtained via bootstrap. (E) and (F) Cell cycle analysis; consensus estimates of the negative-binomial model parameters for the same genes; points are medians, error bars comprise 90% HPD CIs.

### Modulation of rates

The overall rate estimates obtained from our fits are largely in agreement with previous findings from similar systems [3]. In fact, estimated values of 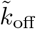 ranged up to ≈2.5 events per minute, with 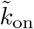 roughly an order of magnitude lower. Increasing the Tet concentration boosts transcription by increasing the average burst size and the frequency 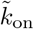 (Figure 3 D), thus shortening the average “off” state duration 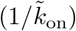. Intriguingly, for the HBB gene, 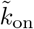 is higher in mutant than WT cells in all cases, while the average burst size is lower in mutant cells in all cases. These patterns are less definite for the *env* gene but appear to support the conclusions from the HBB gene (Figure 3 E and Additional file 1: section S6). In other words, the 3’-5’ crosstalk imposes a constraint on the transcriptional dynamics whose removal can cause bursts to be more frequent and smaller than in the WT gene.

### PolII-mediated 3’-5’ interactions by ChIA-PET

To jointly study the expression of a gene and its 3’-5’ interactions we analysed publicly available datasets for the human cell line K562, obtained from chromatin-interaction analysis by paired-end tag sequencing (ChIA-PET) [61] and single-cell RNA-seq data (scRNAseq) [52]. We chose to use ChIA-PET against PolII to target chromatin interactions that are involved in transcription. We generated HiC-style interaction matrices (whose entries correspond to 2-Kb regions) from the ChIA-PET data using CHIA-PET2 [62]. We filtered the list of genes from the RefGene database with the hg19 reference genome to only contain those with unique gene symbols on chromosomes 1-22 and X, thus excluding alternatively spliced genes. As a proxy of the 3’-5’ interaction of a gene, we first aggregated the reads corresponding to the interaction between the bins that include its transcription start site (TSS) and transcription end site (TES). The resulting metrics depend on the gene length, which we addressed by dividing the number of reads for each gene by the average read number from 10^4^ genomic intervals of the same length as the gene, randomly sampled across the chromosome. We then applied the arcsinh 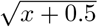 transformation to obtain a variance-stable interaction score [63]. Note that 5’ to 3’ interaction scores correlate with those for 5’ to gene body interactions; this appears unsurprising, given that spatial proximity at one location will favour interaction signals at neighbouring regions, and is tangential to our analyses. We also discarded genes that are shorter than the resolution of our interaction matrices.

Fitting a negative binomial distribution to the scRNA-seq UMI counts data of [52] allows us to conveniently classify expressed genes (sample UMI mean > 0.05) based on the estimated noise 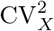, the burst frequency *k*_on_, and the average burst size *α/k*_off_ (*Materials and Methods*, see also [64, 65, 7]). These are plotted against the mean expression *μ_X_* in Figure 5 A-C. It is worth noting that burst frequency averaged over all the genes 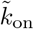, seems to determine the average trends of 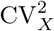 and *α/k*_off_. The noise trend appears to be explained by the curve 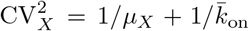 (derived under the negative-binomial assumption, see Additional file 1: section S2), which in fact separates the genes whose noise levels are higher than the mean predicts (blue and orange markers in Figure 5) from those whose noise is lower than the prediction (yellow markers). As a measure of the deviation from this prediction, for each gene, we calculated the vertical distance *ν* of its expression noise to the curve 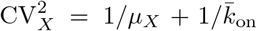 in logarithmic scale, further separating noisy genes for which *ν* > *ν*_1_ (blue makers in Figure 5) from those for which 0 < ν < *ν*_1_ (orange makers). The interaction score of the high-noise genes is significantly higher than the score of the intermediate group, which in turn is higher than the low-noise genes’ (Mann–Whitney U-test, P< 2.2 · 10^16^).

**Figure 5.**
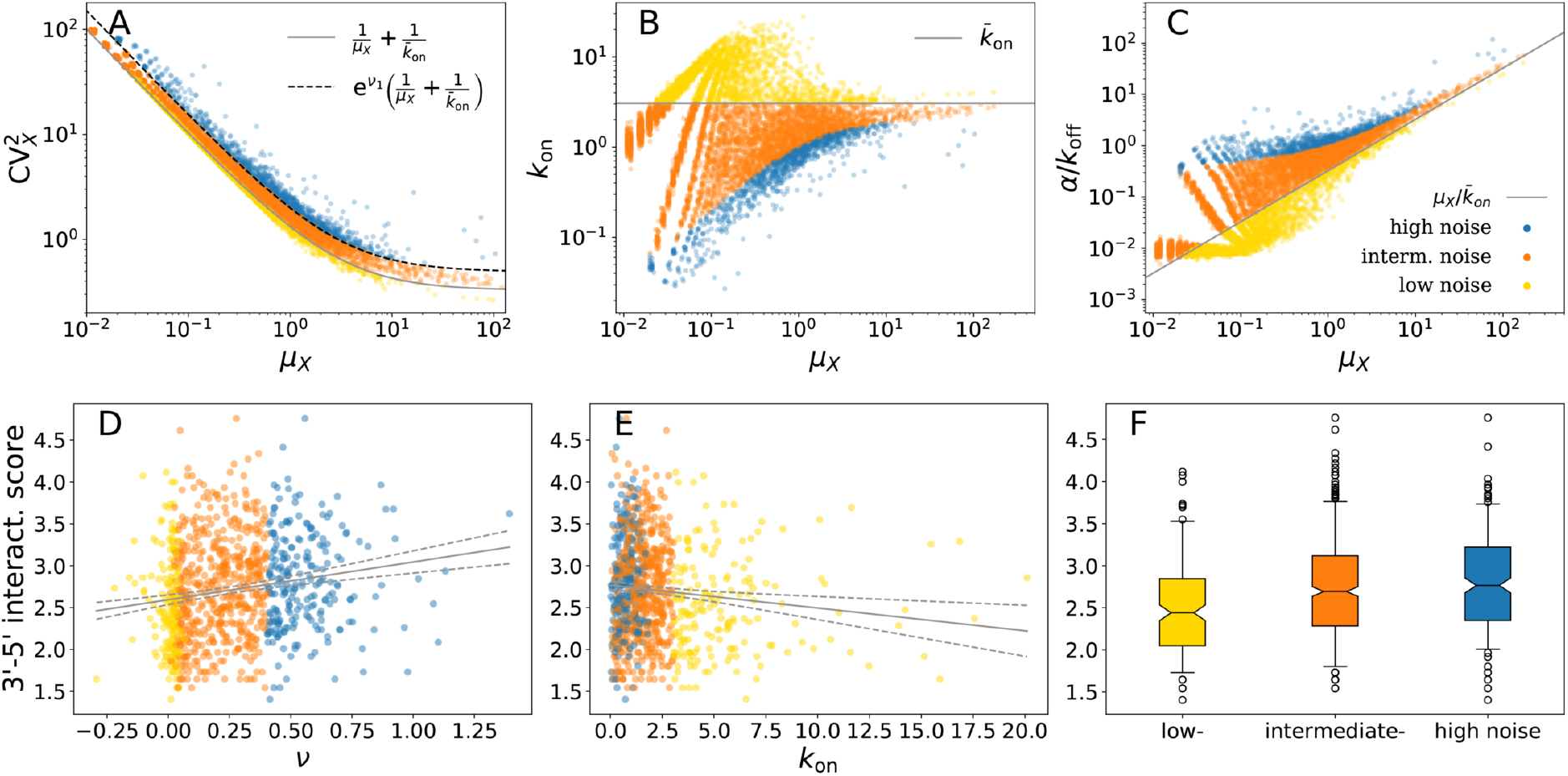
Genome-wide estimates of transcription kinetics and 3’-5’ interactions. (A)-(C) Scatter points correspond to genes, axes are medians of posterior distributions for expression parameters *μ_X_* and 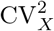, *k*_on_ and *α/k*_off_, respectively, obtained by Bayesian model fitting. Solid lines correspond to the predictions obtained by assuming that all genes have burst frequency equal to the sample average 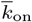. Genes are divided into three groups corresponding to low-, intermediate-, and high-noise levels (yellow, orange, and blue markers, respectively). Dashed line is obtained by setting *ν*_1_ = 4.5 (equation inset in (A)) to separate intermediate- and high-noise genes. (D)-(E) 3’-5’ interaction scores against expression noise (measured as distance *ν* from the solid-line prediction of and burst frequency *k*on; (F) Partitioning the genes by *ν* shows that the interaction score is significantly higher in higher-noise genes than in lower-noise genes (Mann-Whitney U-test, P< 2.2 *·* 10^−16^).

There is a significant positive correlation between the distance *ν* and the interaction score (P< 2.2 · 10^−16^, lm), thus showing that the noise level of genes with high interaction score is typically higher than the mean predicts; we also observe a significant negative correlation between the interaction score and the burst frequency *k*_on_ (P< 2.2 · 10^−16^, lm) and a significant positive correlation between the interaction score and the burst size (P< 2.2 · 10^−16^, lm), consistent with the results on the transgenes. Filtering out zero-count genes, for which there is little statistical information, increases the P-values above to 2.0 · 10^−7^, 2.0 · 10^−3^, and 1.68 · 10^−5^, respectively, due to smaller sample sizes, and yields the scatter plots of Figure 5 D-E and the boxplot of Figure 5 F for the three groups. These results agree with those obtained from different ChIA-PET biological repeats and different bin resolutions (1 Kb and 7 Kb; Additional file 1: section S8 and Fig. S16).

### Microscopic model

To shed further light on the biological mechanisms involved and test whether PolII shuttling can a priori alter the transcriptional noise as seen in the previous section, we constructed and simulated a more complex stochastic model that captures the most important features of our expression system, i.e., induction, polymerase flux between the LLPS droplet (or, more generically, a cluster of PolII [20]) and the gene, transcription, and decay, whilst stripping away non-essential details (Figure 6 A). Its precise formulation, along with additional details, is illustrated in *Materials and Methods* and Additional file 1: section S9. The model is designed around the idea that each PolII waits in a compartment until the transcription occurs [22], where the compartment represents an LLPS droplet (Figure 6 A). This is immersed in its nuclear environment, which adds and removes PolIIs at rates *γ* and *δ*, respectively. In addition to this, by transcribing at rate *β*, the PolIIs leave the compartment with probability 1 − *l* or are re-injected otherwise. This latter reaction represents the crosstalk between the 3’-end processing and the transcription initiation and helps to sustain the compartment population despite the presence of initiation, which on average contributes to depleting it. Consistently with the two genes integrated in our cell lines, the model encodes a Tet-repressor binding site downstream of the TSS which binds to the TetR factor, present at concentration *n*. Such a binding event interrupts the transcription, therefore tuning *n* allows us to control the blocking rate *λ*_off_. The model parameters *l* and *n* are akin to the pA mutation and the Tet concentration, respectively, in the experimental settings. We assume that the pA mutation hinders but does not completely block PolII flux back to the compartment (which can also be facilitated by diffusion, see for instance [16, 24]), therefore the parameter *l* is assumed to be small but still strictly positive even in the presence of pA mutation. During a TetR blockade, PolIIs cannot transcribe and accumulate in the compartment. When the blockade is released, the transcription occurs at a rate directly proportional to the available PolII (consistently with the law of mass action and experimental observations [60, 22]); therefore, at the end of the TetR blockade the compartment is highly populated and the transcription occurs repeatedly while the PolII population quickly drops. As the simulation results demonstrate, the model is able to reproduce an increase of transcriptional bursting upon increasing the recycling probability *l* (Figure 6).

**Figure 6.**
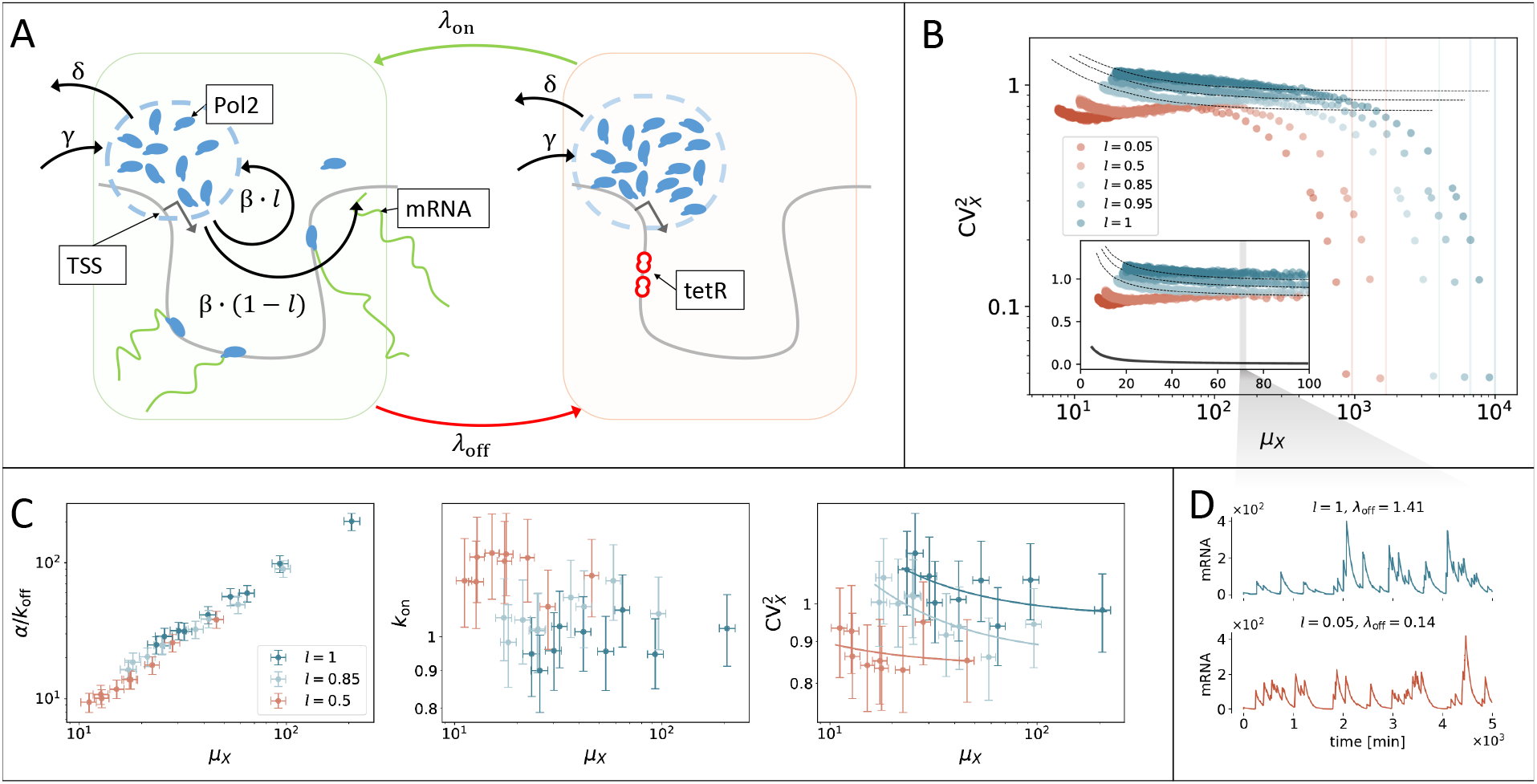
Microscopic model of transcription in Tet-inducible genes. (A) PolIIs (blue) are stored in a compartment (dashed circle) in the proximity of the TSS. With rate *β*, each PolII leaves the compartment to transcribe mRNA and is re-injected with probability *l*. When TetR (tetracycline repressor) binds to the TetO_2_ operator downstream of the TSS (this occurs at rate *λ*_off_), transcription is interrupted and PolIIs accumulate in the compartment. At rate *λ*on, TetR unbinds, thus releasing the large amount of PolIIs accumulated in the compartment to cause bursts, which can be phenomenologically described in terms of the rates 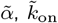, and 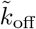. The compartment also exchanges PolIIs with the nuclear environment (at rates *δ* and *γ*). The transcription rate is directly proportional to the abundance of PolIIs, which fluctuates in time and in turn elicits transcriptional noise. Similarly to our experimental system, here we can simulate different Tet concentrations and the recycling probability by tuning the “off”-switch rate *λ*_off_ and *l*, respectively. (B) Noise plots of simulated mRNA abundances. Setting *λ*_*off*_ = *nK*_*λ*_ and *λ*_on_ = *K_λ_*, we imitate the effect of different TetR concentration values by tuning *n*. As Tet presence prevents TetR-TetO_2_ binding, small values of *n* correspond to high Tet-induction levels. For extremely small values of *n*, the gene can be thought of as being always in “on” state, CV^2^ becomes very low, and the limiting value of *μ_X_* can be analytically obtained (vertical lines, see also *SI Appendix*, section S8). *n* ranges from 0.1 to 100, values of the other parameters are (*γ, β, d, δ, K_λ_*) = (10, 10, 0.01, 1, 0.01). Inset: Same scatter plot, axes in linear scale. At intermediate expression levels, 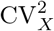 always increases with *l*. Dashed lines are orthogonal-distance regression curves 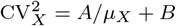, solid line is Poisson-noise curve 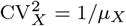. (C) Negative-binomial model fit to 500 mRNA abundances simulated from the microscopic model with *λ*_off_ = 0.5, 1, 1.5, 2, 2.5, 3, 2.5, 3.5, 4, 4.5, values of other parameters as in B. (D) Simulated mRNA-population traces; the two parameter combinations yield almost identical average expressions (sample means 71.3 *±* 0.7 and 70.4 *±* 0.6 over 10^4^ realisations, respectively, SEs obtained via bootstrap), but different biological noise (sample CV^2^s 0.78 *±* 0.01 and 1.07 *±* 0.02, respectively).

This behaviour is conserved under a broad range of different parameter settings, demonstrating that this is a generic result of our model. Fitting a negative binomial distribution with vague prior distributions to an ensemble of mRNA abundances, simulated from this microscopic model, shows patterns consistent with those obtained from the experimental data (Figure 6 C and Additional file 1: section S9).

While actual transcriptional mechanisms are more complex than our idealised model, the latter provides a significant step towards a mechanistic explanation of our observations. In fact, it captures the essential features of the two gene constructs, and naturally reproduces the observed pattern by tuning only the shuttling probability *l* and the factor abundance *n*. Notably, our results demonstrate a minor role for extrinsic contributions to noise (Figure 6 B); in fact, intrinsic factors suffice to yield the noise floor for a wide range of *λ*_off_ and *μ_X_*, which contrasts with several other studies [57, 56, 58, 59, 60].

### Alternative model settings

The pA-site mutations in HBB and *env* transgenes cause termination defects which in turn affect the mRNA degradation rate (Additional file 1: section S5, and [33]). To establish whether the observed noise patterns are ascribable to this, we considered both single-cell expression data and numerical simulations. We analysed human genes in the publicly available dataset of [66], which includes scRNA-seq UMI count data from both influenza-infected and uninfected human A594 cells. Influenza infection causes termination defects in human genes, where transcription can continue for tens of kilobases after the pA site [67, 68]. Native elongation transcript sequencing (NET-seq) also shows that infected cells do not have a difference in initiation of transcription [68]. As suggested in [66], we assumed that a cell is infected if it has at least 0.02% of transcripts coming from influenza genes after 6 hours from virus inoculation; otherwise it is assumed to be uninfected. We then computed the mean expression levels *μ*_*X*,inf_. and *μ*_*X*,uninf_. and the noise levels *ν*_inf_. and *ν*_uninf_. for all human genes (where the subscripts “inf.” and “uninf.” indicate infected and uninfected conditions, respectively). The presence of the termination defect increases the transcript degradation rate, which lowers the UMI counts; we found indeed that *μ*_*X*,inf_. < *μ*_*X*,uninf._ for the overwhelming majority of genes. We also found an overall increase in noise with infection, i.e., *ν*_inf._ > *ν*_uninf._ for many genes, as illustrated in Figure 7 A). A similar scenario is obtained simulating our model with increasing values of mRNA degradation rate *d* and with recycling rate *l* held fixed (Figure 7 B): increasing *d* lowers the average amount of in-silico mRNA and increments its CV^2^.

**Figure 7.**
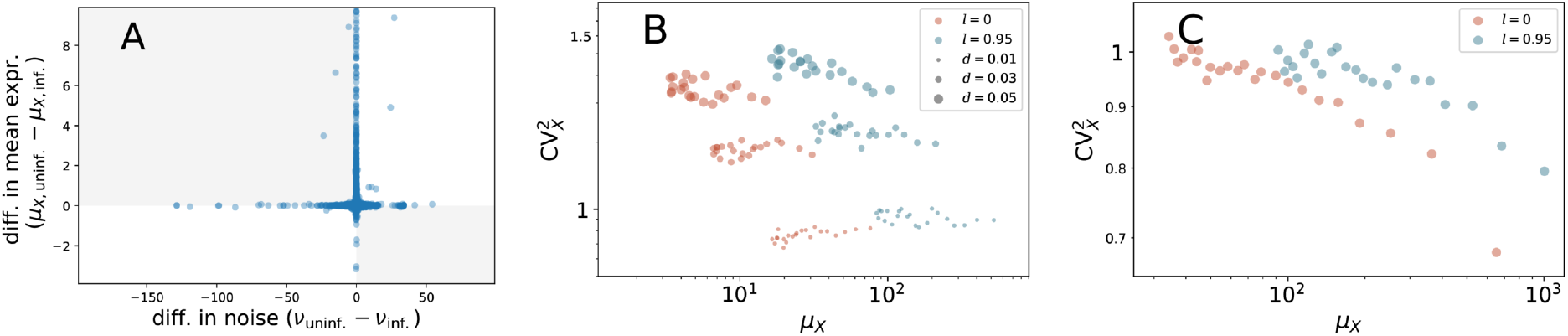
Alternative model settings. (A) Upon influenza infection, termination is altered genome wide, thus affecting observed mRNA counts; *μ*_*X*, uninf_. (*μ_X,_*_inf._) and *ν*_uninf_. (*ν*_inf._) are the mean expression and noise levels in uninfected (infected) cells (measured as the distance from solid-line prediction as in Figure 5), respectively, computed from the scRNA-seq data of [66]. The majority of genes (58%) reports an increase in noise and independently a decrease in mean expression upon infection (upper-left quadrant in scatter plot is the most populated). (B) Noise plots of mRNA abundances simulated according to the model of Figure 6 with same parameter values except *l* and *d* as in legend; incrementing the mRNA degradation rate *d* suppresses *μ_X_* for both values of *l*, whilst increasing CV^2^. (C) Noise plots of mRNA abundances in a variant of the looping model; recycling is allowed but PolIIs cannot pile up before initiation, with noise being virtually unaffected when *l* is tuned from 0.95 to 0; other model parameters are as in Figure 6(B) except (*γ, β*) = (50, 100). The settings of (B) and (C) cannot explain the noise patterns observed in HBB and *env* transgenes.

This scenario does not fit the experimental transgenes observations, where pA mutation equally lowers *μ_X_* but decreases 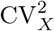, and therefore it is not a plausible representation of their true biological mechanisms.

Further, we considered a variant of our model where PolIIs are not allowed to condensate in a compartment before the transcription begins. The importance of particle condensation to fluctuations in the presence of an on-off switch has been mathematically described [69]. In the modified in-silico model, indeed, increasing the recycling rate does not increase the noise (Figure 7 C), thus suggesting that a reservoir of PolIIs may be a crucial component of gene regulation.

## Discussion

The wealth of existing results strongly suggests the occurrence of 3’-5’ crosstalk in the WT variants of our transgene systems, involving physical interaction between factors at either gene end and recycling of polymerases, which can be disrupted or strongly reduced upon a point mutation. Similarly, information of the interactions between the ends of genes involved in transcription can be accessed genome-wide by means of PolII ChIA-PET sequencing.

Based on both, an in-depth analyses of the transgene systems (which provide a controlled experimental setting) and an observational study of ChIA-PET sequencing data (which provide a genome-wide view of chromatin interactions involved in transcription), we present results to suggest that PolII-mediated 3’-5’ interactions are major contributors to transcriptional noise.

Building on standard phenomenological models, transcription parameters, such as average burst size and frequency, are consistently inferred across the different conditions using a Bayesian methodology, to demonstrate the presence of association between 3’-5’ interactions and transcription kinetics. Modelling transcription requires abstraction and simplification due to the complexity of the molecular processes involved and the inadequacy of current experimental methodologies to dynamically resolve structural interactions at individual loci. Furthermore, the Bayesian estimates of the kinetic parameters reflect the incomplete quantitative information available on the experimental device. Also note that our transgenes might not exactly represent the average endogenous gene. Nevertheless, our setting is sufficient to resolve specific patterns, which can be reproduced by an *ab-initio* mechanistic model, thus supporting our conclusions.

The analysis suggests that recycling of the polymerase typically increases noise at a given expression level, while an alternative symmetric interpretation is possible, viz., that recycling permits higher expression at a given noise level. These relations are either a byproduct of the construction of the transcriptional machinery or were selected for. It will be interesting to further explore our findings from an evolutionary perspective. In particular, many studies show how selection of noisy expression can be critical by contributing to cell fate diversity [70, 71] and by favouring their long-term survival in adverse environments [72]. This could also have implications in synthetic biology, where the optimisation of gene expression and the control of its noise are desirable features [73, 74]. Our work provides an important contribution to the field of systems biology by identifying a single base, and thus a genetic determinant, that modulates the balance between the average expression level and its variation.

## Materials and Methods

### Measurement equation and Monte Carlo estimation

We assume that the measured fluorescence *Y_i_* of cell *i* is proportional to the true mRNA abundance *X_i_* and therefore can be expressed as 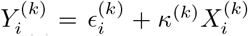 where (*k*) indexes the replicate, *κ*^(*k*)^ can be thought of as a scale, and 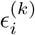 is the zero of such a scale, also corresponding to the background of unspecific staining and auto-fluorescence of the *i*th cell [48]. The background noise is measured, for each replicate *k*, by means of control cells whose gene of interest has been deleted. These are used to define informative priors for 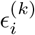. Our choice is 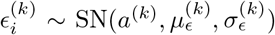, i.e., the control-cell fluorescence *y* is supposed to have Azzalini’s skew-normal distribution

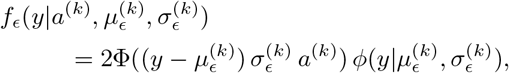

where Φ and *φ* are the standard normal CDF and normal PDF, respectively, while the mean 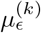, the standard deviation 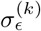, and the skewness parameter *a*^(*k*)^ are point estimates from the control data sets. Prior distributions for *κ*^(*k*)^ are chosen based on the regression coefficients of gamma generalised linear model fits with identity link. For the remaining parameters we assume vague gamma priors with mean 1 and variance 10^3^. Adaptive Metropolis–Hastings samplers for model fitting were implemented (Additional file 1: section S4).

### Phenomenological two-state gene-expression models

The transcriptional bursting is fully characterised by the rates 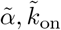 and 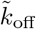 in units of min^−1^. It is convenient to express the rates in units of the inverse of the mean mRNA life-time 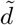, i.e., 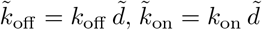 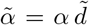. It can be shown that the stationary mRNA abundance *X* for this model is Poisson beta with probability density function (PDF)

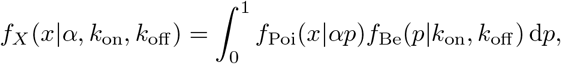

where *f*_Poi_(*x*|*α*) = *α^x^*e^−α^/*x*! and 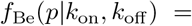 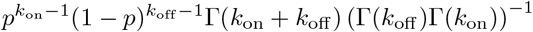 are PDFs of Poisson and beta random variables (RVs), respectively. This expresses the hierarchy

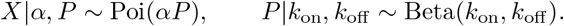

It is convenient to reparametrise the Poisson-beta PDF in terms of its mean *μ_X_* = *αk*_on_*/*(*k*_off_ + *k*_on_), to get

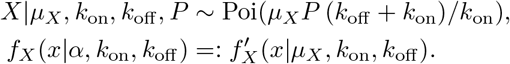

In fact, this allows us to exploit knowledge on *μ_X_* in the form of informative priors and infer the dimensionless rates *α*, *k*_off_ and *k*_on_. These are converted to min^−1^ by using 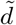 estimated from data (Additional file 1: section S5). In the limit as *k*_off_ → ∞, *α* → ∞, with their ratio *α/k*_off_ held finite, the population mean satisfies *μ_X_* = *k*_on_*α/k*_off_, while the PDF of *X* approaches the negative binomial distribution

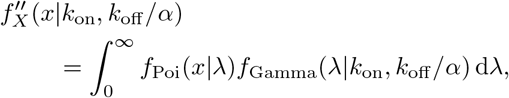

where *f*_Gamma_(*x*|*k*_on_*, k*_off_ */α*) is the density of a Gamma RV with mean *μ_X_* and variance *μ_X_k*_off_ */α*; when this RV concentrates near the mean as *k*_on_ → ∞ and *k*_off_ */α* → 0, *X* is Poisson with PDF *f*_Poi_(*x*|*μ_X_*).

### Microscopic model

The microscopic model is defined by means of the following chemical reaction scheme:

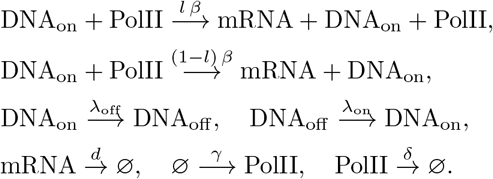

By the law of mass action, *λ*_off_ = *nK_λ_*, *λ*_on_ = *K_λ_*, where *K_λ_* and *n* represent the chemical affinity and concentration of TetR homodimers that bind to the TetO_2_ operators downstream of the TSS, respectively. When such a binding event occurs, the transcription is inhibited as elongation is impeded and the resulting locked DNA configuration is represented by the species DNA_off_. The switch to DNA_on_ corresponds to the release of the lock. A variant of this model that does not allow PolII to accumulate before trascription is obtained with *γ* > 0 when the PolII compartment is empty and *γ* = 0 otherwise.

## Additional Files

Additional file 1 — “Supporting Information to: “3’-5’ crosstalk contributes to transcriptional bursting”

## Data availability

Custom scripts have been made available at https://github.com/mcavallaro/gLoop under GNU-GPLv3.0 licence [88]. Data that support the findings of this study have been deposited in Zenodo [89] and in the National Center for Biotechnology Information Gene Expression Omnibus with accession number GSE124682 [90].

## Acknowledgements

We thank Louise Dyson, Matt Moores, Lucy Ternent, and Jonathan Keith for valuable discussions, and Sharon Collier and Charlotte Petersen for minor contributions.

## Ethics approval and consent to participate

Not applicable.

## Authors’ contributions

M.C., S.T., B.F., and D.H. designed research; M.C., M.D.W., M.J., J.T., and D.H. performed research; M.C., M.D.W., M.J., and D.H. analyzed data; M.C., M.D.W., and D.H. wrote the paper with input from all authors. All authors read and approved the final manuscript.

## References

1. Golding, I., Paulsson, J., Zawilski, S.M., Cox, E.C.: Real-time kinetics of gene activity in individual bacteria. Cell 123(6), 1025–36 (2005). doi:10.1016/j.cell.2005.09.031

2. Chubb, J.R., Trcek, T., Shenoy, S.M., Singer, R.H.: Transcriptional pulsing of a developmental gene. Current Biology 16(10), 1018–1025 (2006). doi:10.1016/J.CUB.2006.03.092

3. Suter, D.M., Molina, N., Gatfield, D., Schneider, K., Schibler, U., Naef, F.: Mammalian genes are transcribed with widely different bursting kinetics. Science 332(6028), 472–474 (2011). doi:10.1126/science.1198817

4. Raj, A., Peskin, C.S., Tranchina, D., Vargas, D.Y., Tyagi, S.: Stochastic mRNA synthesis in mammalian cells. PLoS Biology 4(10), 309 (2006). doi:10.1371/journal.pbio.0040309

5. Munsky, B., Neuert, G., van Oudenaarden, A.: Using Gene Expression Noise to Understand Gene Regulation. Science 336(6078), 183–187 (2012). doi:10.1126/science.1216379

6. Singh, A., Razooky, B., Cox, C.D., Simpson, M.L., Weinberger, L.S.: Transcriptional bursting from the HIV-1 promoter is a significant source of stochastic noise in hiv-1 gene expression. Biophysical Journal 98(8), 32–34 (2010). doi:10.1016/j.bpj.2010.03.001

7. Larsson, A.J.M., Johnsson, P., Hagemann-Jensen, M., Hartmanis, L., Faridani, O.R., Reinius, B., Segerstolpe, Å., Rivera, C.M., Ren, B., Sandberg, R.: Genomic encoding of transcriptional burst kinetics. Nature 565(7738), 251–254 (2019). doi:10.1038/s41586-018-0836-1

8. Elowitz, M.B., Levine, A.J., Siggia, E.D., Swain, P.S.: Stochastic gene expression in a single cell. Science 297(5584), 1183–1186 (2002). doi:10.1126/science.1070919

9. Zopf, C.J., Quinn, K., Zeidman, J., Maheshri, N.: Cell-cycle dependence of transcription dominates noise in gene expression. PLoS Computational Biology 9(7), 1003161 (2013). doi:10.1371/journal.pcbi.1003161

10. Sherman, M.S., Lorenz, K., Lanier, M.H., Cohen, B.A.: Cell-to-cell variability in the propensity to transcribe explains correlated fluctuations in gene expression. Cell Systems 1(5), 315–325 (2015). doi:10.1016/j.cels.2015.10.011

11. Skinner, S.O., Xu, H., Nagarkar-Jaiswal, S., Freire, P.R., Zwaka, T.P., Golding, I.: Single-cell analysis of transcription kinetics across the cell cycle. eLife 5(2016). doi:10.7554/eLife.12175

12. Padovan-Merhar, O., Nair, G., Biaesch, A., Mayer, A., Scarfone, S., Foley, S., Wu, A., Churchman, L.S., Singh, A., Raj, A.: Single mammalian cells compensate for differences in cellular volume and DNA copy number through independent global transcriptional mechanisms. Molecular Cell 58(2), 339–352 (2015). doi:10.1016/J.MOLCEL.2015.03.005

13. Battich, N., Stoeger, T., Pelkmans, L.: Control of transcript variability in single mammalian cells. Cell 163(7), 1596–610 (2015). doi:10.1016/j.cell.2015.11.018

14. Raser, J.M., O’Shea, E.K.: Control of stochasticity in eukaryotic gene expression. Science 304(5678), 1811–4 (2004). doi:10.1126/science.1098641

15. Weinberger, L., Voichek, Y., Tirosh, I., Hornung, G., Amit, I., Barkai, N.: Expression Noise and Acetylation Profiles Distinguish HDAC Functions. Molecular Cell 47(2), 193–202 (2012). doi:10.1016/J.MOLCEL.2012.05.008

16. van Zon, J.S., Morelli, M.J., Tanase-Nicola, S., ten Wolde, P.R.: Diffusion of transcription factors can drastically enhance the noise in gene expression. Biophysical Journal 91(12), 4350–4367 (2006). doi:10.1529/BIOPHYSJ.106.086157

17. Chong, S., Chen, C., Ge, H., Xie, X.S.: Mechanism of transcriptional bursting in bacteria. Cell 158(2), 314–326 (2014). doi:10.1016/j.cell.2014.05.038

18. Fukaya, T., Lim, B., Levine, M.: Enhancer control of transcriptional bursting. Cell 166(2), 358–368 (2016). doi:10.1016/j.cell.2016.05.025

19. Bartman, C., Hsu, S., Hsiung, C.-S., Raj, A., Blobel, G.: Enhancer regulation of transcriptional bursting parameters revealed by forced chromatin looping. Molecular Cell 62(2), 237–247 (2016). doi:10.1016/J.MOLCEL.2016.03.007

20. Papantonis, A., Cook, P.R.: Transcription factories: Genome organization and gene regulation. Chemical Reviews 113(11), 8683–8705 (2013). doi:10.1021/cr300513p

21. Cisse, I.I., Izeddin, I., Causse, S.Z., Boudarene, L., Senecal, A., Muresan, L., Dugast-Darzacq, C., Hajj, B., Dahan, M., Darzacq, X.: Real-time dynamics of RNA polymerase II clustering in live human cells. Science 341(6146), 664–667 (2013). doi:10.1126/science.1239053

22. Cho, W.K., Jayanth, N., English, B.P., Inoue, T., Andrews, J.O., Conway, W., Grimm, J.B., Spille, J.H., Lavis, L.D., Lionnet, T., Cisse, I.I.: RNA Polymerase II cluster dynamics predict mRNA output in living cells. eLife 5(MAY2016) (2016). doi:10.7554/eLife.13617

23. Plys, A.J., Kingston, R.E.: Dynamic condensates activate transcription. Science 361(6400), 329–330 (2018). doi:10.1126/science.aau4795

24. Boehning, M., Dugast-Darzacq, C., Rankovic, M., Hansen, A.S., Yu, T., Marie-Nelly, H., McSwiggen, D.T., Kokic, G., Dailey, G.M., Cramer, P., Darzacq, X., Zweckstetter, M.: RNA polymerase II clustering through carboxy-terminal domain phase separation. Nature Structural and Molecular Biology 25(9), 833–840 (2018). doi:10.1038/s41594-018-0112-y

25. Chong, S., Dugast-Darzacq, C., Liu, Z., Dong, P., Dailey, G.M., Cattoglio, C., Heckert, A., Banala, S., Lavis, L., Darzacq, X., Tjian, R.: Imaging dynamic and selective low-complexity domain interactions that control gene transcription. Science 361(6400), 2555 (2018). doi:10.1126/science.aar2555

26. Sabari, B.R., Dall’Agnese, A., Boija, A., Klein, I.A., Coffey, E.L., Shrinivas, K., Abraham, B.J., Hannett, N.M., Zamudio, A.V., Manteiga, J.C., Li, C.H., Guo, Y.E., Day, D.S., Schuijers, J., Vasile, E., Malik, S., Hnisz, D., Lee, T.I., Cisse, I.I., Roeder, R.G., Sharp, P.A., Chakraborty, A.K., Young, R.A.: Coactivator condensation at super-enhancers links phase separation and gene control. Science 361(6400), 3958 (2018). doi:10.1126/science.aap9195

27. Cho, W.K., Spille, J.H., Hecht, M., Lee, C., Li, C., Grube, V., Cisse, I.I.: Mediator and RNA polymerase II clusters associate in transcription-dependent condensates. Science 361(6400), 412–415 (2018). doi:10.1126/science.aar4199

28. Cramer, P.: Organization and regulation of gene transcription. Nature (2019). doi:10.1038/s41586-019-1517-4

29. Shin, Y., Chang, Y.C., Lee, D.S.W., Berry, J., Sanders, D.W., Ronceray, P., Wingreen, N.S., Haataja, M., Brangwynne, C.P.: Liquid Nuclear Condensates Mechanically Sense and Restructure the Genome. Cell 175(6), 1481–149113 (2018). doi:10.1016/j.cell.2018.10.057

30. Wei, M.T., Chang, Y.C., Shimobayashi, S.F., Shin, Y., Strom, A.R., Brangwynne, C.P.: Nucleated transcriptional condensates amplify gene expression. Nature Cell Biology (2020). doi:10.1038/s41556-020-00578-6

31. Moore, M.J., Proudfoot, N.J.: Pre-mRNA Processing Reaches Back toTranscription and Ahead to Translation. Cell 136(4), 688–700 (2009). doi:10.1016/J.CELL.2009.02.001

32. Kuehner, J.N., Pearson, E.L., Moore, C.: Unravelling the means to an end: RNA polymerase II transcription termination. Nature Reviews Molecular Cell Biology 12(5), 283–294 (2011). doi:10.1038/nrm3098

33. Mapendano, C.K., Lykke-Andersen, S., Kjems, J., Bertrand, E., Jensen, T.H.: Crosstalk between mRNA 3’ End Processing and Transcription Initiation. Molecular Cell 40(3), 410–422 (2010). doi:10.1016/j.molcel.2010.10.012

34. Fang, X., Wang, L., Ishikawa, R., Li, Y., Fiedler, M., Liu, F., Calder, G., Rowan, B., Weigel, D., Li, P., Dean, C.: Arabidopsis FLL2 promotes liquid–liquid phase separation of polyadenylation complexes. Nature 569(7755), 265–269 (2019). doi:10.1038/s41586-019-1165-8

35. Hebenstreit, D.: Are gene loops the cause of transcriptional noise? Trends in Genetics 29(6), 333–338 (2013)

36. Damgaard, C.K., Kahns, S., Lykke-Andersen, S., Nielsen, A.L., Jensen, T.H., Kjems, J.: A 5’ splice site enhances the recruitment of basal transcription initiation factors in vivo. Molecular Cell 29(2), 271–278 (2008). doi:10.1016/J.MOLCEL.2007.11.035

37. Carter, D., Chakalova, L., Osborne, C.S., Dai, Y.-f., Fraser, P.: Long-range chromatin regulatory interactions in vivo. Nature Genetics 32(4), 623–626 (2002). doi:10.1038/ng1051

38. Reik, A., Telling, A., Zitnik, G., Cimbora, D., Epner, E., Groudine, M.: The locus control region is necessary for gene expression in the human beta-globin locus but not the maintenance of an open chromatin structure in erythroid cells. Molecular and cellular biology 18(10), 5992–6000 (1998)

39. Tolhuis, B., Palstra, R.J., Splinter, E., Grosveld, F., de Laat, W.: Looping and interaction between hypersensitive sites in the active beta-globin locus. Molecular cell 10(6), 1453–65 (2002)

40. Hnisz, D., Shrinivas, K., Young, R.A., Chakraborty, A.K., Sharp, P.A.: A Phase Separation Model for Transcriptional Control. Cell 169(1), 13–23 (2017). doi:10.1016/j.cell.2017.02.007

41. Wang, Y., Fairley, J.A., Roberts, S.G.E.: Phosphorylation of TFIIB links transcription initiation and termination. Current biology : CB 20(6), 548–53 (2010). doi:10.1016/j.cub.2010.01.052

42. Ansari, A., Hampsey, M.: A role for the CPF 3’-end processing machinery in RNAP II-dependent gene looping. Genes and Development 19(24), 2969–2978 (2005). doi:10.1101/gad.1362305

43. Wani, S., Yuda, M., Fujiwara, Y., Yamamoto, M., Harada, F., Ohkuma, Y., Hirose, Y.: Vertebrate Ssu72 regulates and coordinates 3’-end formation of RNAs transcribed by RNA polymerase II. PLoS ONE 9(8), 106040 (2014). doi:10.1371/journal.pone.0106040

44. Tan-ong, S.M., Zaugg, J.B., Camblong, J., Xu, Z., Zhang, D.W., Mischo, H.E., Ansari, A.Z., Luscombe, N.M., Steinmetz, L.M., Proudfoot, N.J.: Gene loops enhance transcriptional directionality. Science 338(6107), 671–675 (2012). doi:10.1126/science.1224350

45. Perkins, K.J., Lusic, M., Mitar, I., Giacca, M., Proudfoot, N.J.: Transcription-dependent gene looping of the HIV-1 provirus is dictated by recognition of pre-mRNA processing signals. Molecular Cell 29(1), 56–68 (2008). doi:10.1016/j.molcel.2007.11.030

46. Shapiro, H.M.: Practical Flow Cytometry. John Wiley & Sons, Inc., Hoboken, NJ, USA (2003). doi:10.1002/0471722731. http://doi.wiley.com/10.1002/0471722731

47. Lo, K., Brinkman, R.R., Gottardo, R.: Automated gating of flow cytometry data via robust model-based clustering. Cytometry Part A 73A(4), 321–332 (2008). doi:10.1002/cyto.a.20531

48. Tiberi, S., Walsh, M., Cavallaro, M., Hebenstreit, D., Finkenstädt, B.: Bayesian inference on stochastic gene transcription from flow cytometry data. Bioinformatics 34(17), 647–655 (2018). doi:10.1093/bioinformatics/bty568

49. Kim, J., Marioni, J.C.: Inferring the kinetics of stochastic gene expression from single-cell RNA-sequencing data. Genome Biology 14(1), 7 (2013). doi:10.1186/gb-2013-14-1-r7

50. Dobrzynski, M., Bruggeman, F.J.: Elongation dynamics shape bursty transcription and translation. Proceedings of the National Academy of Sciences 106(8), 2583–2588 (2009). doi:10.1073/pnas.0803507106

51. Zenklusen, D., Larson, D.R., Singer, R.H.: Single-RNA counting reveals alternative modes of gene expression in yeast. Nature Structural and Molecular Biology 15(12), 1263–1271 (2008). doi:10.1038/nsmb.1514

52. Klein, A.M., Mazutis, L., Akartuna, I., Tallapragada, N., Veres, A., Li, V., Peshkin, L., Weitz, D.A., Kirschner, M.W.: Droplet barcoding for single-cell transcriptomics applied to embryonic stem cells. Cell 161(5), 1187–1201 (2015). doi:10.1016/j.cell.2015.04.044

53. Dar, R.D., Shaffer, S.M., Singh, A., Razooky, B.S., Simpson, M.L., Raj, A., Weinberger, L.S.: Transcriptional bursting explains the noise-versus-mean relationship in mRNA and protein levels. PLoS ONE 11(7), 0158298 (2016). doi:10.1371/journal.pone.0158298

54. Dar, R.D., Razooky, B.S., Weinberger, L.S., Cox, C.D., Simpson, M.L.: The low noise limit in gene expression. PLoS ONE 10(10), 0140969 (2015). doi:10.1371/journal.pone.0140969

55. Soltani, M., Vargas-Garcia, C.A., Antunes, D., Singh, A.: Intercellular Variability in Protein Levels from Stochastic Expression and Noisy Cell Cycle Processes. PLoS Computational Biology 12(8), 1004972 (2016). doi:10.1371/journal.pcbi.1004972

56. Bar-Even, A., Paulsson, J., Maheshri, N., Carmi, M., O’Shea, E., Pilpel, Y., Barkai, N.: Noise in protein expression scales with natural protein abundance. Nature Genetics 38(6), 636–643 (2006). doi:10.1038/ng1807

57. Newman, J.R.S., Ghaemmaghami, S., Ihmels, J., Breslow, D.K., Noble, M., DeRisi, J.L., Weissman, J.S.: Single-cell proteomic analysis of S. cerevisiae reveals the architecture of biological noise. Nature 441(7095), 840–846 (2006). doi:10.1038/nature04785

58. Taniguchi, Y., Choi, P.J., Li, G.-W., Chen, H., Babu, M., Hearn, J., Emili, A., Xie, X.S.: Quantifying E. coli proteome and transcriptome with single-molecule sensitivity in single cells. Science 329(5991), 533–8 (2010). doi:10.1126/science.1188308

59. Stewart-Ornstein, J., Weissman, J.S., El-Samad, H.: Cellular noise regulons underlie fluctuations in Saccharomyces cerevisiae. Molecular cell 45(4), 483–93 (2012). doi:10.1016/j.molcel.2011.11.035

60. Yang, S., Kim, S., Rim Lim, Y., Kim, C., An, H.J., Kim, J.-H., Sung, J., Lee, N.K.: Contribution of RNA polymerase concentration variation to protein expression noise. Nature Communications 5(1), 4761 (2014). doi:10.1038/ncomms5761

61. Li, G., Ruan, X., Auerbach, R.K., Sandhu, K.S., Zheng, M., Wang, P., Poh, H.M., Goh, Y., Lim, J., Zhang, J., Sim, H.S., Peh, S.Q., Mulawadi, F.H., Ong, C.T., Orlov, Y.L., Hong, S., Zhang, Z., Landt, S., Raha, D., Euskirchen, G., Wei, C.-L., Ge, W., Wang, H., Davis, C., Fisher-Aylor, K.I., Mortazavi, A., Gerstein, M., Gingeras, T., Wold, B., Sun, Y., Fullwood, M.J., Cheung, E., Liu, E., Sung, W.-K., Snyder, M., Ruan, Y.: Extensive promoter-centered chromatin interactions provide a topological basis for transcription regulation. Cell 148(1-2), 84–98 (2012). doi:10.1016/j.cell.2011.12.014

62. Li, G., Chen, Y., Snyder, M.P., Zhang, M.Q.: ChIA-PET2: a versatile and flexible pipeline for ChIA-PET data analysis. Nucleic Acids Research 45(1), 4–4 (2017). doi:10.1093/nar/gkw809

63. Bartlett, M.S.: The Use of Transformations. Biometrics 3(1), 39–52 (1947). doi:10.2307/3001536

64. Svensson, V.: Droplet scRNA-seq is not zero-inflated. Nature Biotechnology 38(2), 147–150 (2020). doi:10.1038/s41587-019-0379-5

65. Townes, F.W., Hicks, S.C., Aryee, M.J., Irizarry, R.A.: Feature selection and dimension reduction for single-cell RNA-Seq based on a multinomial model. Genome Biology 20(1), 295 (2019). doi:10.1186/s13059-019-1861-6

66. Russell, A.B., Trapnell, C., Bloom, J.D.: Extreme heterogeneity of influenza virus infection in single cells. eLife (2018). doi:10.7554/eLife.32303

67. Zhao, N., Sebastiano, V., Moshkina, N., Mena, N., Hultquist, J., Jimenez-Morales, D., Ma, Y., Rialdi, A., Albrecht, R., Fenouil, R., Sánchez-Aparicio, M.T., Ayllon, J., Ravisankar, S., Haddad, B., Ho, J.S.Y., Low, D., Jin, J., Yurchenko, V., Prinjha, R.K., Tarakhovsky, A., Squatrito, M., Pinto, D., Allette, K., Byun, M., Smith, M.L., Sebra, R., Guccione, E., Tumpey, T., Krogan, N., Greenbaum, B., van Bakel, H., García-Sastre, A., Marazzi, I.: Influenza virus infection causes global RNAPII termination defects. Nature Structural and Molecular Biology (2018). doi:10.1038/s41594-018-0124-7

68. Bauer, D.L.V., Tellier, M., Martínez-Alonso, M., Nojima, T., Proudfoot, N.J., Murphy, S., Fodor, E.: Influenza Virus Mounts a Two-Pronged Attack on Host RNA Polymerase II Transcription. Cell Reports (2018). doi:10.1016/j.celrep.2018.04.047

69. Cavallaro, M., Mondragón, R.J., Harris, R.J.: Temporally correlated zero-range process with open boundaries: Steady state and fluctuations. Phys. Rev. E 92(2), 022137 (2015). doi:10.1103/PhysRevE.92.022137

70. Kussell, E., Leibler, S.: Phenotypic diversity, population growth, and information in fluctuating environments. Science 309(5743), 2075–8 (2005). doi:10.1126/science.1114383

71. Eldar, A., Elowitz, M.B.: Functional roles for noise in genetic circuits. Nature 467(7312), 167–173 (2010). doi:10.1038/nature09326

72. Beaumont, H.J.E., Gallie, J., Kost, C., Ferguson, G.C., Rainey, P.B.: Experimental evolution of bet hedging. Nature 462(7269), 90–93 (2009). doi:10.1038/nature08504

73. Murphy, K.F., Adams, R.M., Wang, X., Balázsi, G., Collins, J.J.: Tuning and controlling gene expression noise in synthetic gene networks. Nucleic Acids Research 38(8), 2712–2726 (2010). doi:10.1093/nar/gkq091

74. Bandiera, L., Furini, S., Giordano, E.: Phenotypic variability in synthetic biology applications: Dealing with noise in microbial gene expression. Frontiers in Microbiology 7, 479 (2016). doi:10.3389/fmicb.2016.00479

75. Patil, A., Huard, D., Fonnesbeck, C.: PyMC : Bayesian Stochastic Modelling in Python. Journal of Statistical Software 35(4), 1–81 (2010). doi:10.18637/jss.v035.i04

76. Hahne, F., LeMeur, N., Brinkman, R.R., Ellis, B., Haaland, P., Sarkar, D., Spidlen, J., Strain, E., Gentleman, R.: flowCore: a Bioconductor package for high throughput flow cytometry. BMC Bioinformatics 10, 106 (2009)

77. Lo, K., Hahne, F., Brinkman, R.R., Gottardo, R.: flowClust: a Bioconductor package for automated gating of flow cytometry data. BMC Bioinformatics 10(1), 145 (2009). doi:10.1186/1471-2105-10-145

78. Spiess, A.: qpcR: Modelling and Analysis of Real-Time PCR Data. https://CRAN.R-project.org/package=qpcR (2013)

79. Martin, M.: Cutadapt removes adapter sequences from high-throughput sequencing reads. EMBnet.journal 17(1), 10 (2011). doi:10.14806/ej.17.1.200

80. Quinlan, A.R., Hall, I.M.: BEDTools: a flexible suite of utilities for comparing genomic features. Bioinformatics 26(6), 841–2 (2010). doi:10.1093/bioinformatics/btq033

81. Dobin, A., Davis, C.A., Schlesinger, F., Drenkow, J., Zaleski, C., Jha, S., Batut, P., Chaisson, M., Gingeras, T.R.: STAR: ultrafast universal RNA-seq aligner. Bioinformatics 29(1), 15–21 (2013). doi:10.1093/bioinformatics/bts635

82. Dyer, N.P., Shahrezaei, V., Hebenstreit, D.: LiBiNorm: an htseq-count analogue with improved normalisation of Smart-seq2 data and library preparation diagnostics. PeerJ 7, 6222 (2019). doi:10.7717/peerj.6222

83. Anders, S., Pyl, P.T., Huber, W.: HTSeq–a Python framework to work with high-throughput sequencing data. Bioinformatics 31(2), 166–169 (2015). doi:10.1093/bioinformatics/btu638

84. Ramírez, F., Dündar, F., Diehl, S., Grüning, B.A., Manke, T.: deepTools: a flexible platform for exploring deep-sequencing data. Nucleic Acids Research 42(Web Server issue), 187–91 (2014). doi:10.1093/nar/gku365

85. Love, M.I., Huber, W., Anders, S.: Moderated estimation of fold change and dispersion for RNA-seq data with DESeq2. Genome Biology 15(12), 550 (2014). doi:10.1186/s13059-014-0550-8

86. Goldstein, L.D., Cao, Y., Pau, G., Lawrence, M., Wu, T.D., Seshagiri, S., Gentleman, R.: Prediction and Quantification of Splice Events from RNA-Seq Data. PLOS ONE 11(5), 0156132 (2016). doi:10.1371/journal.pone.0156132

87. Liao, Y., Smyth, G.K., Shi, W.: The R package Rsubread is easier, faster, cheaper and better for alignment and quantification of RNA sequencing reads. Nucleic Acids Research 47(8), 47–47 (2019). doi:10.1093/nar/gkz114

88. Cavallaro, M., Walsh, M.D., Jones, M., Teahan, J., Tiberi, S., Finkenstädt, B., Hebenstreit, D.: Supporting software to “3’-5’ crosstalk contributes to transcriptional bursting”. https://github.com/mcavallaro/gLoop. GitHub (2020). doi:10.5281/zenodo.4127058

89. Cavallaro, M., Walsh, M.D., Jones, M., Teahan, J., Tiberi, S., Finkenstädt, B., Hebenstreit, D.: Supporting data to “3’-5’ crosstalk contributes to transcriptional bursting”. Zenodo (2020). doi:10.5281/zenodo.4304833

90. Cavallaro, M., Walsh, M.D., Jones, M., Teahan, J., Tiberi, S., Finkenstädt, B., Hebenstreit, D.: Supporting RNA-seq data to “3’-5’ crosstalk contributes to transcriptional bursting”. https://www.ncbi.nlm.nih.gov/geo/query/acc.cgi?acc=GSE124682. Gene Expression Omnibus (2020)

