## Additional file 1 for "3’-5’ crosstalk contributes to transcriptional bursting"

### CONTENTS

S1. Description of data 1
  A. Flow-cytometry data 1
  B. Control cells 2
  C. SmFISH and Nanostring barcoding data 2

S2. Phenomelogical gene expression models 6

S3. Measurement and technical error model 7

S4. Monte Carlo estimation 9
  A. MCMC samplers 9
  B. Consensus posteriors 10
  C. Goodness of fit 11

S5. mRNA decay rates 11

S6. Summary of MCMC results 13
  A. Poisson-beta distribution 13
  B. Negative binomial distribution 20
  C. Poisson distribution 27

S7. Cell cycle 29

S8. PolII-mediated 3'-5' interactions by ChIA-PET 29

S9. Microscopic gene expression model 31

S10. Materials 32
  A. Cell lines and cell culture 32
  B. Single-molecule RNA fluorescence in situ hybridization 32
  C. RNA isolation and preparation, and degradation rate estimation 33
  D. Nanostring 33
  E. RNA-seq 34

References 36

### S1. DESCRIPTION OF DATA

#### A. Flow-cytometry data

We obtained flow cytometry data (from the BD LSRFortessa$^{TM}$ cell analyzer and BD FACSDiva$^{TM}$ software) for cell lines expressing the genes *env* and HBB, in both their wild-type (WT) and mutated (mut) versions. For each gene, experimental data were collected in four replicates (8 in total), each containing groups of observations corresponding to cells stimulated with tetracycline (Tet) at concentrations of $5, 10, 20, 40, 80$, and $250$ ng/mL, respectively.

Each data-set was stored in a `.fcs` format file and it was imported and pre-processed in R as an object of class `flowFrame`, which consists of an annotated data-frame class defined in the flowCore R package [1] and designed to deal with flow-cytometry data. Rows in such data frames correspond to single measurements. Each row contains the values of two fluorescence intensities that correspond to staining for mRNA and total DNA and are labeled by `R640-670/14-A` and `UV355-450/50-A`, respectively. These readings were compesated for spectral overlap with flowCore. In addition to this, the values of four scattering observations, namely `FSC.H`, `FSC.W`, `SSC.H`, and `SSC.W`, were recorded. Such observations are thought to be correlated to cell size and granularity. Values for each observation are stored in so-called "arbitrary units" (a.u.) [2].

The first task is to identify records in the data sets corresponding to either cell debris or clumps of cells, which have to be removed from subsequent analysis. We apply the robust model-based clustering approach of Ref. [3], distributed as the flowClust package [4], to identify cell populations in the data. Based on the scattering observations, the points were grouped into 3 clusters, and the set corresponding to single cells is the one with intermediate size and granularity, as suggested by the

DNA content (see Fig. S1). For the data sets where the three detected clusters overlap, points where grouped into two clusters, instead. For this second case, inspection shows that the cluster with lower size and granularity corresponds to single cells. Standard rectangle gates were applied to remove a few outlier points whose `UV355-450/50-A` reads were lower than 500. Kernel density estimates (KDE) for the populations after this sub-setting are plotted in Fig. 2 (Main Text) and Fig. S2. Technical variation affects the shapes of the distributions only for some HIV replicates, with shoulders at the lower ends sometimes merging with the main mode. The cells corresponding to the data points in the shoulders exhibit normal characteristics (in terms of cell size and DNA content) and thus probably reflect cells without mRNA. The parameter estimates, reported in section S6, appear robust with regards to the absence or presence of the shoulders save for the highest $\mu_X$s.

#### B. Control cells

For each replicate $k$, we consider control cells, where the gene of interest has been deleted (see *Main text*). Such control cells were subjected to the same staining procedure as the others, which leaves a background of fluorescence probes that are not specifically bound to the mRNA. We argue that such background fluorescence stain are also present in the cells expressing the transgenes and contribute a term $\epsilon_i^{(k)}$ to the signal detected by the cell-analyzer channel of label `R640-670/14-A` for each cell $i$. The histograms of the signal from control cells appear skewed, as illustrated for example in Fig. S3 (left). We chose to fit the Azzalini's skew-normal distribution, that has PDF

$$f_\epsilon(y|a^{(k)}, \mu_\epsilon^{(k)}, \sigma_\epsilon^{(k)}) \\ = 2\Phi((y-\mu_\epsilon^{(k)})\,\sigma_\epsilon^{(k)}\,a^{(k)})\,\phi(y|\mu_\epsilon^{(k)}, \sigma_\epsilon^{(k)}), \quad (1)$$

to such data, where $\Phi$ and $\phi$ are the standard normal CDF and normal PDF, respectively, while the mean $\mu_\epsilon^{(k)}$, the standard deviation $\sigma_\epsilon^{(k)}$, and the skewness parameter $a^{(k)}$ are point estimates from the control data sets. The maximum likelihood estimates for each replicate are reported in Table S1 (see also Figs. S3(left)).

#### C. SmFISH and Nanostring barcoding data

Flow-FISH data are supplemented by microscopy-based single-molecule FISH counts (which we simply refer to as smFISH) and Nanostring nCounter® Technology bar-coding measurements. These assays are used to choose informative priors for the mean mRNA abundance and, in turn, to calibrate the flow-FISH readouts. Symbols $\overline{x}$, $s_x^2$, and $s_{\overline{x}}$ represent sample mean, sample variance, and standard error of the mean, respectively. Based on these, we chose truncated normal informative priors for the average expression level $\mu_X \sim \mathcal{N}(\overline{x}, s_{\overline{x}})$, with the constraint $\mu_X > 0$, for all replicates $k$.

HEK293 cells are not ideal for smFISH, since they tend to overlap when growing, producing dense clusters after dividing. A further problem with smFISH is the limited dynamic range of suitable microscopes. In fact, images tend to be overexposed when recorded during transcriptional bursts at settings that are otherwise optimal for lower transcript numbers and, conversely, optimal settings for transcriptional bursts do not cope well when transcript numbers are low. Therefore we only exploit smFISH results to infer the average expression level, and rely on flow-FISH to study the noise. SmFISH data yield the summary statistics of Table S2 for the mean expression of HBB. Nanostring data (*SI Dataset 1*) yields the summary statistics of Table S3 for the mean expression of *env*.

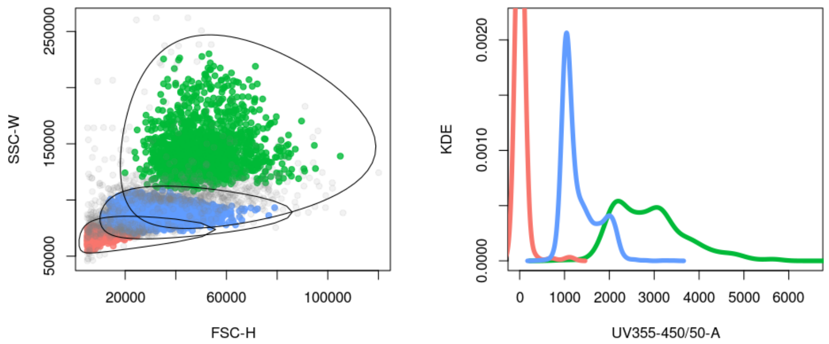


FIG. S1. Clustering of flow-cytometry data. (left) Clusters are projected to the `FSC.W-SSC.H` plane and plotted with the ellipses that delimit the 0.60 quantiles of fitted t-distributions. (right) Inspection of `UV355-450/50-A` shows signature distributions for DNA content, thus suggesting that the central cluster (in blue color) contains single-cell reads. Duplets have twice the amount of DNA content than single cells.

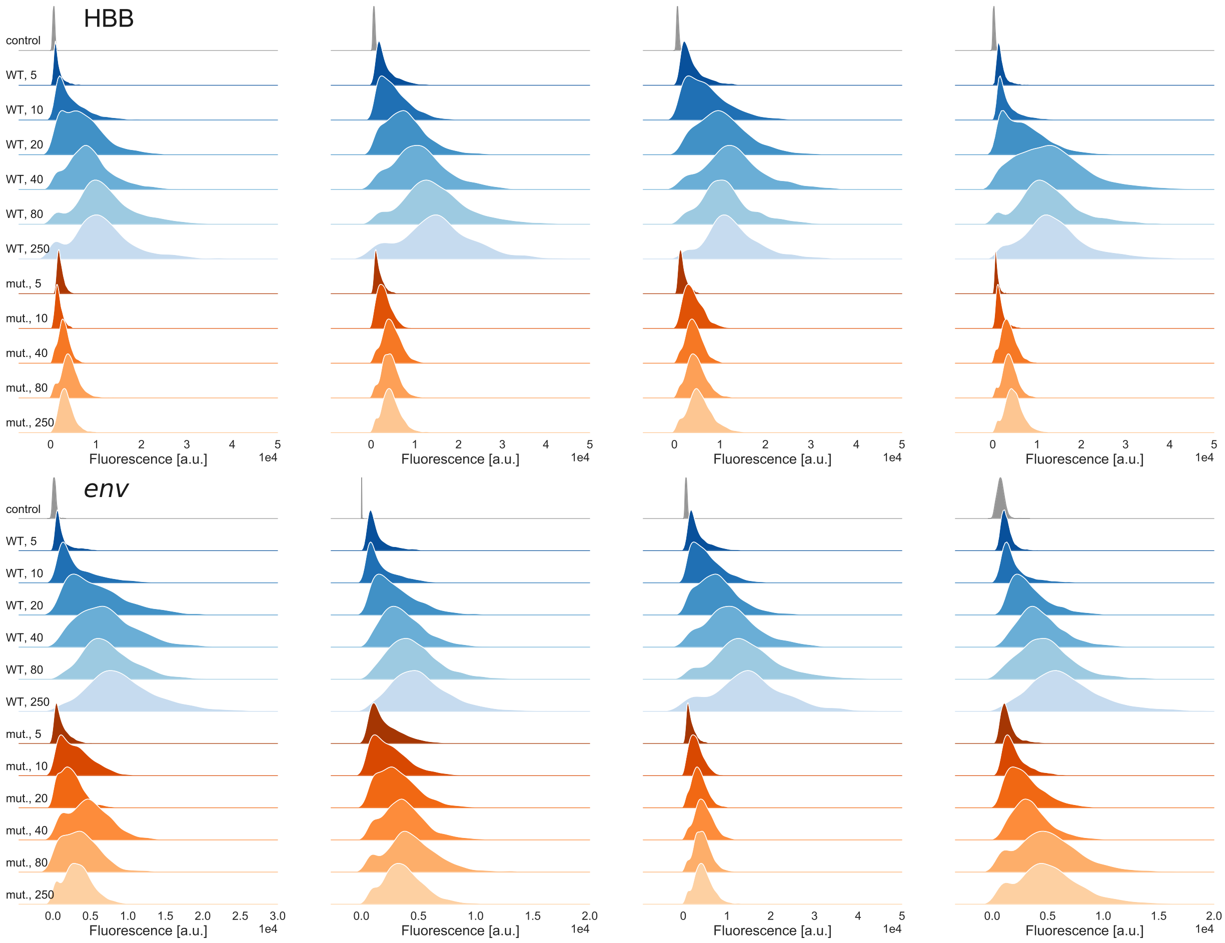


FIG. S2. KDEs of the flow-FISH single-cell readings corresponding to the abundances of HBB (top) and *env* (bottom) transcripts, from wild-type (blue), mutant (orange), and control (gray) cells, from 4 replicates per transgene, $k = 1, 2, 3, 4$ (left to right), at the different induction level (Tet concentrations in unit of ng/mL, shades of colors). Gene expression saturates upon increasing Tet concentration and mutant-cell expressions is lower than the wild-type. Fluorescence is given in arbitrary units (a.u.), $y$-axes are not to scale.

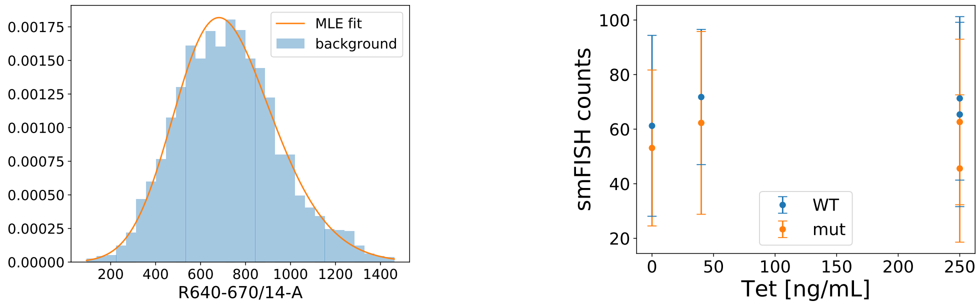


FIG. S3. Control measurements. (left) The MLE skew-normal density (line) of the background data for Tet=40 ng/mL, $k = 1$, WT, is in good agreement with the empirical histogram. (right) Microscopy FISH count summary of housekeeping gene Akt1 vs induction levels, for both wild-type and mutant cells. Points and error bars are sample means and standard deviations, respectively.

TABLE S1. MLE estimation of the control-cell fluorescence (replicates 2 (HBB) and 4 (HIV) have the same background parameters as measurements were performed the same day with the same control cells).

| gene | k | Tet | $\mu^{(k)}$ | $\sigma^{(k)}$ | $a^{(k)}$ | $s_\mu^{(k)}$ | $s_\sigma^{(k)}$ | $s_a^{(k)}$ |
|---|---|---|---|---|---|---|---|---|
| HBB | 1 | 0 | 389.504 | 348.098 | 1.817 | 15.670 | 12.396 | 0.214 |
| HBB | 1 | 250 | 514.828 | 307.460 | 1.632 | 11.667 | 9.027 | 0.153 |
| HBB | 2 | 0 | 625.804 | 458.715 | 1.863 | 21.687 | 17.915 | 0.230 |
| HBB | 2 | 250 | 459.139 | 311.688 | 2.075 | 9.098 | 7.703 | 0.169 |
| HBB | 3 | 0 | 539.613 | 360.195 | 2.046 | 11.838 | 10.270 | 0.182 |
| HBB | 3 | 250 | 493.913 | 337.605 | 2.091 | 10.440 | 9.088 | 0.178 |
| HBB | 4 | 0 | 53.140 | 320.685 | 2.303 | 7.201 | 6.514 | 0.151 |
| HBB | 4 | 250 | 112.667 | 256.183 | 1.748 | 5.507 | 4.497 | 0.094 |
| HIV | 1 | 0 | 565.327 | 390.016 | 1.834 | 8.397 | 6.936 | 0.102 |
| HIV | 1 | 250 | 443.898 | 463.603 | 1.813 | 14.774 | 12.117 | 0.148 |
| HIV | 2 | 0 | -31.395 | 230.794 | 1.108 | 13.129 | 8.356 | 0.145 |
| HIV | 2 | 250 | -23.282 | 312.375 | 2.066 | 8.109 | 7.156 | 0.145 |
| HIV | 3 | 0 | 196.970 | 401.184 | 5.423 | 6.238 | 8.517 | 0.425 |
| HIV | 3 | 250 | 407.813 | 259.586 | 1.445 | 16.865 | 12.125 | 0.226 |
| HIV | 4 | 0 | 625.804 | 458.715 | 1.863 | 21.687 | 17.915 | 0.230 |
| HIV | 4 | 250 | 459.139 | 311.688 | 2.075 | 9.098 | 7.703 | 0.169 |

TABLE S2. Summary statistics for the mean expression of HBB, obtained from microscopy FISH.

| gene | Tet | $\overline{x}$ | $s_x^2$ | sample size |
|---|---|---|---|---|
| WT | 0 | 64.868 | 2984.538 | 585 |
| WT | 5 | 115.450 | 7640.059 | 202 |
| WT | 10 | 175.150 | 18093.547 | 193 |
| WT | 20 | 312.945 | 32327.429 | 347 |
| WT | 40 | 384.111 | 23962.077 | 190 |
| WT | 80 | 414.105 | 31613.582 | 437 |
| WT | 250 | 565.351 | 54765.760 | 342 |
| mut | 0 | 38.953 | 1431.185 | 379 |
| mut | 5 | 41.645 | 717.115 | 279 |
| mut | 10 | 62.995 | 3573.429 | 198 |
| mut | 40 | 90.709 | 3301.445 | 468 |
| mut | 80 | 115.413 | 7096.164 | 179 |
| mut | 250 | 163.547 | 7375.582 | 892 |

TABLE S3. Summary statistics for the mean expression of HIV, obtained from nCounter® data. The standard error of the mean $s_{\overline{x}}^2$ is obtained propagating the errors from the nCounter® and the Atk1 smFISH measurements used for normalisation.

| gene | Tet | $\overline{x}$ | $s_{\overline{x}}^2$ |
|---|---|---|---|
| WT | 0 | 24.723 | 3.247 |
| WT | 5 | 72.975 | 3.247 |
| WT | 10 | 115.872 | 3.247 |
| WT | 20 | 151.462 | 3.247 |
| WT | 40 | 199.433 | 3.247 |
| WT | 80 | 232.644 | 3.247 |
| WT | 250 | 238.178 | 3.247 |
| mut | 0 | 21.875 | 3.247 |
| mut | 5 | 36.842 | 3.247 |
| mut | 10 | 58.108 | 3.247 |
| mut | 20 | 95.460 | 3.247 |
| mut | 40 | 139.979 | 3.247 |
| mut | 80 | 162.874 | 3.247 |
| mut | 250 | 187.287 | 3.247 |

### S2. PHENOMELOGICAL GENE EXPRESSION MODELS

We describe the gene expression in terms of the standard phenomenological two-state model [5]. This model assumes that the gene randomly alternates between an "on" and an "off" state, and that the mRNA is only transcribed, at rate $\tilde{\alpha}$, during the on state. The gene switches from "off" to "on" and from "on" to "off" states after an exponentially distributed random time with mean $1/k_{\text{on}}$ and $1/k_{\text{off}}$, respectively. Consequently, the transcriptional bursting is fully characterised by the rates $\tilde{\alpha}$, $\tilde{k}_{\text{on}}$, and $\tilde{k}_{\text{off}}$. In addition to this, mRNA is degraded at rate $\tilde{d}$. It is convenient to express the rates in units of the inverse of the mean mRNA life-time $\tilde{d}$, i.e.,

$$\tilde{k}_{\text{off}} = k_{\text{off}}\,\tilde{d}, \tag{2}$$

$$\tilde{k}_{\text{on}} = k_{\text{on}}\,\tilde{d}, \tag{3}$$

$$\tilde{\alpha} = \alpha\,\tilde{d}. \tag{4}$$

It can be shown that the stationary probability density function (PDF) of the mRNA population $x$ for this model is (see, e.g., Ref. [6])

$$f_X(x|\alpha, k_{\text{on}}, k_{\text{off}}) = \frac{\alpha^x \mathrm{e}^{-\alpha}\,\Gamma(k_{\text{on}}+x)\Gamma(k_{\text{on}}+k_{\text{off}})}{x!\,\Gamma(k_{\text{on}}+k_{\text{off}}+x)\,\Gamma(k_{\text{on}})}\,{}_1F_1(k_{\text{off}}, k_{\text{on}}+k_{\text{off}}+x;\alpha), \tag{5}$$

where $\Gamma$ is the gamma function and ${}_1F_1$ is the confluent hyper-geometric function of the first kind. An alternative representation of the PDF (5) is

$$f_X(x|\alpha, k_{\text{on}}, k_{\text{off}}) = \int_0^1 f_{\text{Poi}}(x|\alpha p) f_{\text{Be}}(p|k_{\text{on}}, k_{\text{off}})\,\mathrm{d}p, \tag{6}$$

where

$$f_{\text{Poi}}(x|\alpha) = \frac{\alpha^x \mathrm{e}^{-\alpha}}{x!}, \tag{7}$$

$$f_{\text{Be}}(p|k_{\text{on}}, k_{\text{off}}) = p^{k_{\text{on}}-1}(1-p)^{k_{\text{off}}-1}\frac{\Gamma(k_{\text{on}}+k_{\text{off}})}{\Gamma(k_{\text{off}})\Gamma(k_{\text{on}})}, \tag{8}$$

are density distributions of Poisson and beta random variables (RVs), respectively.

The PDF of equation (6) encodes the following hierarchy

$$X|\alpha, P \sim \text{Poi}(\alpha P), \tag{9}$$

$$P|k_{\text{on}}, k_{\text{off}} \sim \text{Beta}(k_{\text{on}}, k_{\text{off}}). \tag{10}$$

Further details can be found, e.g., in Refs. [6, 7]. It is convenient to reparametrise the Poisson-beta distribution in terms of its mean

$$\mu_X = \alpha\frac{k_{\text{on}}}{k_{\text{off}}+k_{\text{on}}}, \tag{11}$$

to get

$$X|\mu_X, k_{\text{on}}, k_{\text{off}}, P \sim \text{Poi}(\mu_X\frac{k_{\text{off}}+k_{\text{on}}}{k_{\text{on}}}P), \tag{12}$$

$$f_X(x|\alpha, k_{\text{on}}, k_{\text{off}}) =: f'_X(x|\mu_X, k_{\text{on}}, k_{\text{off}}). \tag{13}$$

In fact, this allows us to exploit knowledge on $\mu_X$ in the form of informative priors of S1 C. The expression for the squared coefficient of variation ($\text{CV}^2$) can also be written in terms of $\mu_X$, i.e.,

$$\text{CV}_X^2 = \frac{1}{\mu_X} + \frac{k_{\text{off}}}{k_{\text{on}}(1+k_{\text{off}}+k_{\text{on}})}, \tag{14}$$

where the second term on the r.h.s. quantifies the overdispersion of $X$ with respect to a Poisson random variable. Such a functional relation between $\mathrm{CV}_X^2$ and $\mu_X$ has been encountered in gene expression data [8–12]. The probability $P$ cannot be directly accessed and therefore is a latent (hidden) variable for the model. In fact, for our data, the mRNA number is a latent variable too, being only inferred from the measured fluorescence signals. This can be encoded into a measurement equation, as explained in the next section.

In the limit as $k_{\mathrm{off}} \to \infty$, $\alpha \to \infty$, with the ratio $\alpha/k_{\mathrm{off}}$ held fixed, the population mean and $\mathrm{CV}^2$ satisfy

$$\mu_X = \frac{\alpha}{k_{\mathrm{off}}} k_{\mathrm{on}}, \tag{15}$$

$$\mathrm{CV}_X^2 = \frac{1}{\mu_X} + \frac{1}{k_{\mathrm{on}}}, \tag{16}$$

respectively, while the distribution of $X$ approaches the negative binomial distribution with PDF

$$f_X''(x|\mu_X, k_{\mathrm{on}}) = \frac{\Gamma(k_{\mathrm{on}} + x)}{x!\,\Gamma(k_{\mathrm{on}})} \left(\frac{k_{\mathrm{on}}}{k_{\mathrm{on}} + \mu_X}\right)^{k_{\mathrm{on}}} \left(\frac{\mu_X}{k_{\mathrm{on}} + \mu_X}\right)^{x}. \tag{17}$$

This can be easily proven using the Poisson-gamma mixture formulation of the negative binomial RV $X$, i.e.,

$$X|\lambda \sim \mathrm{Poi}(\lambda), \tag{18}$$

$$\lambda|k_{\mathrm{on}}, k_{\mathrm{off}}/\alpha \sim \mathrm{Gamma}(k_{\mathrm{on}}, k_{\mathrm{off}}/\alpha). \tag{19}$$

In fact, the beta distribution scaled by $\alpha > 0$ approaches the gamma distribution as $k_{\mathrm{off}} \to \infty$, $\alpha \to \infty$, i.e.,

$$\frac{1}{\alpha} \frac{\Gamma(k_{\mathrm{on}} + k_{\mathrm{off}})}{\Gamma(k_{\mathrm{on}})\Gamma(k_{\mathrm{off}})} \left(\frac{x}{\alpha}\right)^{k_{\mathrm{on}}-1} \left(1 - \frac{x}{\alpha}\right)^{k_{\mathrm{off}}-1} \asymp \frac{\frac{k_{\mathrm{off}}}{\alpha}^{k_{\mathrm{on}}} x^{k_{\mathrm{on}}-1}\, \mathrm{e}^{-\frac{k_{\mathrm{off}}}{\alpha} x}}{\Gamma(k_{\mathrm{on}})}, \tag{20}$$

which follows from known asymptotic relations

$$\lim_{k_{\mathrm{off}} \to \infty} \frac{\Gamma(k_{\mathrm{off}} + k_{\mathrm{on}})}{\Gamma(k_{\mathrm{off}}) {k_{\mathrm{off}}}^{k_{\mathrm{on}}}} = 1, \tag{21}$$

$$\lim_{\alpha \to \infty} \left(1 - \frac{x}{\alpha}\right)^{\alpha \frac{k_{\mathrm{off}}}{\alpha}} = \mathrm{e}^{-\frac{k_{\mathrm{off}}}{\alpha} x}. \tag{22}$$

The ratio $\alpha/k_{\mathrm{off}}$ has a simple interpretation, being the expected number of transcription events during an on phase. In Ref. [13] this ratio has been referred to as the "expected burst size". In the limit as $k_{\mathrm{on}} \to \infty$ with the mean expression $\mu_X$ held fixed, the negative binomial distribution (17) approaches the distribution

$$f_X'''(x|\mu_X) := f_{\mathrm{Poi}}(x|\mu_X). \tag{23}$$

### S3. MEASUREMENT AND TECHNICAL ERROR MODEL

The DB FACSDiva™ software manual [14] specifies that the light intensity from fluorescent dyes is amplified linearly within a wide range (see also, e.g., Refs. [9, 15]). Based on this, we assume that the measured fluorescence $Y_i$ of cell $i$ is proportional to the true mRNA abundance $X_i$ and therefore can be expressed as in the following "measurement" equation,

$$Y_i^{(k)} = \epsilon_i^{(k)} + \kappa^{(k)} X_i^{(k)}, \tag{24}$$

where $k$ indexes the replicate, $\kappa$ can be thought of as a scale and $\epsilon_i$ is the zero of such a scale, also corresponding to the background of unspecific staining and auto-fluorescence of the $i$th cell [7].

The dispersion of biological data is typically due to both technical errors, caused by the measurement process, and the variability intrinsic to the underlying biology. In our measurement model, for the variables $X_i^{(k)}$ to best accommodate the true biological noise of $Y_i^{(k)}$, it is important that $\epsilon_i^{(k)}$ and $\kappa^{(k)}$ are specified with sufficient precision and accuracy.

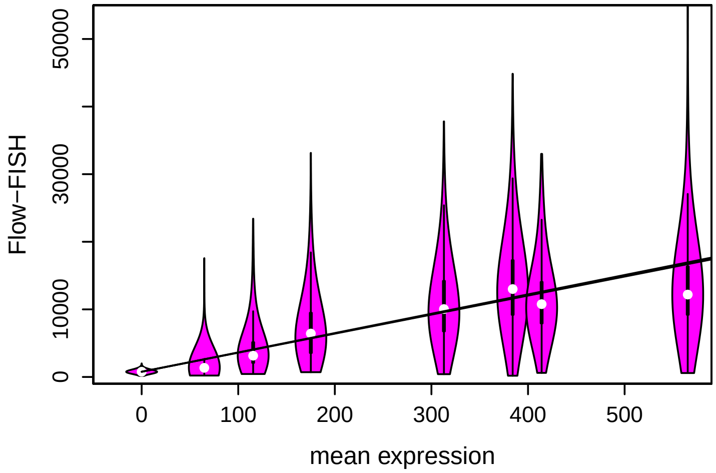


FIG. S4. Flow-FISH data (violin plots [16]) vs mean expression levels obtained from FISH data for the replicate $k = 3$, WT HBB gene. Their relation is captured by a linear model with coefficient $\kappa$.

Informative priors for $\epsilon_i^{(k)}$ are chosen according to section S1 B, i.e.,

$$\epsilon_i^{(k)} \sim \mathrm{SN}(a^{(k)}, \mu_\epsilon^{(k)}, \sigma_\epsilon^{(k)}), \tag{25}$$

where the parameters $a^{(k)}$, $\mu_\epsilon^{(k)}$, and $\sigma_\epsilon^{(k)}$ are estimated from the control cells at 250 ng/mL Tet. The standard errors of the maximum likelihood estimates are neglected. As a consequence, all the single-cell measurements can be thought of as being subjected to the same random background, thus mitigating tractability issues. For a more comprehensive fully-Bayesian hierarchical approach see Ref. [7].

To pin down informative priors for $\kappa^{(k)}$, we perform gamma regression. For each flow-FISH data-set, 500 random cell readings are selected for the main Monte Carlo estimation of section S4. The remaining reads are used as response variables for a gamma regression with identity link. Covariates are mean expression level point estimates from section S1 C. As an example, this is illustrated in Fig. S4 for $k = 3$, WT HBB gene. The GLM estimates of the expected values $\mu'_\kappa$ along with the standard errors $s'_\kappa$ are reported in Table S4. Our prior choice is the truncated normal RV

$$\kappa \sim \mathcal{N}(\mu'_\kappa, s'_\kappa), \tag{26}$$

with the constraint $\kappa > 1$, where the mean $\mu'_\kappa$ and standard deviation $s'_\kappa$ are obtained from the 16 values in Table S4 according to the laws of total expectation and variance, respectively, i.e.,

$$\mu'_\kappa = \overline{\mu_\kappa}, \qquad {s'}_\kappa^2 = \overline{s_\kappa^2} + \overline{\mu_\kappa^2} - \overline{\mu_\kappa}^2, \tag{27}$$

where the bar notation represents averages.

For the remaining parameters we assume

$$k_{\mathrm{on}} \sim \mathrm{Gamma}(\alpha_{k_{\mathrm{on}}}, \beta_{k_{\mathrm{on}}}), \tag{28}$$

$$k_{\mathrm{off}} \sim \mathrm{Gamma}(\alpha_{k_{\mathrm{off}}}, \beta_{k_{\mathrm{off}}}), \tag{29}$$

$$\alpha_{k_{\mathrm{on}}} = \beta_{k_{\mathrm{on}}} = \alpha_{k_{\mathrm{off}}} = \beta_{k_{\mathrm{off}}} = 0.001, \tag{30}$$

which is a classical choice for vague priors with positive support [17].

Since the replicates are independent, the likelihood of the parameters of the Poisson-beta model, for a data-set of $N$ measurements $y_{1:N}^{(k)}$, is

$$\mathcal{L}'_k(y_{1:N}^{(k)}|\theta^{(k)}, \mu_X, k_{\mathrm{on}}, k_{\mathrm{off}}) = \prod_{j=1}^{N}\left(\sum_x f_\epsilon(y_j^{(k)} - \kappa x|a^{(k)}, \mu_\epsilon^{(k)}, \sigma_\epsilon^{(k)}) f'_X(x|\mu_X, k_{\mathrm{on}}, k_{\mathrm{off}})\right), \tag{31}$$

TABLE S4. Estimated coefficients (means and standard errors of the mean, $\mu_\kappa^{(k)}$ and $s_\kappa^{(k)}$, respectively) of the gamma GLM.

| k | gene | $\mu_\kappa^{(k)}$ | $s_\kappa^{(k)}$ |
|---|---|---|---|
| 1 | HBB WT | 20.904 | 0.230 |
| 2 | HBB WT | 27.080 | 0.283 |
| 3 | HBB WT | 28.394 | 0.241 |
| 4 | HBB WT | 21.631 | 0.244 |
| 1 | HBB mut | 20.308 | 0.233 |
| 2 | HBB mut | 29.621 | 0.402 |
| 3 | HBB mut | 32.267 | 0.353 |
| 4 | HBB mut | 26.612 | 0.322 |
| 1 | HIV WT | 17.715 | 0.153 |
| 2 | HIV WT | 34.771 | 0.341 |
| 3 | HIV WT | 17.118 | 0.183 |
| 4 | HIV WT | 50.748 | 0.547 |
| 1 | HIV mut | 23.035 | 0.219 |
| 2 | HIV mut | 32.463 | 0.381 |
| 3 | HIV mut | 26.951 | 0.296 |
| 4 | HIV mut | 25.487 | 0.300 |

where $\theta^{(k)} := (\kappa, a^{(k)}, \mu_\epsilon^{(k)}, \sigma_\epsilon^{(k)})$ is the vector of the parameters that describe the experimental setting. This completes the definition of the first Bayesian model for the observed data. The directed acyclic graph (DAG) of the full posterior of this model is illustrated in Fig. S5(A).

Consistently, the likelihood of the parameters of the negative-binomial model is

$$\mathcal{L}_k''(y_{1:N}^{(k)}|\theta^{(k)}, \mu_X, k_{\text{on}}) = \prod_{j=1}^{N} \left( \sum_x f_\epsilon(y_j^{(k)} - \kappa x | a^{(k)}, \mu_\epsilon^{(k)}, \sigma_\epsilon^{(k)}) f_X''(x|\mu_X, k_{\text{on}}) \right), \tag{32}$$

as illustrated in Fig. S5(B). The simplest Poisson model, the likelihood is

$$\mathcal{L}_k'''(y_{1:N}^{(k)}|\theta^{(k)}, \mu_X) = \prod_{j=1}^{N} \left( \sum_x f_\epsilon(y_j^{(k)} - \kappa x | a^{(k)}, \mu_\epsilon^{(k)}, \sigma_\epsilon^{(k)}) f_X'''(x|\mu_X) \right), \tag{33}$$

whose DAG is illustrated in Fig. S5(C).

### S4. MONTE CARLO ESTIMATION

#### A. MCMC samplers

Adaptive Metropolis–Hastings samplers to fit the model to the data where implemented using the PyMC library for probabilistic programming [18], version `2.3.7`, which has a flexible object oriented syntax and provides tools to handle long traces and perform diagnostics. To improve the convergence speed, the array containing the $N$ elements of a dataset was numerically sorted by value and split into $M = N/10$ blocks of size 10. The latent random-variable arrays $P_{1:N}$ and $X_{1:N}$ were batched too, as the RVs conditioned on the data of a single block are strongly correlated and are conveniently updated during single Metropolis–Hastings steps. Using the symbol $\mathbf{1}_x$ for the identity matrix and $\mathcal{N}_x$ to represent a multivariate normal RV of dimension $x$, the simulation of posterior samples for the Poisson-beta model proceeds as follows:

- For $i = 0, 1, \ldots, M$, values of the 10-value blocks $P_{(i\,10+1):(i+1)10}$ and $(X_{i\,10+1):(i+1)10}$ are updated according to a random walk Metropolis with proposals $\mathcal{N}_{10}(\mu_i^{(P)}, \sigma_i^{(P)}\,\mathbf{1}_{10})$ and $\mathcal{N}_{10}(\mu_i^{(X)}, \sigma_i^{(X)}\,\mathbf{1}_{10})$ respectively,

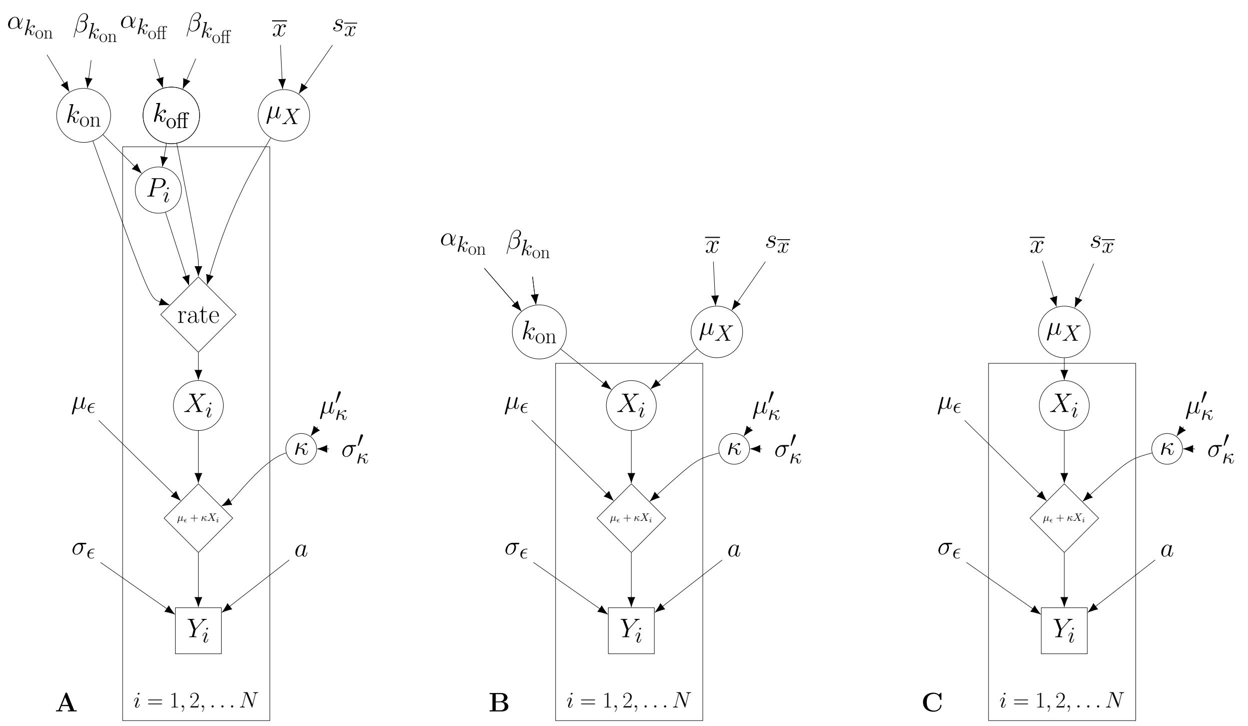


FIG. S5. DAG for the Poisson-beta model (A), the negative-binomial model (B), and the naïve Poisson model (C) with measurement equation. Circle nodes represent parameters to be estimated, blank nodes represent set parameters, diamonds correspond to deterministic functions of their parents, and square nodes represent observations.

- The RVs $\mu_X$, $\kappa$, $k_{\text{on}}$, and $k_{\text{off}}$ are updated as a single block according to the adaptive Metropolis–Hastings method of Ref. [19] with proposal $\mathcal{N}_4(\mu, \Sigma)$.

The proposal parameters $\mu_i^{(X)}$, $\sigma_i^{(X)}$, $\mu_i^{(P)}$, $\sigma_i^{(P)}$, $(i = 1, \ldots, M)$, $\mu$, and $\Sigma$ are chosen adaptively. To improve the adaptation (noting that the posterior for $k_{\text{off}}$ is more disperse than those of $\mu_X$, $\kappa$, and $k_{\text{on}}$), $\Sigma$ is initialised to the diagonal matrix $\text{diag}(0.1, 0.1, 0.1, 1)$.

In order to mitigate tractability issues (which is mainly due to the large number of presence of latent variables), the model is only fitted to a randomly sampled subset of $N = 500$ data points. For a more modern approach to cope with latent variables see Ref. [7], which also defines a more complex hierarchical model.

The sampler implemented for the negative-binomial model is similar to the one implemented for the Poisson-beta model (with data organised into $M$ ranked batches) but converges and mixes more rapidly, as it does not encode for the latent variables $P_i$, $i = 1, 2, \ldots, N$. The simulation proceeds as follows:

- For $i = 0, 1, \ldots, M$, values of $X_{(i\,10+1):(i+1)\,10}$ are updated according to a random-walk Metropolis with proposal $\mathcal{N}_{10}(\mu_i^{(X)}, \sigma_i^{(X)}\,\mathbf{1}_{10})$.
- The random variables $\mu_X, \kappa$ and $k_{\text{on}}$ are updated simultaneously according to the adaptive Metropolis–Hastings method with proposal $\mathcal{N}_3(\mu, \Sigma)$,

where the quantities $\mathbf{1}_x$, $\mu_i^{(X)}$, $\sigma_i^{(X)}$, $(i = 1, \ldots, M)$, $\mu$, and $\Sigma$ are defined as in the former case. The Poisson-model sampler is analogous to the negative binomial, except that it does not include the parameter $k_{\text{on}}$. All the samplers were successfully tested with simulated data.

### B. Consensus posteriors

The posterior

$$p(\vartheta | y_{1:N}^{(1)}, y_{1:N}^{(2)} y_{1:N}^{(3)}, y_{1:N}^{(4)}) \propto \prod_{k=1}^{4} \left[ \mathcal{L}_k(y_{1:N}^{(k)} | \vartheta) \right] p(\vartheta) \tag{34}$$

represents the consensus belief on the vector of parameters $\vartheta$ among all the replicates $k = 1, 2, 3, 4$, with $p(\vartheta)$ being the prior. We approximate such a posterior by means of a consensus Monte Carlo approach, i.e., by running a separate MCMC on each of

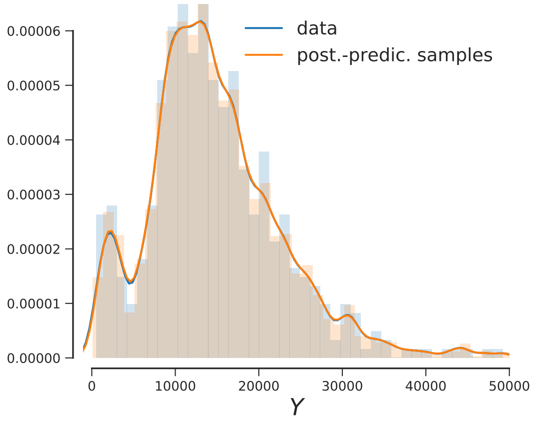


FIG. S6. KDE and histogram of the data (blue) (HBB, WT, 80 ng/mL Tet, $k = 1$) and of the draws from the corresponding posterior-predictive distribution (orange), according to the negative-binomial model. It is possible to visually appreciate their overlap. To quantify the extent to which the posterior predictive reproduces the true data distribution for all the fit, we studied the Wasserstein distance of the two distribution.

the datasets $y^{(k)}_{1:N}$, $k = 1, 2, 3, 4$, and then averaging individual Monte Carlo draws across, as in Ref. [20]. The draws $\vartheta^{(k)}_j$, $j = 1, \ldots, L$, $k = 1, 2, 3, 4$, are combined into the weighted averages

$$\vartheta_j = \frac{\sum_k \vartheta^{(k)}_j w_{(k)}}{\sum_k w_{(k)}}, \tag{35}$$

where $w_{(k)}$ is the vector of the reciprocal of the marginal posterior variances. This method has been only justified rigorously for Gaussian posteriors and only yields approximate posteriors, in general. However, it allowed us to distribute the Bayesian analysis across different machines , and therefore was of great utility to aggregate results.

#### C. Goodness of fit

To evaluate the goodness of fit (GoF) we estimate the posterior predictive distribution

$$p(\tilde{y}^{(k)}_{1:N}|y^{(k)}_{1:N}) = \int p(\tilde{y}^{(k)}_{1:N}|\theta)p(\theta|y^{(k)}_{1:N})\, \mathrm{d}\theta, \tag{36}$$

where $\theta$ is the vector of all parameters, by generating pseudo-data $\tilde{y}^{(k)}_{1:N}$ for the model using the parameters drawn from the posterior $p(\theta|y^{(k)}_{1:N})$ alongside each MCMC chain. A GoF test follows by measuring to what extent the pseudo-data deviate from $y^{(k)}_{1:N}$. Specifically, we calculate the root mean square displacement (RMSD) of the data $y^{(k)}_i$, $i = 1, \ldots, N$ with respect to the sample mean $\bar{y}^{(k)}_i$ of the draws from the marginal posterior predictive, i.e.,

$$\mathrm{RMSD}(\bar{y}^{(k)}_{1:N}, y^{(k)}_{1:N}) = \sqrt{\frac{1}{N}\sum_{i=1}^{N}(\bar{y}^{(k)}_i - y^{(k)}_i)^2}. \tag{37}$$

Comparison between the RMSD results for the Poisson-beta and the negative-binomial models is shown in Fig. S7, which suggests that both these models fit the data equally well. Conversely, the RMSDs for the Poisson model are always higher that the RMSDs for the two former models (see Fig. S7) implying that the Poisson model does not fit as well. Further, we computed the Wasserstein distance in distribution between the data and the pseudo-data. The Wasserstein distance between two distributions $u$ and $v$ is defined as

$$l_1(u, v) = \int_{-\infty}^{\infty} |U - V|, \tag{38}$$

where $U$ and $V$ are the empirical cumulative distribution functions associated to and $u$ and $v$, respectively. Fig. S7 shows that the distances are always smaller than the 95% percentile of bootstrap-samples distances from the true data, thus confirming GoF. Bin sizes for the empirical distribution were chosen according to the Freedman–Diaconis rule.

### S5. MRNA DECAY RATES

The draws from the posteriors of the dimensionless rates $k_{\rm on}$, $k_{\rm off}$, and $\alpha$ are converted to number of events per minute $\tilde{k}_{\rm on}$, $\tilde{k}_{\rm off}$, and $\tilde{\alpha}$ by using estimated decay rates of mRNA. For the HBB gene, decay rates were measured in Ref. [22] (and are reported in Table S5). Due to the detection of two different mRNA isoforms, viz., "rt" and "pA", the empirical mRNA distribution can be thought of a Gaussian mixture density with PDF

$$f(x) = p_{\rm pA}\, \phi(x|\mu_{\rm pA}, \sigma_{\rm pA}) + p_{\rm rt}\, \phi(x|\mu_{\rm rt}, \sigma_{\rm rt}), \tag{39}$$

$p_{\rm rt}+p_{\rm rt} = 1$, with parameters of Table S5. According to equations (2)-(4), the traces from $k_{\rm on}$, $k_{\rm off}$, and $\alpha$ were multiplied by draws from this Gaussian mixture to obtain $\tilde{k}_{\rm on}$, $\tilde{k}_{\rm off}$, and $\tilde{\alpha}$.

We measured total *env* RNA content (including non-poly-adenylated transcripts) with RT-qPCR, using gene-specific primers (forward primers bind exon 1 and reverse primers bind the 3'UTR, see section S10 C). The decay rates of *env* transcripts were obtained by fitting a linear model to the logarithm of RT-qPCR measurements of transcripts vs time in

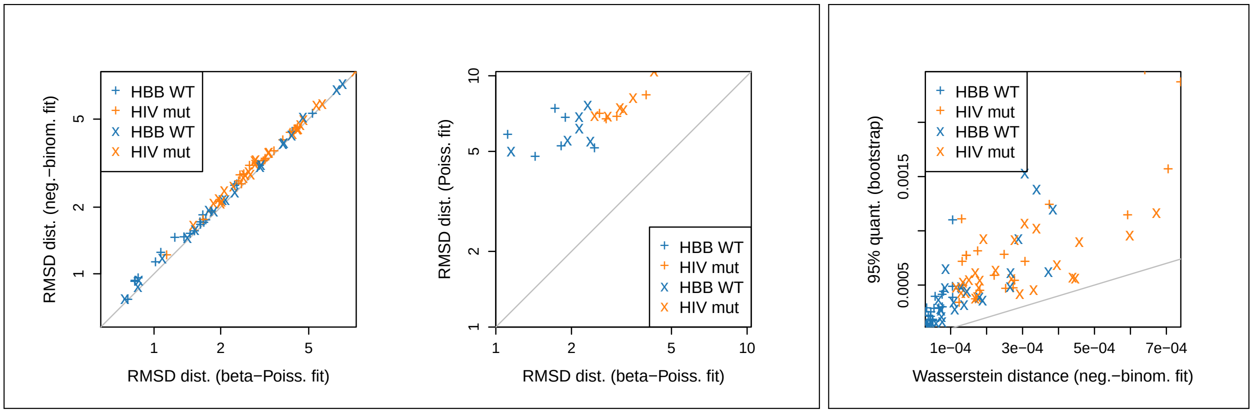


FIG. S7. GoF analysis. (left and centre) RMSDs of the data with respect to the sample means of the (posterior predictive (see eq. (37)) for each dataset. Comparison of Poisson-beta model vs the negative-binomial model (left scatter plot) shows that the two models achieve similar GoF, while the RMSDs obtained from the Poisson model are always the largest (central scatter plot). (right) Wasserstein distances between the empirical histogram of the data and the negative-binomial model posterior predictive ($x$-axis) is always smaller that the 95% quantiles of the bootstrapped distances from the true data, which suggests that that the posterior-predictive samples for the negative-binomial model always reproduce the true flow-FISH data (as in, e.g., Fig. S6).

TABLE S5. . Decay rates in number of events per minutes of the two mRNA isoforms for the HBB gene. "pA" and "rt" refer to polyadenylated and read-through isoforms, respectively (it is possible to foresee larger dispersion for the mutant than for the WT).

| SNP | isoform | $\mu$ | $\sigma$ | $p$ |
|---|---|---|---|---|
| WT | pA | 0.0024 | 0.0002 | 0.72 |
| WT | rt | 0.0067 | 0.0016 | 0.28 |
| mut | pA | 0.0036 | 0.0003 | 0.11 |
| mut | rt | 0.0112 | 0.0023 | 0.89 |

minutes. The inferred mean $\mu_d$ and standard error $\sigma_d$ of the decay rates are reported in Fig. S8. The draws from the posteriors of dimensionless rates $k_{\rm on}$, $k_{\rm off}$, and $\alpha$ were converted to number of events per minute by multiplying by normal draws $\mathcal{N}(\mu_d, \sigma_d)$, with parameters listed in Fig. S8.

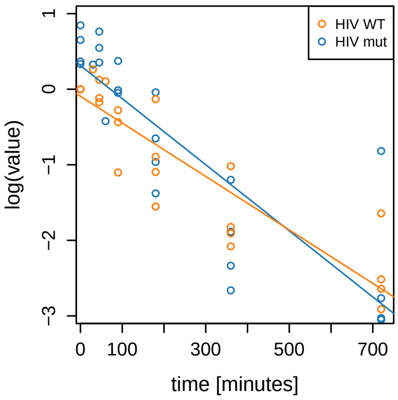


| SNP | $\mu_d$ | $\sigma_d$ |
|---|---|---|
| WT | 0.0044 | 6e-04 |
| mut | 0.0035 | 4e-04 |

FIG. S8. Logarithm of RT-qPCR $2^{-\Delta\Delta C_t}$ measurements of residual transcripts vs time of the mutant and WT HIV gene transcripts. From linear regression, decay rates in unit of events per minutes are obtained.

### S6. SUMMARY OF MCMC RESULTS

#### A. Poisson-beta distribution

Fig. S9 shows two results obtained from fitting the Poisson-beta model. The traces of the posterior chains from each replicate were combined according to the consensus Monte Carlo procedure (see section S4 B) to obtain a representation of the consensus belief of Fig. 3 (*Main text*). It is worth noting that the credible intervals for $\tilde{k}_{\rm off}$ and $\tilde{\alpha}$ are very wide, while the MCMC draws of $k_{\rm off}$ and $\alpha$ appear strongly cross-correlated (see, e.g., Fig. S10), where the drawn samples form an angle $\mathrm{arccot}(\alpha/k_{\rm off})$ with the abscissae axis.

The 90% highest posterior density credible intervals (HPD CIs) and medians of the estimated parameters $\kappa$, $\mu_X$, $k_{\rm on}$, $k_{\rm off}$, and $\alpha/k_{\rm off}$ are reported in Tables S6-S10 (HBB gene) and Tables S11-S15 (HIV gene).

TABLE S6. $\kappa$ of HBB gene

| k | SNP | tet | low.HPD | median | upp.HPD |
|---|---|---|---|---|---|
| 1 | WT | 5 | 9.559 | 11.257 | 13.031 |
| 1 | WT | 10 | 22.384 | 25.504 | 28.796 |
| 1 | WT | 20 | 18.866 | 20.662 | 22.473 |
| 1 | WT | 40 | 20.378 | 22.284 | 24.023 |
| 1 | WT | 80 | 25.675 | 27.393 | 29.289 |
| 1 | WT | 250 | 17.021 | 18.443 | 19.965 |
| 1 | mut | 5 | 32.504 | 35.358 | 38.254 |
| 1 | mut | 10 | 16.315 | 18.734 | 21.498 |
| 1 | mut | 40 | 23.342 | 25.416 | 27.553 |
| 1 | mut | 80 | 27.117 | 30.304 | 33.601 |
| 1 | mut | 250 | 17.047 | 18.166 | 19.282 |
| 2 | WT | 5 | 23.567 | 26.676 | 30.352 |
| 2 | WT | 10 | 22.676 | 24.647 | 28.696 |
| 2 | WT | 20 | 21.508 | 23.394 | 25.370 |
| 2 | WT | 40 | 25.727 | 27.715 | 29.846 |
| 2 | WT | 80 | 32.172 | 34.216 | 35.902 |
| 2 | WT | 250 | 24.276 | 25.938 | 27.678 |
| 2 | mut | 5 | 26.707 | 29.680 | 32.950 |
| 2 | mut | 10 | 31.601 | 36.919 | 41.979 |
| 2 | mut | 40 | 41.755 | 44.750 | 48.189 |
| 2 | mut | 80 | 27.094 | 30.426 | 33.609 |
| 2 | mut | 250 | 22.821 | 24.260 | 25.754 |
| 3 | WT | 5 | 25.938 | 29.050 | 32.440 |
| 3 | WT | 10 | 32.783 | 32.859 | 38.434 |
| 3 | WT | 20 | 30.479 | 32.865 | 35.207 |
| 3 | WT | 40 | 30.875 | 32.961 | 35.374 |
| 3 | WT | 80 | 25.001 | 26.456 | 28.116 |
| 3 | WT | 250 | 20.897 | 21.763 | 23.000 |
| 3 | mut | 5 | 24.799 | 27.457 | 30.501 |
| 3 | mut | 10 | 42.772 | 47.404 | 51.766 |
| 3 | mut | 40 | 35.653 | 38.381 | 41.145 |
| 3 | mut | 80 | 31.024 | 34.699 | 38.285 |
| 3 | mut | 250 | 28.080 | 29.963 | 31.857 |
| 4 | WT | 5 | 18.060 | 20.483 | 23.184 |
| 4 | WT | 10 | 17.540 | 19.780 | 22.412 |
| 4 | WT | 20 | 23.540 | 23.855 | 24.425 |
| 4 | WT | 40 | 30.582 | 32.931 | 35.604 |
| 4 | WT | 80 | 29.660 | 31.397 | 33.534 |
| 4 | WT | 250 | 23.480 | 25.104 | 26.733 |
| 4 | mut | 5 | 14.313 | 16.032 | 17.731 |
| 4 | mut | 10 | 23.044 | 26.306 | 30.166 |
| 4 | mut | 40 | 32.880 | 35.456 | 38.080 |
| 4 | mut | 80 | 28.650 | 31.858 | 35.336 |
| 4 | mut | 250 | 26.352 | 27.831 | 29.268 |

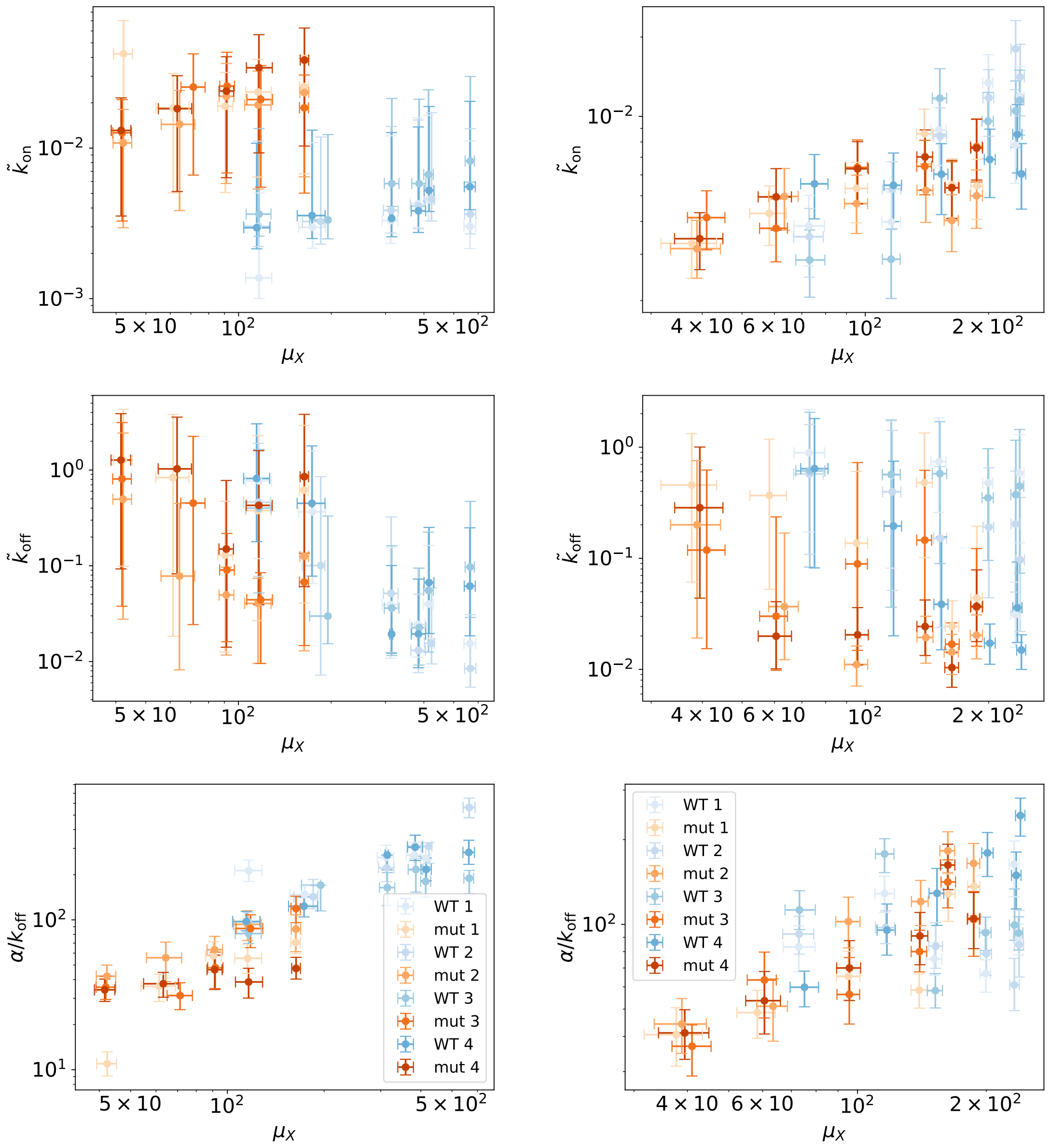


**A** HBB. **B** HIV.

FIG. S9. Estimates for parameters $\tilde{k}_{\rm on}$, $\tilde{k}_{\rm off}$, $\mu_X$, and $\alpha/k_{\rm off}$ of the Poisson- beta model, from wild-type (blue) and mutant (orange) cell data, for all induction levels, shades of colors correspond to replicates. Points are medians, error bars comprise 90% HPD CIs. HBB-gene results show results consistent across all the replicates (panel **A**). The HIB-gene results are reported in panels **B**. Increasing expression levels, three of the HIV replicates show a drop-off in the average burst size and an increase in the burst, see also Fig. S2. The consensus estimates are reported in Fig. 3, *Main text*.

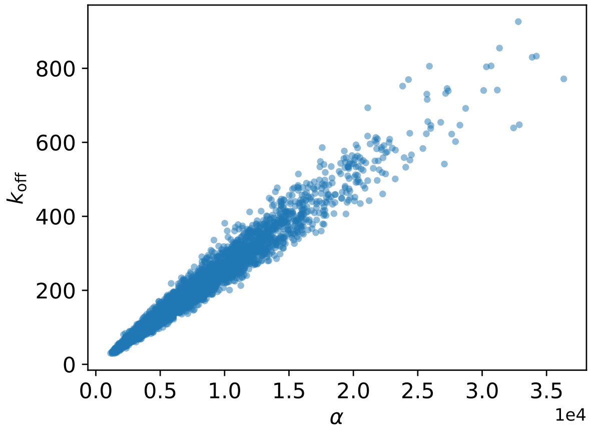


FIG. S10. Cross-correlation between the MCMC draws for the dimensionless parameters $k_{\mathrm{off}}$ and $\alpha$.

TABLE S7. $\mu_X$ HBB

| k | SNP | tet | low.HPD | median | upp.HPD |
|---|---|---|---|---|---|
| 1 | WT | 5 | 105.402 | 116.589 | 128.100 |
| 1 | WT | 10 | 156.821 | 173.998 | 191.793 |
| 1 | WT | 20 | 292.726 | 311.989 | 329.181 |
| 1 | WT | 40 | 360.207 | 382.007 | 402.900 |
| 1 | WT | 80 | 398.889 | 414.140 | 429.588 |
| 1 | WT | 250 | 539.820 | 564.476 | 589.263 |
| 1 | mut | 5 | 39.239 | 42.300 | 45.212 |
| 1 | mut | 10 | 53.895 | 61.338 | 69.030 |
| 1 | mut | 40 | 85.582 | 90.517 | 95.846 |
| 1 | mut | 80 | 105.474 | 115.611 | 127.808 |
| 1 | mut | 250 | 157.627 | 163.197 | 168.546 |
| 2 | WT | 5 | 103.760 | 116.053 | 126.933 |
| 2 | WT | 10 | 161.211 | 184.961 | 190.736 |
| 2 | WT | 20 | 295.510 | 313.086 | 329.837 |
| 2 | WT | 40 | 362.581 | 384.120 | 404.709 |
| 2 | WT | 80 | 401.521 | 423.421 | 432.840 |
| 2 | WT | 250 | 542.177 | 565.898 | 589.598 |
| 2 | mut | 5 | 39.110 | 42.207 | 44.976 |
| 2 | mut | 10 | 56.058 | 64.338 | 72.119 |
| 2 | mut | 40 | 86.483 | 91.288 | 96.615 |
| 2 | mut | 80 | 104.644 | 115.765 | 127.515 |
| 2 | mut | 250 | 158.168 | 163.425 | 168.913 |
| 3 | WT | 5 | 105.803 | 116.504 | 127.453 |
| 3 | WT | 10 | 170.108 | 195.097 | 195.097 |
| 3 | WT | 20 | 297.477 | 314.601 | 331.874 |
| 3 | WT | 40 | 365.211 | 385.872 | 405.793 |
| 3 | WT | 80 | 399.257 | 413.887 | 429.090 |
| 3 | WT | 250 | 544.380 | 565.718 | 583.439 |
| 3 | mut | 5 | 39.047 | 41.876 | 44.904 |
| 3 | mut | 10 | 64.976 | 71.285 | 77.862 |
| 3 | mut | 40 | 86.882 | 91.723 | 96.974 |
| 3 | mut | 80 | 106.385 | 118.060 | 128.903 |
| 3 | mut | 250 | 157.720 | 163.555 | 168.630 |
| 4 | WT | 5 | 103.952 | 114.542 | 126.282 |
| 4 | WT | 10 | 155.066 | 173.247 | 191.033 |
| 4 | WT | 20 | 309.627 | 314.120 | 316.703 |
| 4 | WT | 40 | 363.798 | 383.977 | 404.975 |
| 4 | WT | 80 | 400.513 | 415.609 | 431.024 |
| 4 | WT | 250 | 540.749 | 564.065 | 587.349 |
| 4 | mut | 5 | 38.580 | 41.557 | 44.668 |
| 4 | mut | 10 | 54.873 | 63.206 | 70.399 |
| 4 | mut | 40 | 86.438 | 91.326 | 96.352 |
| 4 | mut | 80 | 106.004 | 116.457 | 128.697 |
| 4 | mut | 250 | 158.561 | 163.656 | 168.862 |

TABLE S8. $\tilde{k}_{\text{on}}$ HBB gene

| k | SNP | tet | low.HPD | median | upp.HPD |
|---|---|---|---|---|---|
| 1 | WT | 5 | 0.001 | 0.001 | 0.005 |
| 1 | WT | 10 | 0.002 | 0.003 | 0.011 |
| 1 | WT | 20 | 0.002 | 0.003 | 0.012 |
| 1 | WT | 40 | 0.003 | 0.004 | 0.016 |
| 1 | WT | 80 | 0.003 | 0.005 | 0.016 |
| 1 | WT | 250 | 0.002 | 0.003 | 0.011 |
| 1 | mut | 5 | 0.012 | 0.042 | 0.072 |
| 1 | mut | 10 | 0.005 | 0.018 | 0.031 |
| 1 | mut | 40 | 0.005 | 0.019 | 0.032 |
| 1 | mut | 80 | 0.006 | 0.023 | 0.038 |
| 1 | mut | 250 | 0.007 | 0.026 | 0.042 |
| 2 | WT | 5 | 0.002 | 0.003 | 0.011 |
| 2 | WT | 10 | 0.002 | 0.003 | 0.012 |
| 2 | WT | 20 | 0.003 | 0.004 | 0.014 |
| 2 | WT | 40 | 0.003 | 0.004 | 0.015 |
| 2 | WT | 80 | 0.003 | 0.005 | 0.016 |
| 2 | WT | 250 | 0.003 | 0.004 | 0.013 |
| 2 | mut | 5 | 0.003 | 0.011 | 0.018 |
| 2 | mut | 10 | 0.004 | 0.014 | 0.024 |
| 2 | mut | 40 | 0.005 | 0.022 | 0.037 |
| 2 | mut | 80 | 0.006 | 0.019 | 0.032 |
| 2 | mut | 250 | 0.006 | 0.024 | 0.039 |
| 3 | WT | 5 | 0.003 | 0.004 | 0.013 |
| 3 | WT | 10 | 0.002 | 0.003 | 0.012 |
| 3 | WT | 20 | 0.004 | 0.006 | 0.021 |
| 3 | WT | 40 | 0.004 | 0.006 | 0.021 |
| 3 | WT | 80 | 0.005 | 0.007 | 0.025 |
| 3 | WT | 250 | 0.006 | 0.008 | 0.031 |
| 3 | mut | 5 | 0.003 | 0.013 | 0.021 |
| 3 | mut | 10 | 0.007 | 0.026 | 0.042 |
| 3 | mut | 40 | 0.007 | 0.026 | 0.044 |
| 3 | mut | 80 | 0.006 | 0.021 | 0.035 |
| 3 | mut | 250 | 0.005 | 0.018 | 0.031 |
| 4 | WT | 5 | 0.002 | 0.003 | 0.011 |
| 4 | WT | 10 | 0.002 | 0.004 | 0.013 |
| 4 | WT | 20 | 0.003 | 0.003 | 0.012 |
| 4 | WT | 40 | 0.003 | 0.004 | 0.014 |
| 4 | WT | 80 | 0.004 | 0.005 | 0.019 |
| 4 | WT | 250 | 0.004 | 0.006 | 0.020 |
| 4 | mut | 5 | 0.003 | 0.013 | 0.021 |
| 4 | mut | 10 | 0.005 | 0.018 | 0.031 |
| 4 | mut | 40 | 0.006 | 0.024 | 0.040 |
| 4 | mut | 80 | 0.009 | 0.034 | 0.057 |
| 4 | mut | 250 | 0.010 | 0.038 | 0.063 |

TABLE S9. $\tilde{k}_{\text{off}}$ HBB gene

| k | SNP | tet | low.HPD | median | upp.HPD |
|---|---|---|---|---|---|
| 1 | WT | 5 | 0.125 | 0.765 | 3.017 |
| 1 | WT | 10 | 0.072 | 0.380 | 1.567 |
| 1 | WT | 20 | 0.012 | 0.039 | 0.154 |
| 1 | WT | 40 | 0.011 | 0.024 | 0.094 |
| 1 | WT | 80 | 0.016 | 0.040 | 0.161 |
| 1 | WT | 250 | 0.009 | 0.015 | 0.056 |
| 1 | mut | 5 | 0.073 | 1.278 | 4.126 |
| 1 | mut | 10 | 0.050 | 0.832 | 3.991 |
| 1 | mut | 40 | 0.013 | 0.130 | 0.474 |
| 1 | mut | 80 | 0.031 | 0.375 | 2.323 |
| 1 | mut | 250 | 0.042 | 0.622 | 3.073 |
| 2 | WT | 5 | 0.083 | 0.468 | 1.868 |
| 2 | WT | 10 | 0.004 | 0.103 | 0.843 |
| 2 | WT | 20 | 0.011 | 0.052 | 0.334 |
| 2 | WT | 40 | 0.007 | 0.013 | 0.048 |
| 2 | WT | 80 | 0.010 | 0.016 | 0.057 |
| 2 | WT | 250 | 0.005 | 0.008 | 0.031 |
| 2 | mut | 5 | 0.024 | 0.496 | 2.425 |
| 2 | mut | 10 | 0.006 | 0.077 | 0.442 |
| 2 | mut | 40 | 0.011 | 0.050 | 0.095 |
| 2 | mut | 80 | 0.010 | 0.040 | 0.073 |
| 2 | mut | 250 | 0.012 | 0.127 | 0.909 |
| 3 | WT | 5 | 0.050 | 0.399 | 1.601 |
| 3 | WT | 10 | 0.013 | 0.030 | 0.343 |
| 3 | WT | 20 | 0.011 | 0.035 | 0.156 |
| 3 | WT | 40 | 0.010 | 0.022 | 0.104 |
| 3 | WT | 80 | 0.013 | 0.055 | 0.230 |
| 3 | WT | 250 | 0.027 | 0.097 | 0.495 |
| 3 | mut | 5 | 0.036 | 0.807 | 3.286 |
| 3 | mut | 10 | 0.033 | 0.440 | 2.258 |
| 3 | mut | 40 | 0.013 | 0.091 | 0.224 |
| 3 | mut | 80 | 0.011 | 0.044 | 0.084 |
| 3 | mut | 250 | 0.014 | 0.067 | 0.137 |
| 4 | WT | 5 | 0.180 | 0.813 | 3.140 |
| 4 | WT | 10 | 0.095 | 0.443 | 1.732 |
| 4 | WT | 20 | 0.013 | 0.019 | 0.101 |
| 4 | WT | 40 | 0.009 | 0.019 | 0.072 |
| 4 | WT | 80 | 0.020 | 0.068 | 0.265 |
| 4 | WT | 250 | 0.019 | 0.060 | 0.254 |
| 4 | mut | 5 | 0.122 | 1.271 | 3.884 |
| 4 | mut | 10 | 0.072 | 1.031 | 3.537 |
| 4 | mut | 40 | 0.020 | 0.149 | 0.768 |
| 4 | mut | 80 | 0.049 | 0.431 | 1.596 |
| 4 | mut | 250 | 0.037 | 0.844 | 3.754 |

TABLE S10. $\alpha/k_{\text{off}}$ HBB

| k | SNP | tet | low.HPD | median | upp.HPD |
|---|---|---|---|---|---|
| 1 | WT | 5 | 180.624 | 213.178 | 250.384 |
| 1 | WT | 10 | 125.805 | 147.598 | 168.798 |
| 1 | WT | 20 | 216.330 | 261.287 | 314.761 |
| 1 | WT | 40 | 212.117 | 270.985 | 329.730 |
| 1 | WT | 80 | 214.432 | 257.880 | 304.762 |
| 1 | WT | 250 | 477.217 | 564.684 | 648.593 |
| 1 | mut | 5 | 9.080 | 10.960 | 13.174 |
| 1 | mut | 10 | 28.529 | 36.058 | 43.356 |
| 1 | mut | 40 | 44.662 | 57.687 | 71.459 |
| 1 | mut | 80 | 43.999 | 55.447 | 69.005 |
| 1 | mut | 250 | 59.522 | 70.628 | 84.777 |
| 2 | WT | 5 | 82.844 | 96.710 | 111.647 |
| 2 | WT | 10 | 118.521 | 142.328 | 186.000 |
| 2 | WT | 20 | 180.061 | 222.460 | 269.372 |
| 2 | WT | 40 | 256.229 | 310.385 | 368.848 |
| 2 | WT | 80 | 238.431 | 309.659 | 327.796 |
| 2 | WT | 250 | 481.478 | 561.546 | 649.773 |
| 2 | mut | 5 | 35.331 | 42.066 | 49.954 |
| 2 | mut | 10 | 38.750 | 55.886 | 71.052 |
| 2 | mut | 40 | 47.767 | 63.226 | 77.583 |
| 2 | mut | 80 | 72.672 | 93.342 | 115.105 |
| 2 | mut | 250 | 61.357 | 86.761 | 108.005 |
| 3 | WT | 5 | 69.549 | 81.039 | 94.196 |
| 3 | WT | 10 | 114.861 | 170.317 | 170.356 |
| 3 | WT | 20 | 124.441 | 164.157 | 206.865 |
| 3 | WT | 40 | 153.175 | 216.801 | 264.885 |
| 3 | WT | 80 | 141.653 | 180.708 | 226.265 |
| 3 | WT | 250 | 149.771 | 189.476 | 213.603 |
| 3 | mut | 5 | 29.455 | 35.547 | 42.080 |
| 3 | mut | 10 | 25.170 | 31.255 | 38.079 |
| 3 | mut | 40 | 34.828 | 47.958 | 60.944 |
| 3 | mut | 80 | 65.273 | 87.569 | 108.009 |
| 3 | mut | 250 | 95.857 | 119.091 | 143.867 |
| 4 | WT | 5 | 84.599 | 97.546 | 113.826 |
| 4 | WT | 10 | 104.599 | 123.292 | 142.147 |
| 4 | WT | 20 | 215.483 | 270.814 | 270.814 |
| 4 | WT | 40 | 249.124 | 305.273 | 366.964 |
| 4 | WT | 80 | 182.471 | 217.225 | 258.564 |
| 4 | WT | 250 | 234.660 | 281.937 | 339.927 |
| 4 | mut | 5 | 28.552 | 34.146 | 40.265 |
| 4 | mut | 10 | 30.453 | 37.481 | 44.636 |
| 4 | mut | 40 | 34.310 | 46.529 | 58.219 |
| 4 | mut | 80 | 30.063 | 38.528 | 47.509 |
| 4 | mut | 250 | 40.276 | 47.439 | 56.269 |

TABLE S11. $\kappa$ HIV

| k | SNP | tet | low.HPD | median | upp.HPD |
|---|---|---|---|---|---|
| 1 | WT | 5 | 8.727 | 9.937 | 11.325 |
| 1 | WT | 10 | 10.993 | 12.288 | 13.592 |
| 1 | WT | 20 | 16.932 | 18.302 | 19.718 |
| 1 | WT | 40 | 18.385 | 19.903 | 20.011 |
| 1 | WT | 80 | 16.735 | 17.896 | 18.957 |
| 1 | WT | 250 | 24.550 | 25.974 | 27.432 |
| 1 | mut | 5 | 16.621 | 19.959 | 23.859 |
| 1 | mut | 10 | 21.241 | 23.881 | 27.318 |
| 1 | mut | 20 | 22.935 | 25.057 | 27.444 |
| 1 | mut | 40 | 19.246 | 20.688 | 22.178 |
| 1 | mut | 80 | 26.489 | 28.430 | 30.467 |
| 1 | mut | 250 | 24.061 | 25.942 | 27.678 |
| 2 | WT | 5 | 15.392 | 17.512 | 19.808 |
| 2 | WT | 10 | 25.774 | 28.301 | 30.527 |
| 2 | WT | 20 | 36.093 | 38.649 | 40.924 |
| 2 | WT | 40 | 34.004 | 35.801 | 37.798 |
| 2 | WT | 80 | 30.599 | 31.966 | 33.382 |
| 2 | WT | 250 | 37.518 | 39.354 | 41.293 |
| 2 | mut | 5 | 21.794 | 25.773 | 29.947 |
| 2 | mut | 10 | 43.278 | 47.496 | 52.058 |
| 2 | mut | 20 | 21.203 | 23.231 | 25.227 |
| 2 | mut | 40 | 30.657 | 32.984 | 35.508 |
| 2 | mut | 80 | 19.609 | 21.327 | 22.957 |
| 2 | mut | 250 | 16.493 | 17.664 | 18.954 |
| 3 | WT | 5 | 10.456 | 12.000 | 13.644 |
| 3 | WT | 10 | 9.830 | 10.997 | 12.398 |
| 3 | WT | 20 | 29.612 | 31.544 | 33.564 |
| 3 | WT | 40 | 15.692 | 16.768 | 17.845 |
| 3 | WT | 80 | 16.705 | 17.712 | 18.798 |
| 3 | WT | 250 | 19.341 | 20.192 | 21.265 |
| 3 | mut | 5 | 33.804 | 38.444 | 42.521 |
| 3 | mut | 10 | 29.922 | 32.648 | 36.088 |
| 3 | mut | 20 | 25.741 | 28.254 | 30.406 |
| 3 | mut | 40 | 21.824 | 23.662 | 25.236 |
| 3 | mut | 80 | 21.917 | 23.597 | 25.190 |
| 3 | mut | 250 | 16.718 | 17.810 | 18.906 |
| 4 | WT | 5 | 34.743 | 38.418 | 42.457 |
| 4 | WT | 10 | 37.419 | 40.703 | 43.863 |
| 4 | WT | 20 | 42.950 | 46.185 | 49.408 |
| 4 | WT | 40 | 48.250 | 51.426 | 54.571 |
| 4 | WT | 80 | 54.240 | 57.360 | 60.494 |
| 4 | WT | 250 | 53.328 | 56.927 | 60.132 |
| 4 | mut | 5 | 25.447 | 29.942 | 34.647 |
| 4 | mut | 10 | 34.322 | 38.062 | 42.392 |
| 4 | mut | 20 | 28.296 | 30.815 | 33.340 |
| 4 | mut | 40 | 26.199 | 28.116 | 29.970 |
| 4 | mut | 80 | 20.818 | 22.328 | 23.790 |
| 4 | mut | 250 | 20.302 | 21.626 | 22.973 |

TABLE S12. $\mu_X$ HIV gene

| k | SNP | tet | low.HPD | median | upp.HPD |
|---|---|---|---|---|---|
| 1 | WT | 5 | 67.005 | 72.838 | 79.536 |
| 1 | WT | 10 | 109.810 | 115.686 | 121.952 |
| 1 | WT | 20 | 144.574 | 151.166 | 157.431 |
| 1 | WT | 40 | 194.485 | 198.005 | 203.512 |
| 1 | WT | 80 | 226.244 | 232.415 | 238.770 |
| 1 | WT | 250 | 231.469 | 238.032 | 244.048 |
| 1 | mut | 5 | 31.686 | 37.672 | 43.353 |
| 1 | mut | 10 | 52.227 | 58.303 | 64.193 |
| 1 | mut | 20 | 89.312 | 95.185 | 101.282 |
| 1 | mut | 40 | 133.357 | 139.395 | 145.681 |
| 1 | mut | 80 | 156.590 | 162.956 | 169.112 |
| 1 | mut | 250 | 180.919 | 187.275 | 193.192 |
| 2 | WT | 5 | 66.834 | 73.001 | 78.969 |
| 2 | WT | 10 | 110.155 | 116.385 | 122.115 |
| 2 | WT | 20 | 146.758 | 152.367 | 157.963 |
| 2 | WT | 40 | 193.705 | 200.206 | 205.751 |
| 2 | WT | 80 | 226.944 | 232.737 | 238.957 |
| 2 | WT | 250 | 233.022 | 238.787 | 244.544 |
| 2 | mut | 5 | 33.431 | 38.818 | 44.292 |
| 2 | mut | 10 | 58.085 | 63.467 | 68.566 |
| 2 | mut | 20 | 88.921 | 95.219 | 101.320 |
| 2 | mut | 40 | 133.678 | 140.521 | 146.283 |
| 2 | mut | 80 | 155.778 | 162.631 | 168.680 |
| 2 | mut | 250 | 180.361 | 186.879 | 193.168 |
| 3 | WT | 5 | 67.657 | 73.147 | 79.668 |
| 3 | WT | 10 | 110.114 | 115.655 | 121.652 |
| 3 | WT | 20 | 145.713 | 152.053 | 157.986 |
| 3 | WT | 40 | 192.742 | 199.274 | 205.308 |
| 3 | WT | 80 | 226.620 | 233.022 | 238.193 |
| 3 | WT | 250 | 232.351 | 238.150 | 244.483 |
| 3 | mut | 5 | 36.811 | 40.994 | 45.435 |
| 3 | mut | 10 | 55.191 | 60.551 | 64.555 |
| 3 | mut | 20 | 89.503 | 95.670 | 101.158 |
| 3 | mut | 40 | 133.490 | 139.695 | 145.130 |
| 3 | mut | 80 | 156.822 | 162.594 | 169.538 |
| 3 | mut | 250 | 180.569 | 186.922 | 192.906 |
| 4 | WT | 5 | 69.528 | 75.142 | 81.061 |
| 4 | WT | 10 | 111.093 | 117.156 | 122.570 |
| 4 | WT | 20 | 146.982 | 153.429 | 159.127 |
| 4 | WT | 40 | 195.212 | 201.345 | 207.570 |
| 4 | WT | 80 | 229.161 | 235.061 | 241.236 |
| 4 | WT | 250 | 234.182 | 240.462 | 246.342 |
| 4 | mut | 5 | 34.228 | 39.432 | 44.903 |
| 4 | mut | 10 | 54.750 | 60.567 | 66.025 |
| 4 | mut | 20 | 89.372 | 95.749 | 101.674 |
| 4 | mut | 40 | 133.614 | 139.938 | 146.017 |
| 4 | mut | 80 | 156.317 | 162.691 | 168.913 |
| 4 | mut | 250 | 180.968 | 186.969 | 193.373 |

TABLE S13. $\tilde{k}_{on}$ HIV gene

| k | SNP | tet | low.HPD | median | upp.HPD |
|---|---|---|---|---|---|
| 1 | WT | 5 | 0.003 | 0.004 | 0.005 |
| 1 | WT | 10 | 0.003 | 0.004 | 0.005 |
| 1 | WT | 20 | 0.006 | 0.009 | 0.011 |
| 1 | WT | 40 | 0.009 | 0.012 | 0.016 |
| 1 | WT | 80 | 0.006 | 0.008 | 0.010 |
| 1 | WT | 250 | 0.009 | 0.012 | 0.015 |
| 1 | mut | 5 | 0.002 | 0.003 | 0.004 |
| 1 | mut | 10 | 0.003 | 0.004 | 0.005 |
| 1 | mut | 20 | 0.004 | 0.005 | 0.007 |
| 1 | mut | 40 | 0.007 | 0.009 | 0.011 |
| 1 | mut | 80 | 0.004 | 0.006 | 0.007 |
| 1 | mut | 250 | 0.004 | 0.005 | 0.007 |
| 2 | WT | 5 | 0.002 | 0.003 | 0.004 |
| 2 | WT | 10 | 0.004 | 0.005 | 0.007 |
| 2 | WT | 20 | 0.006 | 0.008 | 0.011 |
| 2 | WT | 40 | 0.008 | 0.012 | 0.015 |
| 2 | WT | 80 | 0.013 | 0.018 | 0.023 |
| 2 | WT | 250 | 0.010 | 0.014 | 0.019 |
| 2 | mut | 5 | 0.002 | 0.003 | 0.004 |
| 2 | mut | 10 | 0.004 | 0.005 | 0.006 |
| 2 | mut | 20 | 0.004 | 0.005 | 0.006 |
| 2 | mut | 40 | 0.004 | 0.005 | 0.007 |
| 2 | mut | 80 | 0.003 | 0.004 | 0.005 |
| 2 | mut | 250 | 0.004 | 0.005 | 0.006 |
| 3 | WT | 5 | 0.002 | 0.003 | 0.004 |
| 3 | WT | 10 | 0.002 | 0.003 | 0.004 |
| 3 | WT | 20 | 0.009 | 0.012 | 0.015 |
| 3 | WT | 40 | 0.007 | 0.010 | 0.012 |
| 3 | WT | 80 | 0.008 | 0.011 | 0.014 |
| 3 | WT | 250 | 0.008 | 0.011 | 0.014 |
| 3 | mut | 5 | 0.003 | 0.004 | 0.005 |
| 3 | mut | 10 | 0.003 | 0.004 | 0.005 |
| 3 | mut | 20 | 0.005 | 0.006 | 0.008 |
| 3 | mut | 40 | 0.005 | 0.006 | 0.008 |
| 3 | mut | 80 | 0.004 | 0.005 | 0.007 |
| 3 | mut | 250 | 0.006 | 0.008 | 0.010 |
| 4 | WT | 5 | 0.004 | 0.006 | 0.007 |
| 4 | WT | 10 | 0.004 | 0.006 | 0.007 |
| 4 | WT | 20 | 0.004 | 0.006 | 0.008 |
| 4 | WT | 40 | 0.005 | 0.007 | 0.009 |
| 4 | WT | 80 | 0.006 | 0.008 | 0.011 |
| 4 | WT | 250 | 0.004 | 0.006 | 0.008 |
| 4 | mut | 5 | 0.003 | 0.003 | 0.004 |
| 4 | mut | 10 | 0.004 | 0.005 | 0.006 |
| 4 | mut | 20 | 0.005 | 0.006 | 0.008 |
| 4 | mut | 40 | 0.005 | 0.007 | 0.009 |
| 4 | mut | 80 | 0.004 | 0.005 | 0.007 |
| 4 | mut | 250 | 0.006 | 0.008 | 0.010 |

TABLE S14. $\tilde{k}_{\text{off}}$ HIV gene

| k | SNP | tet | low.HPD | median | upp.HPD |
|---|---|---|---|---|---|
| 1 | WT | 5 | 0.144 | 0.881 | 2.155 |
| 1 | WT | 10 | 0.084 | 0.562 | 1.685 |
| 1 | WT | 20 | 0.229 | 0.744 | 1.857 |
| 1 | WT | 40 | 0.038 | 0.078 | 0.654 |
| 1 | WT | 80 | 0.015 | 0.031 | 0.060 |
| 1 | WT | 250 | 0.135 | 0.590 | 1.298 |
| 1 | mut | 5 | 0.073 | 0.454 | 1.354 |
| 1 | mut | 10 | 0.051 | 0.368 | 1.166 |
| 1 | mut | 20 | 0.018 | 0.140 | 0.616 |
| 1 | mut | 40 | 0.123 | 0.478 | 1.361 |
| 1 | mut | 80 | 0.014 | 0.024 | 0.042 |
| 1 | mut | 250 | 0.016 | 0.044 | 0.189 |
| 2 | WT | 5 | 0.103 | 0.570 | 1.574 |
| 2 | WT | 10 | 0.052 | 0.401 | 1.406 |
| 2 | WT | 20 | 0.023 | 0.151 | 0.667 |
| 2 | WT | 40 | 0.045 | 0.191 | 0.656 |
| 2 | WT | 80 | 0.047 | 0.204 | 0.605 |
| 2 | WT | 250 | 0.023 | 0.097 | 0.353 |
| 2 | mut | 5 | 0.022 | 0.201 | 0.769 |
| 2 | mut | 10 | 0.013 | 0.037 | 0.169 |
| 2 | mut | 20 | 0.007 | 0.011 | 0.017 |
| 2 | mut | 40 | 0.012 | 0.019 | 0.031 |
| 2 | mut | 80 | 0.009 | 0.014 | 0.021 |
| 2 | mut | 250 | 0.012 | 0.020 | 0.031 |
| 3 | WT | 5 | 0.068 | 0.611 | 2.029 |
| 3 | WT | 10 | 0.038 | 0.556 | 1.761 |
| 3 | WT | 20 | 0.096 | 0.582 | 1.690 |
| 3 | WT | 40 | 0.088 | 0.348 | 0.988 |
| 3 | WT | 80 | 0.037 | 0.374 | 1.164 |
| 3 | WT | 250 | 0.059 | 0.117 | 0.226 |
| 3 | mut | 5 | 0.015 | 0.120 | 0.623 |
| 3 | mut | 10 | 0.011 | 0.030 | 0.233 |
| 3 | mut | 20 | 0.015 | 0.088 | 0.720 |
| 3 | mut | 40 | 0.024 | 0.146 | 0.626 |
| 3 | mut | 80 | 0.010 | 0.017 | 0.026 |
| 3 | mut | 250 | 0.016 | 0.036 | 0.122 |
| 4 | WT | 5 | 0.103 | 0.649 | 1.825 |
| 4 | WT | 10 | 0.020 | 0.195 | 0.755 |
| 4 | WT | 20 | 0.014 | 0.038 | 0.156 |
| 4 | WT | 40 | 0.011 | 0.017 | 0.026 |
| 4 | WT | 80 | 0.017 | 0.036 | 0.091 |
| 4 | WT | 250 | 0.010 | 0.015 | 0.021 |
| 4 | mut | 5 | 0.037 | 0.288 | 0.992 |
| 4 | mut | 10 | 0.010 | 0.020 | 0.041 |
| 4 | mut | 20 | 0.011 | 0.020 | 0.035 |
| 4 | mut | 40 | 0.013 | 0.024 | 0.042 |
| 4 | mut | 80 | 0.007 | 0.010 | 0.014 |
| 4 | mut | 250 | 0.017 | 0.037 | 0.078 |

TABLE S15. $\alpha/k_{\text{off}}$ HIV

| k | SNP | tet | low.HPD | median | upp.HPD |
|---|---|---|---|---|---|
| 1 | WT | 5 | 67.863 | 83.165 | 98.742 |
| 1 | WT | 10 | 110.233 | 128.710 | 148.311 |
| 1 | WT | 20 | 66.024 | 75.388 | 86.230 |
| 1 | WT | 40 | 62.055 | 86.423 | 86.468 |
| 1 | WT | 80 | 125.449 | 163.587 | 197.794 |
| 1 | WT | 250 | 78.058 | 89.165 | 100.338 |
| 1 | mut | 5 | 31.371 | 40.575 | 50.588 |
| 1 | mut | 10 | 39.406 | 48.621 | 58.372 |
| 1 | mut | 20 | 53.780 | 65.401 | 80.227 |
| 1 | mut | 40 | 50.390 | 58.418 | 66.771 |
| 1 | mut | 80 | 102.514 | 128.589 | 150.897 |
| 1 | mut | 250 | 107.239 | 136.430 | 165.491 |
| 2 | WT | 5 | 78.383 | 92.406 | 106.888 |
| 2 | WT | 10 | 85.438 | 97.416 | 111.716 |
| 2 | WT | 20 | 69.807 | 83.687 | 101.449 |
| 2 | WT | 40 | 65.879 | 78.702 | 95.247 |
| 2 | WT | 80 | 49.412 | 61.013 | 75.818 |
| 2 | WT | 250 | 65.047 | 84.639 | 112.098 |
| 2 | mut | 5 | 34.809 | 44.291 | 54.536 |
| 2 | mut | 10 | 38.469 | 51.197 | 64.148 |
| 2 | mut | 20 | 82.526 | 102.341 | 124.937 |
| 2 | mut | 40 | 98.967 | 120.615 | 143.267 |
| 2 | mut | 80 | 152.628 | 182.466 | 213.368 |
| 2 | mut | 250 | 135.805 | 164.695 | 193.826 |
| 3 | WT | 5 | 95.847 | 112.554 | 131.718 |
| 3 | WT | 10 | 152.746 | 177.896 | 201.429 |
| 3 | WT | 20 | 50.485 | 58.166 | 66.866 |
| 3 | WT | 40 | 80.785 | 93.509 | 106.082 |
| 3 | WT | 80 | 87.801 | 99.460 | 133.676 |
| 3 | WT | 250 | 88.618 | 103.986 | 120.259 |
| 3 | mut | 5 | 28.936 | 36.907 | 43.947 |
| 3 | mut | 10 | 46.523 | 63.475 | 79.732 |
| 3 | mut | 20 | 44.259 | 56.383 | 76.276 |
| 3 | mut | 40 | 67.822 | 80.040 | 95.357 |
| 3 | mut | 80 | 114.493 | 141.460 | 169.534 |
| 3 | mut | 250 | 77.043 | 104.075 | 130.875 |
| 4 | WT | 5 | 50.970 | 59.826 | 68.217 |
| 4 | WT | 10 | 77.752 | 95.427 | 118.147 |
| 4 | WT | 20 | 99.153 | 128.960 | 158.025 |
| 4 | WT | 40 | 147.939 | 179.362 | 211.985 |
| 4 | WT | 80 | 113.452 | 149.386 | 180.708 |
| 4 | WT | 250 | 205.845 | 243.531 | 280.996 |
| 4 | mut | 5 | 33.166 | 41.173 | 49.774 |
| 4 | mut | 10 | 40.803 | 53.581 | 68.012 |
| 4 | mut | 20 | 53.655 | 70.015 | 87.576 |
| 4 | mut | 40 | 71.864 | 90.996 | 110.250 |
| 4 | mut | 80 | 132.897 | 162.495 | 192.626 |
| 4 | mut | 250 | 82.023 | 104.932 | 129.862 |

#### B. Negative binomial distribution

In contrast to the Poisson-beta model, the negative-binomial model directly encodes the ratio $\alpha/k_{\text{off}}$ as a single parameter, which is inferred with rather narrow credible intervals. Parsimony suggests that the negative-binomial model is a reasonable choice for the genes considered here, as it encodes for the most relevant kinetic parameters, viz., the average burst size $\alpha/k_{\text{off}}$ and the burst frequency $k_{\text{on}}$. Fig. S11 shows the parameters estimated from the negative-binomial model. The traces of the posteriors chains from each replicates were combined according to the consensus Monte Carlo procedure (see section S4 B) to obtain a representation of the consensus belief in Fig. S12.

The 90% HPD CIs and medians of the estimated parameters $\kappa$, $\mu_X$, $k_{\text{on}}$, and $\alpha/k_{\text{off}}$ are reported in Tables S16–S19 (HBB gene) and Tables S20–S23 (HIV gene).

TABLE S16. $\kappa$ HBB

| k | SNP | tet | low.HPD | median | upp.HPD |
|---|---|---|---|---|---|
| 1 | WT | 5 | 9.675 | 11.307 | 13.213 |
| 1 | WT | 10 | 22.041 | 25.232 | 28.377 |
| 1 | WT | 20 | 18.950 | 20.813 | 22.843 |
| 1 | WT | 40 | 20.381 | 22.383 | 24.447 |
| 1 | WT | 80 | 25.673 | 27.661 | 29.771 |
| 1 | WT | 250 | 17.300 | 18.830 | 20.649 |
| 1 | mut | 5 | 32.550 | 35.329 | 38.490 |
| 1 | mut | 10 | 16.072 | 18.688 | 21.615 |
| 1 | mut | 40 | 23.375 | 25.503 | 27.566 |
| 1 | mut | 80 | 27.251 | 30.309 | 34.188 |
| 1 | mut | 250 | 16.967 | 18.178 | 19.357 |
| 2 | WT | 5 | 23.578 | 26.572 | 29.961 |
| 2 | WT | 10 | 23.453 | 26.572 | 29.997 |
| 2 | WT | 20 | 21.562 | 23.610 | 25.909 |
| 2 | WT | 40 | 25.687 | 28.089 | 30.456 |
| 2 | WT | 80 | 31.358 | 33.931 | 36.457 |
| 2 | WT | 250 | 24.861 | 26.446 | 28.603 |
| 2 | mut | 5 | 26.626 | 29.697 | 33.019 |
| 2 | mut | 10 | 31.988 | 36.180 | 41.474 |
| 2 | mut | 40 | 40.925 | 44.272 | 47.412 |
| 2 | mut | 80 | 27.407 | 30.832 | 34.251 |
| 2 | mut | 250 | 22.856 | 24.409 | 26.037 |
| 3 | WT | 5 | 25.676 | 29.020 | 32.801 |
| 3 | WT | 10 | 32.162 | 35.936 | 39.914 |
| 3 | WT | 20 | 30.366 | 32.843 | 35.501 |
| 3 | WT | 40 | 30.841 | 33.200 | 35.696 |
| 3 | WT | 80 | 24.637 | 26.419 | 28.015 |
| 3 | WT | 250 | 20.441 | 21.780 | 23.367 |
| 3 | mut | 5 | 24.569 | 27.336 | 30.290 |
| 3 | mut | 10 | 42.637 | 47.605 | 52.771 |
| 3 | mut | 40 | 35.782 | 38.444 | 41.379 |
| 3 | mut | 80 | 31.078 | 35.045 | 38.636 |
| 3 | mut | 250 | 28.041 | 30.084 | 32.136 |
| 4 | WT | 5 | 17.636 | 20.198 | 23.004 |
| 4 | WT | 10 | 17.386 | 19.747 | 22.473 |
| 4 | WT | 20 | 21.240 | 23.109 | 25.149 |
| 4 | WT | 40 | 29.894 | 32.568 | 35.174 |
| 4 | WT | 80 | 29.411 | 31.375 | 33.856 |
| 4 | WT | 250 | 23.425 | 25.036 | 26.811 |
| 4 | mut | 5 | 14.236 | 15.954 | 17.716 |
| 4 | mut | 10 | 22.721 | 26.243 | 30.138 |
| 4 | mut | 40 | 32.983 | 35.546 | 38.257 |
| 4 | mut | 80 | 28.575 | 31.849 | 35.621 |
| 4 | mut | 250 | 26.358 | 27.852 | 29.440 |

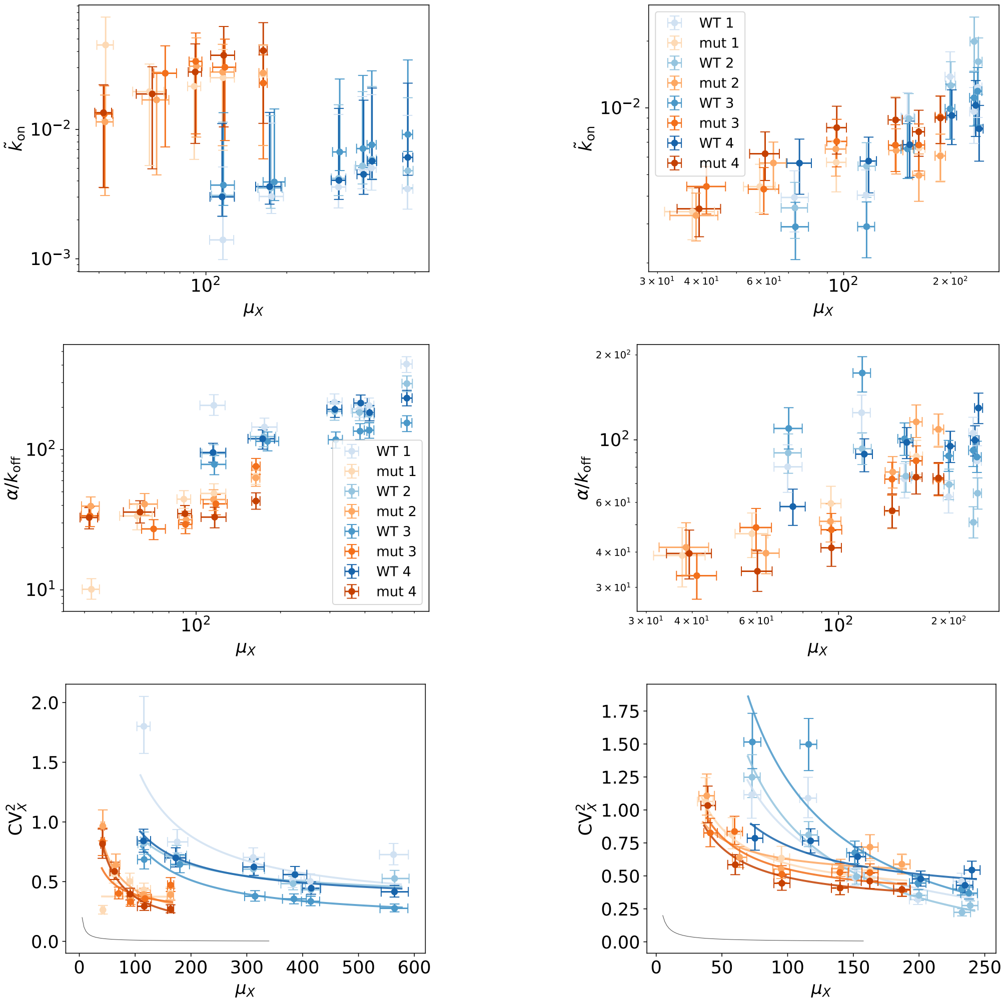


**A** HBB. **B** HIV.

FIG. S11. Estimates for parameters $\tilde{k}_{on}$, $\mu_X$, and $\alpha/k_{off}$ of the negative-binomial model, from wild-type (blue) and mutant (orange) cell data, for all induction levels, shades of colors corresponds to replicates. Points are medians, error bars comprise 90% HPD CIs. HBB-gene results show consistent results across all the replicates (panel **A**). The HIV-gene results are reported in panels **B**. Results are consistent with the Poisson-beta model estimates (Fig. S9). Increasing expression levels, three replicates show a drop-off in the average burst size and an increase in the burst frequency, see also Fig. S2.

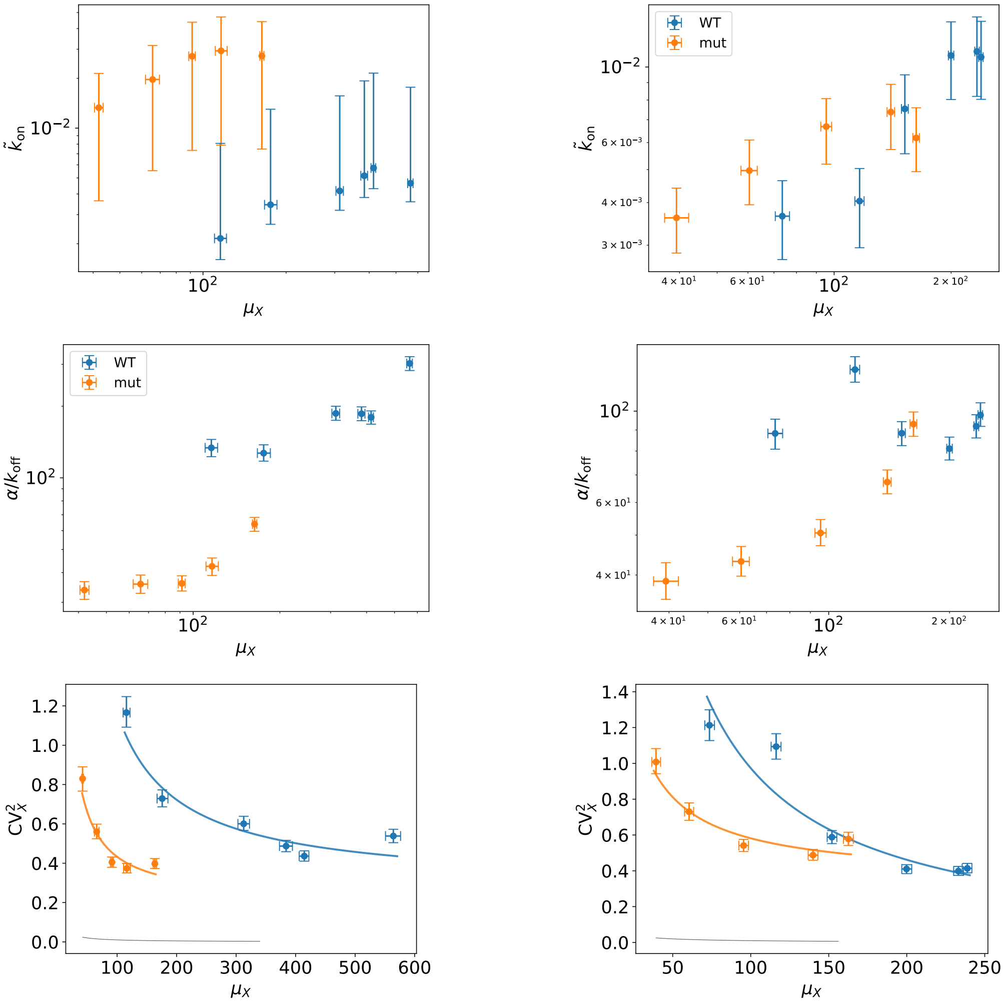


**A** HBB. **B** HIV.

FIG. S12. Consensus estimates of the parameters $\tilde{k}_{\mathrm{on}}$, $\mu_X$, and $\alpha/k_{\mathrm{off}}$ from the negative-binomial model, from wild-type (blue) and mutant (orange) cell data, for all induction levels. Points are medians, error bars comprise 90% HPD CIs. HBB-gene results are in panel **A**, HIV-gene results are reported in panel **B**.

TABLE S17. $\mu_X$ HBB

| k | SNP | tet | low.HPD | median | upp.HPD |
|---|---|---|---|---|---|
| 1 | WT | 5 | 103.193 | 115.687 | 126.875 |
| 1 | WT | 10 | 157.904 | 175.269 | 194.327 |
| 1 | WT | 20 | 293.749 | 311.584 | 331.666 |
| 1 | WT | 40 | 357.077 | 382.068 | 402.882 |
| 1 | WT | 80 | 397.085 | 413.625 | 430.396 |
| 1 | WT | 250 | 537.710 | 562.996 | 589.504 |
| 1 | mut | 5 | 39.255 | 42.325 | 45.219 |
| 1 | mut | 10 | 53.368 | 61.634 | 69.735 |
| 1 | mut | 40 | 85.470 | 90.459 | 95.633 |
| 1 | mut | 80 | 103.857 | 115.778 | 127.620 |
| 1 | mut | 250 | 157.575 | 163.136 | 168.842 |
| 2 | WT | 5 | 104.884 | 115.671 | 127.339 |
| 2 | WT | 10 | 156.119 | 174.189 | 192.233 |
| 2 | WT | 20 | 292.999 | 311.784 | 331.770 |
| 2 | WT | 40 | 360.835 | 382.742 | 404.849 |
| 2 | WT | 80 | 397.159 | 414.805 | 430.893 |
| 2 | WT | 250 | 542.191 | 564.530 | 590.432 |
| 2 | mut | 5 | 39.031 | 42.105 | 45.158 |
| 2 | mut | 10 | 57.835 | 65.531 | 72.791 |
| 2 | mut | 40 | 86.987 | 92.086 | 96.961 |
| 2 | mut | 80 | 103.723 | 115.390 | 127.323 |
| 2 | mut | 250 | 157.508 | 163.312 | 168.777 |
| 3 | WT | 5 | 104.159 | 116.177 | 127.657 |
| 3 | WT | 10 | 163.259 | 179.811 | 197.030 |
| 3 | WT | 20 | 296.072 | 314.846 | 332.531 |
| 3 | WT | 40 | 363.753 | 384.298 | 406.525 |
| 3 | WT | 80 | 397.951 | 414.531 | 430.464 |
| 3 | WT | 250 | 538.464 | 564.609 | 588.360 |
| 3 | mut | 5 | 38.955 | 41.838 | 44.995 |
| 3 | mut | 10 | 63.927 | 70.558 | 77.724 |
| 3 | mut | 40 | 86.541 | 91.645 | 96.597 |
| 3 | mut | 80 | 105.958 | 117.686 | 129.102 |
| 3 | mut | 250 | 157.977 | 163.500 | 169.119 |
| 4 | WT | 5 | 103.541 | 114.909 | 127.623 |
| 4 | WT | 10 | 153.130 | 172.692 | 190.326 |
| 4 | WT | 20 | 292.635 | 312.606 | 330.812 |
| 4 | WT | 40 | 365.063 | 386.377 | 407.528 |
| 4 | WT | 80 | 399.614 | 415.485 | 432.185 |
| 4 | WT | 250 | 540.558 | 564.417 | 589.218 |
| 4 | mut | 5 | 38.556 | 41.557 | 44.534 |
| 4 | mut | 10 | 55.032 | 63.112 | 71.160 |
| 4 | mut | 40 | 85.960 | 91.252 | 96.281 |
| 4 | mut | 80 | 104.157 | 116.484 | 127.851 |
| 4 | mut | 250 | 157.935 | 163.479 | 169.027 |

TABLE S18. $\tilde{k}_{\mathrm{on}}$ HBB

| k | SNP | tet | low.HPD | median | upp.HPD |
|---|---|---|---|---|---|
| 1 | WT | 5 | 0.001 | 0.001 | 0.005 |
| 1 | WT | 10 | 0.002 | 0.003 | 0.011 |
| 1 | WT | 20 | 0.003 | 0.004 | 0.013 |
| 1 | WT | 40 | 0.004 | 0.005 | 0.018 |
| 1 | WT | 80 | 0.004 | 0.005 | 0.019 |
| 1 | WT | 250 | 0.002 | 0.003 | 0.013 |
| 1 | mut | 5 | 0.012 | 0.045 | 0.074 |
| 1 | mut | 10 | 0.005 | 0.019 | 0.032 |
| 1 | mut | 40 | 0.006 | 0.021 | 0.035 |
| 1 | mut | 80 | 0.007 | 0.025 | 0.041 |
| 1 | mut | 250 | 0.007 | 0.027 | 0.044 |
| 2 | WT | 5 | 0.002 | 0.003 | 0.011 |
| 2 | WT | 10 | 0.003 | 0.004 | 0.013 |
| 2 | WT | 20 | 0.003 | 0.004 | 0.015 |
| 2 | WT | 40 | 0.004 | 0.005 | 0.019 |
| 2 | WT | 80 | 0.004 | 0.006 | 0.022 |
| 2 | WT | 250 | 0.003 | 0.005 | 0.018 |
| 2 | mut | 5 | 0.003 | 0.011 | 0.019 |
| 2 | mut | 10 | 0.005 | 0.017 | 0.027 |
| 2 | mut | 40 | 0.008 | 0.031 | 0.051 |
| 2 | mut | 80 | 0.007 | 0.028 | 0.045 |
| 2 | mut | 250 | 0.007 | 0.028 | 0.046 |
| 3 | WT | 5 | 0.003 | 0.004 | 0.014 |
| 3 | WT | 10 | 0.003 | 0.004 | 0.015 |
| 3 | WT | 20 | 0.005 | 0.007 | 0.025 |
| 3 | WT | 40 | 0.005 | 0.007 | 0.026 |
| 3 | WT | 80 | 0.005 | 0.008 | 0.027 |
| 3 | WT | 250 | 0.006 | 0.009 | 0.034 |
| 3 | mut | 5 | 0.003 | 0.013 | 0.021 |
| 3 | mut | 10 | 0.008 | 0.027 | 0.045 |
| 3 | mut | 40 | 0.009 | 0.033 | 0.054 |
| 3 | mut | 80 | 0.009 | 0.030 | 0.050 |
| 3 | mut | 250 | 0.006 | 0.023 | 0.038 |
| 4 | WT | 5 | 0.002 | 0.003 | 0.011 |
| 4 | WT | 10 | 0.003 | 0.004 | 0.013 |
| 4 | WT | 20 | 0.003 | 0.004 | 0.015 |
| 4 | WT | 40 | 0.003 | 0.005 | 0.017 |
| 4 | WT | 80 | 0.004 | 0.006 | 0.021 |
| 4 | WT | 250 | 0.004 | 0.006 | 0.022 |
| 4 | mut | 5 | 0.003 | 0.013 | 0.023 |
| 4 | mut | 10 | 0.005 | 0.019 | 0.031 |
| 4 | mut | 40 | 0.008 | 0.028 | 0.045 |
| 4 | mut | 80 | 0.010 | 0.037 | 0.063 |
| 4 | mut | 250 | 0.011 | 0.041 | 0.065 |

TABLE S19. $\alpha/k_{\text{on}}$ HBB

| k | SNP | tet | low.HPD | median | upp.HPD |
|---|---|---|---|---|---|
| 1 | WT | 5 | 175.298 | 206.600 | 246.085 |
| 1 | WT | 10 | 120.820 | 144.691 | 167.516 |
| 1 | WT | 20 | 190.560 | 217.633 | 249.246 |
| 1 | WT | 40 | 170.125 | 194.471 | 222.404 |
| 1 | WT | 80 | 181.291 | 206.076 | 232.972 |
| 1 | WT | 250 | 354.322 | 406.867 | 459.164 |
| 1 | mut | 5 | 8.579 | 10.098 | 12.020 |
| 1 | mut | 10 | 26.797 | 33.634 | 40.524 |
| 1 | mut | 40 | 38.121 | 44.326 | 50.962 |
| 1 | mut | 80 | 41.023 | 48.685 | 57.037 |
| 1 | mut | 250 | 56.503 | 64.628 | 73.724 |
| 2 | WT | 5 | 79.762 | 93.656 | 108.321 |
| 2 | WT | 10 | 104.729 | 123.788 | 143.569 |
| 2 | WT | 20 | 161.758 | 186.721 | 211.985 |
| 2 | WT | 40 | 160.712 | 183.674 | 208.313 |
| 2 | WT | 80 | 156.316 | 177.567 | 201.871 |
| 2 | WT | 250 | 258.811 | 295.198 | 334.765 |
| 2 | mut | 5 | 33.600 | 39.522 | 45.973 |
| 2 | mut | 10 | 34.079 | 40.868 | 48.606 |
| 2 | mut | 40 | 27.354 | 31.636 | 36.378 |
| 2 | mut | 80 | 36.745 | 44.057 | 51.733 |
| 2 | mut | 250 | 54.699 | 62.845 | 71.400 |
| 3 | WT | 5 | 66.198 | 78.414 | 90.605 |
| 3 | WT | 10 | 97.887 | 114.499 | 133.520 |
| 3 | WT | 20 | 102.297 | 117.574 | 133.503 |
| 3 | WT | 40 | 117.655 | 135.267 | 152.972 |
| 3 | WT | 80 | 120.396 | 137.453 | 155.178 |
| 3 | WT | 250 | 134.325 | 154.819 | 174.205 |
| 3 | mut | 5 | 28.241 | 33.741 | 39.190 |
| 3 | mut | 10 | 22.808 | 27.192 | 31.662 |
| 3 | mut | 40 | 25.141 | 29.293 | 33.804 |
| 3 | mut | 80 | 34.292 | 41.021 | 47.862 |
| 3 | mut | 250 | 66.549 | 75.912 | 86.434 |
| 4 | WT | 5 | 81.257 | 95.302 | 110.891 |
| 4 | WT | 10 | 100.651 | 119.384 | 138.147 |
| 4 | WT | 20 | 169.562 | 193.318 | 219.519 |
| 4 | WT | 40 | 187.460 | 214.568 | 244.764 |
| 4 | WT | 80 | 160.411 | 183.310 | 206.170 |
| 4 | WT | 250 | 204.957 | 232.671 | 264.311 |
| 4 | mut | 5 | 27.219 | 32.786 | 38.579 |
| 4 | mut | 10 | 29.960 | 35.848 | 43.176 |
| 4 | mut | 40 | 30.133 | 34.940 | 39.916 |
| 4 | mut | 80 | 27.685 | 33.007 | 38.815 |
| 4 | mut | 250 | 37.564 | 42.970 | 49.161 |

TABLE S20. $\kappa$ HIV

| k | SNP | tet | low.HPD | median | upp.HPD |
|---|---|---|---|---|---|
| 1 | WT | 5 | 8.683 | 9.925 | 11.355 |
| 1 | WT | 10 | 10.830 | 12.201 | 13.502 |
| 1 | WT | 20 | 16.946 | 18.291 | 19.696 |
| 1 | WT | 40 | 17.995 | 19.136 | 20.300 |
| 1 | WT | 80 | 16.814 | 17.998 | 19.187 |
| 1 | WT | 250 | 24.456 | 25.898 | 27.413 |
| 1 | mut | 5 | 16.450 | 19.977 | 23.853 |
| 1 | mut | 10 | 20.826 | 23.864 | 27.041 |
| 1 | mut | 20 | 22.626 | 24.912 | 27.281 |
| 1 | mut | 40 | 19.162 | 20.677 | 22.160 |
| 1 | mut | 80 | 26.548 | 28.642 | 30.929 |
| 1 | mut | 250 | 24.056 | 26.022 | 27.857 |
| 2 | WT | 5 | 15.406 | 17.501 | 19.880 |
| 2 | WT | 10 | 25.483 | 28.210 | 30.905 |
| 2 | WT | 20 | 36.121 | 38.563 | 41.320 |
| 2 | WT | 40 | 33.654 | 35.716 | 37.949 |
| 2 | WT | 80 | 30.499 | 32.001 | 33.633 |
| 2 | WT | 250 | 37.389 | 39.279 | 41.400 |
| 2 | mut | 5 | 21.886 | 25.819 | 30.735 |
| 2 | mut | 10 | 42.808 | 47.476 | 52.389 |
| 2 | mut | 20 | 21.292 | 23.474 | 25.662 |
| 2 | mut | 40 | 30.748 | 33.257 | 35.874 |
| 2 | mut | 80 | 19.858 | 21.585 | 23.433 |
| 2 | mut | 250 | 16.597 | 17.858 | 19.331 |
| 3 | WT | 5 | 10.305 | 11.911 | 13.704 |
| 3 | WT | 10 | 9.759 | 10.994 | 12.394 |
| 3 | WT | 20 | 15.478 | 16.882 | 18.327 |
| 3 | WT | 40 | 15.642 | 16.744 | 17.818 |
| 3 | WT | 80 | 16.849 | 17.904 | 19.029 |
| 3 | WT | 250 | 19.059 | 20.252 | 21.472 |
| 3 | mut | 5 | 33.159 | 38.139 | 43.447 |
| 3 | mut | 10 | 28.905 | 32.666 | 36.895 |
| 3 | mut | 20 | 25.677 | 28.163 | 30.685 |
| 3 | mut | 40 | 21.877 | 23.641 | 25.428 |
| 3 | mut | 80 | 22.105 | 23.864 | 25.581 |
| 3 | mut | 250 | 16.736 | 17.863 | 19.027 |
| 4 | WT | 5 | 34.330 | 38.261 | 42.130 |
| 4 | WT | 10 | 37.272 | 40.394 | 44.002 |
| 4 | WT | 20 | 43.020 | 46.210 | 50.115 |
| 4 | WT | 40 | 48.515 | 51.534 | 54.970 |
| 4 | WT | 80 | 54.224 | 57.441 | 60.923 |
| 4 | WT | 250 | 54.376 | 57.496 | 61.232 |
| 4 | mut | 5 | 25.398 | 29.929 | 34.621 |
| 4 | mut | 10 | 34.261 | 38.357 | 42.825 |
| 4 | mut | 20 | 28.323 | 31.007 | 33.655 |
| 4 | mut | 40 | 26.348 | 28.262 | 30.242 |
| 4 | mut | 80 | 21.116 | 22.690 | 24.363 |
| 4 | mut | 250 | 20.337 | 21.627 | 23.114 |

TABLE S21. $\mu_X$ HIV

| k | SNP | tet | low.HPD | median | upp.HPD |
|---|---|---|---|---|---|
| 1 | WT | 5 | 66.749 | 72.714 | 79.573 |
| 1 | WT | 10 | 109.215 | 115.559 | 121.995 |
| 1 | WT | 20 | 144.952 | 151.119 | 157.583 |
| 1 | WT | 40 | 192.656 | 199.155 | 205.249 |
| 1 | WT | 80 | 225.999 | 232.337 | 238.487 |
| 1 | WT | 250 | 232.122 | 238.168 | 244.560 |
| 1 | mut | 5 | 31.397 | 37.570 | 43.353 |
| 1 | mut | 10 | 52.305 | 58.144 | 64.224 |
| 1 | mut | 20 | 89.505 | 95.336 | 101.545 |
| 1 | mut | 40 | 133.432 | 139.663 | 146.087 |
| 1 | mut | 80 | 156.547 | 162.818 | 169.259 |
| 1 | mut | 250 | 180.854 | 187.190 | 193.263 |
| 2 | WT | 5 | 66.749 | 72.966 | 79.146 |
| 2 | WT | 10 | 109.748 | 115.981 | 122.719 |
| 2 | WT | 20 | 145.901 | 152.450 | 158.287 |
| 2 | WT | 40 | 193.656 | 199.980 | 206.311 |
| 2 | WT | 80 | 226.495 | 232.703 | 239.197 |
| 2 | WT | 250 | 232.607 | 238.913 | 244.831 |
| 2 | mut | 5 | 32.501 | 38.487 | 44.321 |
| 2 | mut | 10 | 58.006 | 63.477 | 68.941 |
| 2 | mut | 20 | 88.766 | 95.055 | 101.497 |
| 2 | mut | 40 | 133.898 | 140.390 | 146.663 |
| 2 | mut | 80 | 156.238 | 162.667 | 168.835 |
| 2 | mut | 250 | 180.541 | 186.876 | 193.151 |
| 3 | WT | 5 | 66.720 | 73.187 | 79.646 |
| 3 | WT | 10 | 109.464 | 116.001 | 122.177 |
| 3 | WT | 20 | 144.933 | 151.139 | 157.537 |
| 3 | WT | 40 | 192.762 | 199.160 | 205.284 |
| 3 | WT | 80 | 225.811 | 232.428 | 238.252 |
| 3 | WT | 250 | 231.709 | 237.918 | 244.421 |
| 3 | mut | 5 | 36.242 | 41.159 | 46.565 |
| 3 | mut | 10 | 53.945 | 59.653 | 65.693 |
| 3 | mut | 20 | 89.646 | 95.617 | 102.024 |
| 3 | mut | 40 | 133.566 | 139.776 | 146.113 |
| 3 | mut | 80 | 156.555 | 162.557 | 168.787 |
| 3 | mut | 250 | 180.705 | 186.932 | 193.327 |
| 4 | WT | 5 | 69.298 | 75.131 | 81.160 |
| 4 | WT | 10 | 111.288 | 117.514 | 123.639 |
| 4 | WT | 20 | 146.956 | 153.289 | 159.350 |
| 4 | WT | 40 | 194.908 | 201.289 | 207.621 |
| 4 | WT | 80 | 228.382 | 234.962 | 240.972 |
| 4 | WT | 250 | 234.384 | 240.198 | 246.468 |
| 4 | mut | 5 | 34.044 | 39.263 | 45.106 |
| 4 | mut | 10 | 54.421 | 60.168 | 65.966 |
| 4 | mut | 20 | 89.558 | 95.636 | 101.791 |
| 4 | mut | 40 | 133.322 | 139.929 | 146.100 |
| 4 | mut | 80 | 156.024 | 162.534 | 168.737 |
| 4 | mut | 250 | 180.686 | 187.024 | 193.207 |

TABLE S22. $\tilde{k}_{on}$ HIV

| k | SNP | tet | low.HPD | median | upp.HPD |
|---|---|---|---|---|---|
| 1 | WT | 5 | 0.003 | 0.004 | 0.005 |
| 1 | WT | 10 | 0.003 | 0.004 | 0.005 |
| 1 | WT | 20 | 0.007 | 0.009 | 0.012 |
| 1 | WT | 40 | 0.010 | 0.014 | 0.018 |
| 1 | WT | 80 | 0.007 | 0.010 | 0.012 |
| 1 | WT | 250 | 0.009 | 0.012 | 0.016 |
| 1 | mut | 5 | 0.003 | 0.003 | 0.004 |
| 1 | mut | 10 | 0.003 | 0.004 | 0.006 |
| 1 | mut | 20 | 0.004 | 0.006 | 0.007 |
| 1 | mut | 40 | 0.007 | 0.009 | 0.011 |
| 1 | mut | 80 | 0.005 | 0.007 | 0.008 |
| 1 | mut | 250 | 0.005 | 0.006 | 0.008 |
| 2 | WT | 5 | 0.003 | 0.004 | 0.005 |
| 2 | WT | 10 | 0.004 | 0.005 | 0.007 |
| 2 | WT | 20 | 0.007 | 0.009 | 0.012 |
| 2 | WT | 40 | 0.009 | 0.013 | 0.016 |
| 2 | WT | 80 | 0.015 | 0.020 | 0.026 |
| 2 | WT | 250 | 0.011 | 0.016 | 0.021 |
| 2 | mut | 5 | 0.002 | 0.003 | 0.004 |
| 2 | mut | 10 | 0.004 | 0.006 | 0.007 |
| 2 | mut | 20 | 0.005 | 0.007 | 0.008 |
| 2 | mut | 40 | 0.005 | 0.006 | 0.008 |
| 2 | mut | 80 | 0.004 | 0.005 | 0.006 |
| 2 | mut | 250 | 0.005 | 0.006 | 0.008 |
| 3 | WT | 5 | 0.002 | 0.003 | 0.004 |
| 3 | WT | 10 | 0.002 | 0.003 | 0.004 |
| 3 | WT | 20 | 0.005 | 0.007 | 0.008 |
| 3 | WT | 40 | 0.007 | 0.010 | 0.013 |
| 3 | WT | 80 | 0.008 | 0.011 | 0.014 |
| 3 | WT | 250 | 0.009 | 0.012 | 0.016 |
| 3 | mut | 5 | 0.003 | 0.004 | 0.006 |
| 3 | mut | 10 | 0.003 | 0.004 | 0.005 |
| 3 | mut | 20 | 0.005 | 0.007 | 0.009 |
| 3 | mut | 40 | 0.005 | 0.007 | 0.009 |
| 3 | mut | 80 | 0.005 | 0.007 | 0.009 |
| 3 | mut | 250 | 0.007 | 0.009 | 0.011 |
| 4 | WT | 5 | 0.004 | 0.006 | 0.007 |
| 4 | WT | 10 | 0.004 | 0.006 | 0.007 |
| 4 | WT | 20 | 0.005 | 0.007 | 0.009 |
| 4 | WT | 40 | 0.007 | 0.009 | 0.012 |
| 4 | WT | 80 | 0.007 | 0.010 | 0.013 |
| 4 | WT | 250 | 0.006 | 0.008 | 0.010 |
| 4 | mut | 5 | 0.003 | 0.004 | 0.004 |
| 4 | mut | 10 | 0.005 | 0.006 | 0.008 |
| 4 | mut | 20 | 0.006 | 0.008 | 0.010 |
| 4 | mut | 40 | 0.007 | 0.009 | 0.011 |
| 4 | mut | 80 | 0.006 | 0.008 | 0.010 |
| 4 | mut | 250 | 0.007 | 0.009 | 0.011 |

TABLE S23. $\alpha/k_{\rm on}$ HIV

| k | SNP | tet | low.HPD | median | upp.HPD |
|---|---|---|---|---|---|
| 1 | WT | 5 | 65.123 | 80.224 | 96.055 |
| 1 | WT | 10 | 105.447 | 124.804 | 144.056 |
| 1 | WT | 20 | 62.196 | 72.693 | 82.359 |
| 1 | WT | 40 | 55.029 | 62.855 | 71.783 |
| 1 | WT | 80 | 91.968 | 105.675 | 120.244 |
| 1 | WT | 250 | 73.419 | 84.125 | 95.301 |
| 1 | mut | 5 | 30.097 | 38.894 | 48.835 |
| 1 | mut | 10 | 38.247 | 46.466 | 55.041 |
| 1 | mut | 20 | 50.656 | 59.480 | 68.448 |
| 1 | mut | 40 | 48.290 | 55.669 | 63.965 |
| 1 | mut | 80 | 75.487 | 87.423 | 99.612 |
| 1 | mut | 250 | 95.084 | 108.486 | 123.429 |
| 2 | WT | 5 | 76.105 | 89.902 | 104.946 |
| 2 | WT | 10 | 81.737 | 93.017 | 105.950 |
| 2 | WT | 20 | 65.181 | 74.363 | 83.731 |
| 2 | WT | 40 | 61.362 | 69.436 | 78.737 |
| 2 | WT | 80 | 44.884 | 51.005 | 57.922 |
| 2 | WT | 250 | 56.796 | 64.669 | 73.245 |
| 2 | mut | 5 | 33.159 | 41.584 | 50.823 |
| 2 | mut | 10 | 33.516 | 39.641 | 45.896 |
| 2 | mut | 20 | 43.426 | 51.414 | 59.328 |
| 2 | mut | 40 | 67.491 | 76.908 | 87.220 |
| 2 | mut | 80 | 101.936 | 115.771 | 132.951 |
| 2 | mut | 250 | 94.257 | 109.003 | 123.450 |
| 3 | WT | 5 | 92.776 | 109.735 | 130.017 |
| 3 | WT | 10 | 147.773 | 172.554 | 197.158 |
| 3 | WT | 20 | 88.084 | 100.818 | 114.809 |
| 3 | WT | 40 | 77.288 | 87.747 | 99.551 |
| 3 | WT | 80 | 80.400 | 91.796 | 103.421 |
| 3 | WT | 250 | 76.617 | 86.925 | 98.877 |
| 3 | mut | 5 | 27.222 | 32.985 | 39.572 |
| 3 | mut | 10 | 40.746 | 48.876 | 57.111 |
| 3 | mut | 20 | 41.247 | 48.004 | 54.827 |
| 3 | mut | 40 | 63.647 | 72.555 | 83.277 |
| 3 | mut | 80 | 72.747 | 84.254 | 95.213 |
| 3 | mut | 250 | 63.329 | 72.322 | 82.431 |
| 4 | WT | 5 | 49.732 | 58.001 | 66.871 |
| 4 | WT | 10 | 77.064 | 88.972 | 100.832 |
| 4 | WT | 20 | 85.802 | 97.901 | 110.523 |
| 4 | WT | 40 | 83.077 | 94.987 | 107.236 |
| 4 | WT | 80 | 87.612 | 99.737 | 112.167 |
| 4 | WT | 250 | 114.176 | 129.687 | 146.438 |
| 4 | mut | 5 | 32.124 | 39.552 | 47.867 |
| 4 | mut | 10 | 29.015 | 34.206 | 40.406 |
| 4 | mut | 20 | 35.600 | 41.423 | 47.690 |
| 4 | mut | 40 | 48.680 | 56.041 | 64.317 |
| 4 | mut | 80 | 64.269 | 73.738 | 84.134 |
| 4 | mut | 250 | 63.794 | 73.262 | 83.058 |

TABLE S24. $\kappa$ HBB

| k | SNP | tet | low.HPD | median | upp.HPD |
|---|---|---|---|---|---|
| 2 | WT | 5 | 260.821 | 267.314 | 273.981 |
| 2 | WT | 10 | 270.677 | 277.100 | 284.186 |
| 2 | WT | 20 | 298.204 | 304.307 | 311.348 |
| 2 | WT | 40 | 305.802 | 312.765 | 319.762 |
| 2 | WT | 80 | 321.045 | 327.146 | 334.050 |
| 2 | WT | 250 | 338.221 | 345.124 | 349.512 |
| 2 | mut | 5 | 165.515 | 172.460 | 179.105 |
| 2 | mut | 10 | 192.112 | 198.697 | 206.055 |
| 2 | mut | 40 | 183.042 | 189.718 | 196.109 |
| 2 | mut | 80 | 185.722 | 192.343 | 199.121 |
| 2 | mut | 250 | 178.993 | 185.852 | 192.476 |

TABLE S25. $\mu_X$ HBB

| k | SNP | tet | low.HPD | median | upp.HPD |
|---|---|---|---|---|---|
| 2 | WT | 5 | 11.310 | 11.728 | 12.148 |
| 2 | WT | 10 | 16.261 | 16.814 | 17.355 |
| 2 | WT | 20 | 23.646 | 24.285 | 25.005 |
| 2 | WT | 40 | 33.593 | 34.477 | 35.487 |
| 2 | WT | 80 | 42.511 | 43.591 | 44.589 |
| 2 | WT | 250 | 42.719 | 43.602 | 44.549 |
| 2 | mut | 5 | 7.300 | 7.704 | 8.128 |
| 2 | mut | 10 | 11.689 | 12.224 | 12.753 |
| 2 | mut | 40 | 21.350 | 22.256 | 23.111 |
| 2 | mut | 80 | 17.868 | 18.619 | 19.384 |
| 2 | mut | 250 | 21.277 | 22.220 | 23.121 |

### C. Poisson distribution

The Poisson model encodes only one biological parameter, viz., the average gene expression level $\mu_X$. We fitted this model to data from one of the replicates as a benchmark. The 90% HPD CIs and medians of the estimated parameters $\kappa$ and $\mu_X$, are reported in Tables S24–S25 (HBB gene) and Tables S26–S27 (HIV gene). It is worth noting that, compared to the prior derived in section S3 and both the estimates from the Poisson-beta and negative-binomial models of Tables S6, S11, S16, and S20, the $\kappa$ is overestimated. In fact, high values of $\kappa$ compensate for the small dispersion encoded in a Poisson random variable. Jointly with the fact that the Poisson model shows lower GoF than the two general models (subsection S4 C and figure S7), we conclude that the expression of the genes HIV and HBB is relative to a Poisson random variable and a flexible gene expression model for $X_i^{(k)}$, such as the Poisson-beta or the negative-binomial models, is necessary to exploit the measurement equation (24).

TABLE S26. $\mathrm{CV}_X^2$ HBB

| k | SNP | tet | low.HPD | median | upp.HPD |
|---|---|---|---|---|---|
| 2 | WT | 5 | 0.082 | 0.085 | 0.088 |
| 2 | WT | 10 | 0.058 | 0.059 | 0.061 |
| 2 | WT | 20 | 0.040 | 0.041 | 0.042 |
| 2 | WT | 40 | 0.028 | 0.029 | 0.030 |
| 2 | WT | 80 | 0.022 | 0.023 | 0.024 |
| 2 | WT | 250 | 0.022 | 0.023 | 0.023 |
| 2 | mut | 5 | 0.123 | 0.130 | 0.137 |
| 2 | mut | 10 | 0.078 | 0.082 | 0.085 |
| 2 | mut | 40 | 0.043 | 0.045 | 0.047 |
| 2 | mut | 80 | 0.052 | 0.054 | 0.056 |
| 2 | mut | 250 | 0.043 | 0.045 | 0.047 |

TABLE S27. $\kappa$ HIV

| k | SNP | tet | low.HPD | median | upp.HPD |
|---|---|---|---|---|---|
| 4 | WT | 5 | 245.093 | 251.782 | 258.632 |
| 4 | WT | 10 | 276.948 | 283.448 | 290.234 |
| 4 | WT | 20 | 291.102 | 297.690 | 304.497 |
| 4 | WT | 40 | 285.262 | 291.524 | 298.090 |
| 4 | WT | 80 | 300.678 | 306.854 | 313.563 |
| 4 | WT | 250 | 315.404 | 321.897 | 328.357 |
| 4 | mut | 5 | 172.173 | 179.215 | 186.314 |
| 4 | mut | 10 | 183.339 | 189.696 | 196.491 |
| 4 | mut | 20 | 178.477 | 185.104 | 191.728 |
| 4 | mut | 40 | 189.708 | 196.348 | 203.244 |
| 4 | mut | 80 | 183.321 | 189.823 | 196.490 |
| 4 | mut | 250 | 185.416 | 192.345 | 198.760 |

TABLE S28. $\mu_X$ HIV

| k | SNP | tet | low.HPD | median | upp.HPD |
|---|---|---|---|---|---|
| 4 | WT | 5 | 11.520 | 11.960 | 12.434 |
| 4 | WT | 10 | 17.091 | 17.645 | 18.199 |
| 4 | WT | 20 | 24.572 | 25.315 | 26.012 |
| 4 | WT | 40 | 37.049 | 38.005 | 39.049 |
| 4 | WT | 80 | 46.374 | 47.601 | 48.746 |
| 4 | WT | 250 | 45.807 | 46.889 | 48.064 |
| 4 | mut | 5 | 6.517 | 6.877 | 7.264 |
| 4 | mut | 10 | 12.027 | 12.547 | 13.114 |
| 4 | mut | 20 | 15.649 | 16.369 | 17.065 |
| 4 | mut | 40 | 19.839 | 20.665 | 21.484 |
| 4 | mut | 80 | 19.157 | 19.943 | 20.770 |
| 4 | mut | 250 | 20.907 | 21.769 | 22.651 |

TABLE S29. $CV_X^2$ HIV

| k | SNP | tet | low.HPD | median | upp.HPD |
|---|---|---|---|---|---|
| 4 | WT | 5 | 0.080 | 0.084 | 0.087 |
| 4 | WT | 10 | 0.055 | 0.057 | 0.058 |
| 4 | WT | 20 | 0.038 | 0.040 | 0.041 |
| 4 | WT | 40 | 0.026 | 0.026 | 0.027 |
| 4 | WT | 80 | 0.021 | 0.021 | 0.022 |
| 4 | WT | 250 | 0.021 | 0.021 | 0.022 |
| 4 | mut | 5 | 0.137 | 0.145 | 0.153 |
| 4 | mut | 10 | 0.076 | 0.080 | 0.083 |
| 4 | mut | 20 | 0.059 | 0.061 | 0.064 |
| 4 | mut | 40 | 0.047 | 0.048 | 0.050 |
| 4 | mut | 80 | 0.048 | 0.050 | 0.052 |
| 4 | mut | 250 | 0.044 | 0.046 | 0.048 |

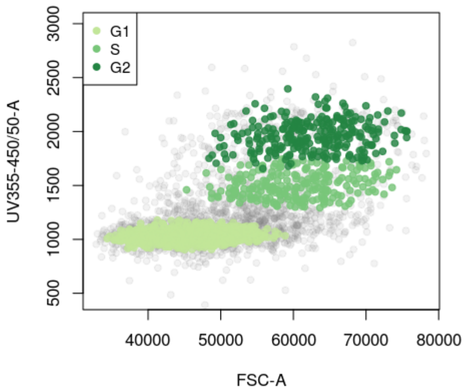


FIG. S13. Scatter plot of the `UV355-450/50-A` vs `FSC-A` signal for the 40 ng/mL Tet-induced HBB gene, replicate $k = 3$. Cells from the three phases, highlighted with different green-scale colors, were separated using flowClust.

### S7. CELL CYCLE

Staining for DNA concentration allows us to heuristically find cells that are in G1, S, and G2 phases of the cell cycle, see Figs. S1(right) and S13. We considered the dataset with cells treated at concentration of 40 ng/mL of Tet. We separated the data points corresponding to the G1 phase from those from S and G2 using flowClust [4]. The less dense cluster S-G2 was further separated in two groups (corresponding to the phases S and G2) running the same algorithm again. Results are shown in Fig. S13 for HBB cell line, $k = 3$.

Data from phase G1, S, and G2 are referred to as $y^{(k)}_{\mathrm{G1}}$, $y^{(k)}_{\mathrm{S}}$, and $y^{(k)}_{\mathrm{G2}}$, respectively, for each replicate $k$. We refer to their averages (sample standard deviations of mean) as $\bar{y}_{\mathrm{G1}}$, $\bar{y}_{\mathrm{S}}$, and $\bar{y}_{\mathrm{G2}}$, ($s_{\bar{y}_{\mathrm{G1}}}$, $s_{\bar{y}_{\mathrm{S}}}$, and $s_{\bar{y}_{\mathrm{G2}}}$) respectively. To take into account that the mean gene expression seems to change with the cell phase, we introduce the conversion factors $c^{(k)} = \bar{x}/\bar{y}^{(k)}$ to obtain $\bar{x}^{(k)}_{\mathrm{i}} = c^{(k)}\bar{y}_{\mathrm{i}}$ and $s^{(k)}_{\bar{x}_{\mathrm{i}}} = c^{(k)}s_{\bar{y}_{\mathrm{i}}}$ which in turn are used in the informative priors

$$\mu^{(k)}_{X_{\mathrm{i}}} \sim \mathcal{N}(\bar{x}^{(k)}_{\mathrm{i}}, s^{(k)}_{\bar{x}_{\mathrm{i}}}), \tag{40}$$

$\mathrm{i} = \mathrm{G1}, \mathrm{S}, \mathrm{G2}$. Equations (40) take the place of $\mu_X$ in the DAG of Fig. S5(B) for the G1, S, and G2 phase, respectively, for each replicate $k$. Fitting the negative-binomial model to 500 samples from each dataset yields the consensus estimates of Fig. 4(C-D) (*Main text*). In addition to this, we also subset reads form each phase into three groups by their size (based on the values of their FSC-A fluorescense signal, see *Main text*). We assume that the cell cycle and the cell size are extrinsic contributors to the total transgene mRNA variability, with the remaining variability sources thought of as being intrinsic. As in Ref. [8], using the symbol $\langle\cdot\rangle_{\mathrm{I}}$ for the average over the intrinsic variables, with the cell phase held fixed, and $\langle\cdot\rangle_{\mathrm{E}}$ for the average over the different cell phases, the law of total variance allows us to write, for the mRNA abundance $X$,

$$\mathrm{CV}^2_X = \frac{\langle\langle X^2\rangle_{\mathrm{I}} - \langle X\rangle^2_{\mathrm{I}}\rangle_{\mathrm{E}}}{\langle\langle X\rangle_{\mathrm{I}}\rangle^2_{\mathrm{E}}} + \frac{\langle\langle X\rangle^2_{\mathrm{I}}\rangle_{\mathrm{E}} - \langle\langle X\rangle_{\mathrm{I}}\rangle^2_{\mathrm{E}}}{\langle\langle X\rangle_{\mathrm{I}}\rangle^2_{\mathrm{E}}}, \tag{41}$$

where the first term on the r.h.s. is the intrinsic noise, while the second term is the extrinsic noise. Computing the two terms gives the intrinsic and extrinsic noise levels of Tables S30 and Fig. 4(D) (*Main text*), which show that the cell cycle and the cell size always contributed only a minor term to the total noise.

TABLE S30. Intrinsic noise, extrinsic noise and total noise from each replicate of wild-type HBB and HIV genes, 40 ng/mL Tet. First table is based on cell-phase only partition, second table is bases on both cell phase and cell size. Extrinsic noise has the lowest contribution to the total noise.

| gene | k | intr. noise | extr. noise | tot. noise |
|---|---|---|---|---|
| HBB | 1 | 0.587 | 0.017 | 0.060 |
| HBB | 2 | 0.399 | 0.033 | 0.014 |
| HBB | 3 | 0.380 | 0.023 | 0.008 |
| HBB | 4 | 0.448 | 0.024 | 0.030 |
| HIV | 1 | 0.322 | 0.053 | 0.015 |
| HIV | 2 | 0.417 | 0.041 | 0.020 |
| HIV | 3 | 0.444 | 0.040 | 0.029 |
| HIV | 4 | 0.289 | 0.036 | 0.021 |

| gene | k | intr. noise | extr. noise | tot. noise |
|---|---|---|---|---|
| HBB | 1 | 0.514 | 0.025 | 0.539 |
| HBB | 2 | 0.467 | 0.054 | 0.521 |
| HBB | 3 | 0.381 | 0.020 | 0.401 |
| HBB | 4 | 0.477 | 0.023 | 0.500 |
| HIV | 1 | 0.324 | 0.044 | 0.369 |
| HIV | 2 | 0.423 | 0.039 | 0.461 |
| HIV | 3 | 0.492 | 0.039 | 0.531 |
| HIV | 4 | 0.301 | 0.029 | 0.330 |

### S8. POLII-MEDIATED 3'-5' INTERACTIONS BY CHIA-PET

We considered scRNAseq and ChIA-PET data accessible at the GEO Series numbers GSE124682 [23] and GSE33664 [24], respectively.

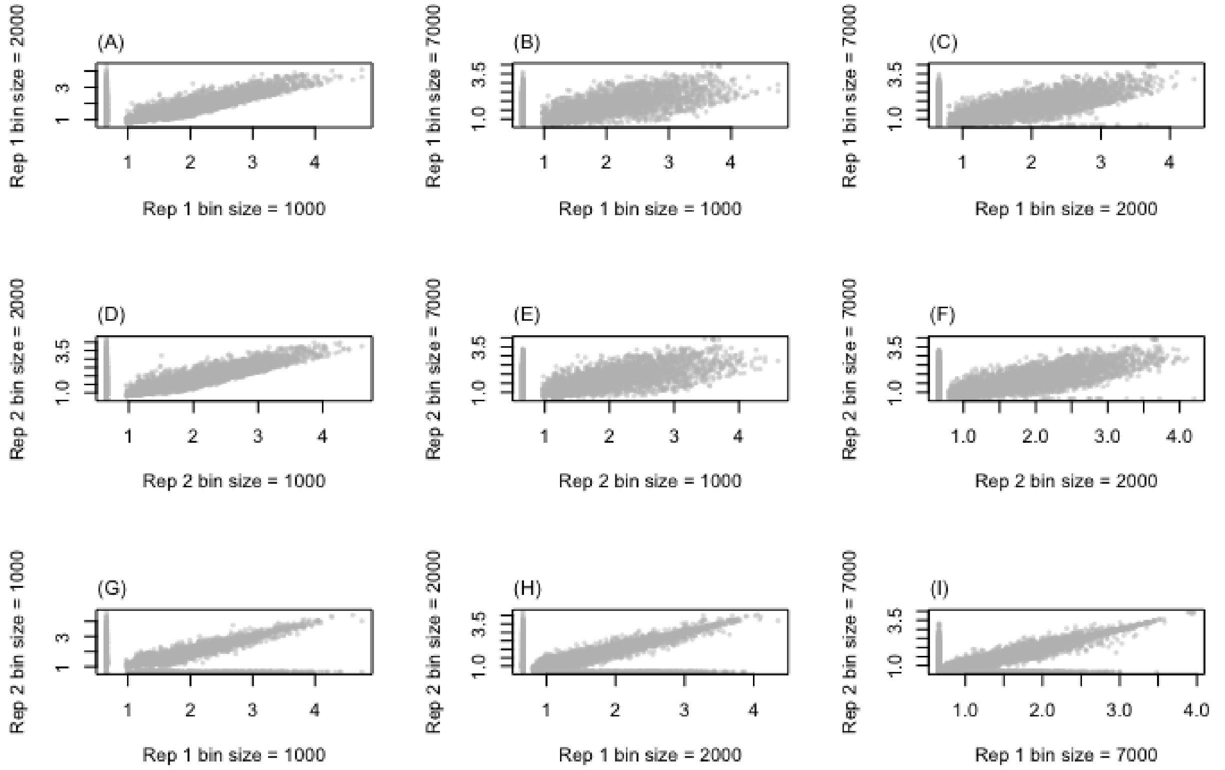


FIG. S14. Joint plots of 3'-5' interaction scores. For each biological replicate (Rep 1 and Rep 2, corresponding to GSM832464 and GSM832465, respectively), the interaction scores obtained at different bin resolutions (1, 2, and 7 Kbs) appear strongly correlated. (G)-(I) Similarly, the two biological repeats appear strongly correlated at each bin resolution.

Raw chromatin contact frequency is highest for small genomic distances. In order to normalise for this and expose deviations from this general relationship, we need to divide by the expected number of reads at a given genomic distance. We calculate this by random sampling 10000 genomic intervals and measuring the contact frequency over this sample. This estimate appears robust to decreasing the number of sampled intervals. The relation between gene length and the normalised 3'-5' interaction score is illustrated in Fig. S15. Genes with length smaller than the resolution of the interaction matrices are discarded from the main analysis.

We fitted the model of equations (54)–(56) to the smRNAseq data, thus enabling a convenient classification of genes based on transcription; see also, e.g., reference [25]. Without correction for incomplete capture of mRNA, the parameter $\bar{k}_{\text{on}}$ incorporates here both biological and technical above-Poisson noise, whilst allowing the ranking of the genes based on their total noise.

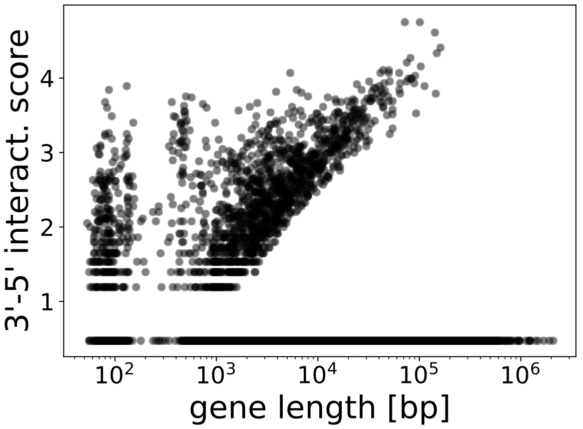


FIG. S15. 3'-5' interaction score vs gene length. The cluster corresponding to small lengths mostly includes pseudo-genes.

### S9. MICROSCOPIC GENE EXPRESSION MODEL

We referred to the models of section S2 as the phenomelogical models. In fact, our main concern there was to exploit a minimal description of the statistics of the transcription events and the stationary mRNA distribution—which is the (observed) phenomenology, indeed. Due to their simplicity, these models allowed us to attain the important goal of separating the technical noise (due to background fluorescence and measurement process) from the biological noise encoded into $X_i$.

Nevertheless, there are specific microscopic biological mechanisms, more difficult to observe, that may give rise to the observed phenomena. In our tetracycline-inducible genes, Tet repressor (TetR) homodimers bind to the operator $\text{TetO}_2$ downstream of the transcription start site (TSS). When such a binding event occurs, the transcription is inhibited as the elongation is impeded. Adding Tet in turn alters the conformation of TetR and hinders the binding events, having the net effect of inducing the gene expression. Crucially, during the "on" phase, the transcription rate is proportional to the abundance of PolII (law of mass action), which can be thought of as waiting in a compartment upstream of the TSS [26]. Therefore, when the gene is actively transcribing, its rate can vary in time according to the amount of PolII ready to initiate transcription. After transcription, PolII can either be re-injected into the compartment and set ready for a new initiation event (PolII recycling), or disposed into the nuclear environment. Also, the compartment recruits PolII from the nuclear environment. This can be described by means of the following chemical reaction scheme:

$$\varnothing \xrightarrow{\gamma} \text{PolII}, \tag{42}$$

$$\text{DNA}_{\text{on}} + \text{PolII} \xrightarrow{l\,\beta} \text{mRNA} + \text{DNA}_{\text{on}} + \text{PolII}, \tag{43}$$

$$\text{DNA}_{\text{on}} + \text{PolII} \xrightarrow{(1-l)\beta} \text{mRNA} + \text{DNA}_{\text{on}}, \tag{44}$$

$$\text{DNA}_{\text{on}} \xrightarrow{\lambda_{\text{off}}} \text{DNA}_{\text{off}}, \tag{45}$$

$$\text{DNA}_{\text{off}} \xrightarrow{\lambda_{\text{on}}} \text{DNA}_{\text{on}}, \tag{46}$$

$$\text{mRNA} \xrightarrow{d} \varnothing, \tag{47}$$

$$\text{PolII} \xrightarrow{\delta} \varnothing, \tag{48}$$

where $\text{DNA}_{\text{on}}$ and $\text{DNA}_{\text{off}}$ are unlocked and locked DNA configurations, respectively. The presence of the 3'-5' crosstalk loop is thought to facilitate the recycling of PolII after each transcription event; therefore we can study the effect of the recycling on the simulated expression data by tuning $l$ in the reaction scheme. Obviously, the pA mutation lowers the recycling probability $l$ with respect to the WT, but $l$ is not supposed to be zero in mutant genes, as the recycling can occur by means of other mechanisms (e.g., diffusion). By the law of mass action

$$\lambda_{\text{off}} = nK_\lambda, \tag{49}$$

$$\lambda_{\text{on}} = K_\lambda, \tag{50}$$

where $K_\lambda$ is a chemical affinity and $n$ is the concentration of TetR. Hence, we can imitate variations in the Tet dose by fine-tuning $n$, with large values of Tet (high induction levels) corresponding to small values of $n$. Unlike the simpler phenomenological models, we do not have an analytical likelihood for this model, thus parameter inference is more challenging, to be addressed with likelihood-free methods. We simulate the model using the Doob-Gillespie algorithm; sample trajectories of mRNA abundances are plotted in Fig. 5 D (*Main text*).

When all the chemical species are highly abundant and the gene is always in "on" state (this can be achieved in the limit as $k_{\text{off}} \to 0$), it is straightforward to derive the following rate equations,

$$\frac{\mathrm{d}}{\mathrm{d}t}[\text{PolII}] = \alpha - [\text{PolII}](\delta - \beta(1-l)), \tag{51}$$

$$\frac{\mathrm{d}}{\mathrm{d}t}[\text{mRNA}] = [\text{PolII}]\beta - [\text{mRNA}]d, \tag{52}$$

where $[X]$ is the abundance of the species $X$. The stationary mRNA abundance is then

$$[\text{mRNA}] = \frac{\beta}{d}\frac{\alpha}{\delta + \beta(1-l)}, \tag{53}$$

which corresponds to the vertical lines of Fig. 4 B (*Main text*). While the parameters $\gamma$, $\beta$, $d$, $\delta$, $K_\lambda$ are chosen to simulate mRNA abundances and noises in ranges consistent with those of the real data, fine-tuning the recycling probability and the induction parameters $l$ and $n$ yields patterns similar to those observed in the experimental setting (i.e., those of Figs. S9, S11, and 2 (*Main text*)). More specifically, a simple scatter plot of the sample averages versus the $\text{CV}^2$ of [mRNA] shows a drift of the noise curve from the Poisson case $\text{CV}^2_X = 1/\mu_X$ as the recycling rate $l$ increases. Fitting a negative binomial (NB) Bayesian model

$$\text{mRNA} \sim \text{NB}(\mu_X, k_{\text{on}}), \tag{54}$$

$$\mu_X \sim \text{Gamma}(0.001, 0.001), \tag{55}$$

$$k_{\text{on}} \sim \text{Gamma}(0.001, 0.001), \tag{56}$$

to 500 simulated stationary mRNA abundances, allowed us to estimate the average burst size $\alpha/k_{\text{off}} = \mu_X/k_{\text{on}}$ and the burst frequency $k_{\text{on}}$ shown in Figs. S16 and 5 C (*Main text*).

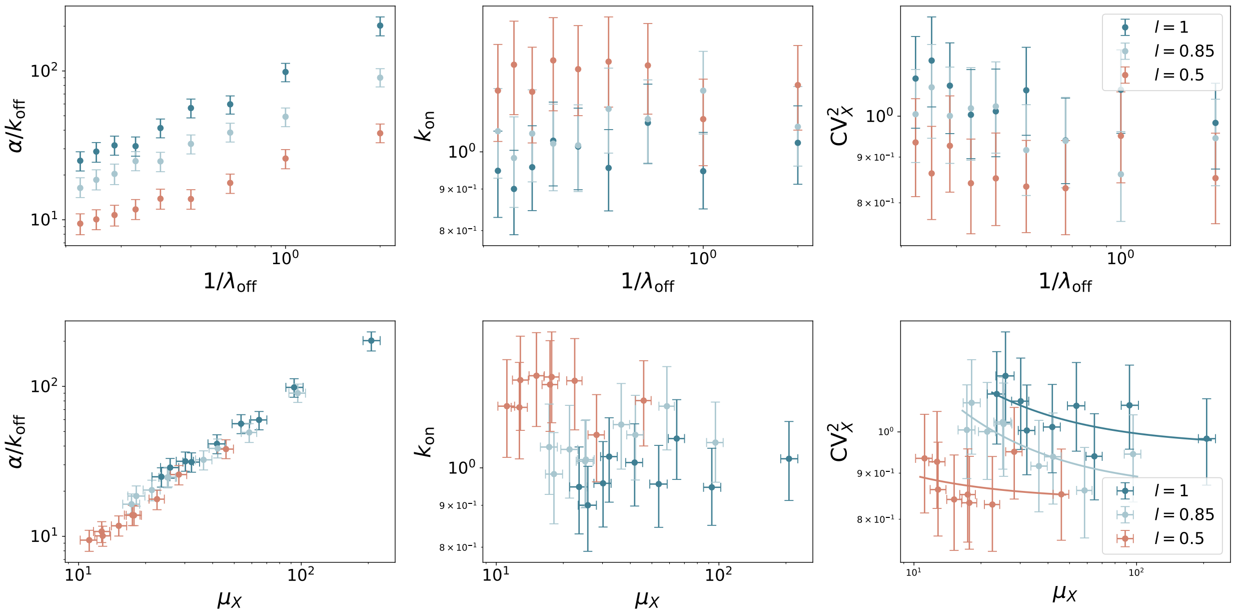


FIG. S16. Negative-binomial model fit to 500 mRNA abundances simulated from the microscopic model. The pattern of the inferred average burst sizes and burst frequencies mirrors those obtained from the real data. For each value of the recycling probability $l$ ($l = 0.5, 0.85, 1$), simulations are performed with $\lambda_{\rm off} = 0.5, 1, 1.5, 2, 2.5, 3, 2.5, 3.5, 4, 4.5$; remaining parameters are $(\gamma, \beta, d, \delta, \lambda_{\rm on}) = (10, 10, 0.01, 1, 0.01)$. The solid lines in the noise plot (lower-left plot) are fitted $\mathrm{CV}_X^2 = A/\mu_X + B$ curves.

### S10. MATERIALS

#### A. Cell lines and cell culture

The wildtype HBB and HIV-1-*env* cell lines have been utilized in previous studies [22, 27, 28]. The nomenclature was changed for the present manuscript, with the cell lines denoted HBB WT, HBB mut, HIV WT and HIV mut, which had been denoted $\beta$ pA+, $\beta$ pA−, HIV-1 pA+ and HIV-1 pA− in [22], respectively. Cells were maintained in DMEM medium supplemented with 10% fetal bovine serum and 100 $\mu$g mL$^{-1}$ penicillin-streptomycin (DMEM-10). Induction of cell lines was carried out for 16 hours before downstream experiments.

**Deletion cell line construction.** The design and construction of the deletion cell line used the protocol detailed in [29], with the following changes. A dual sgRNA strategy was employed with a 5' guide binding between the AmpR promoter and CMV enhancer and a 3' guide binding just after the 3' FRT site. The use of plasmid pSpCas9(BB)-2A-GFP (PX458) and dual targeting necessitated the transfection with two plasmids, each containing a respective guide. Transfection was carried out using calcium phosphate, followed by washing the cells with warm PBS after 16-24 hours and replacing the media. Cells were allowed to recover for 48-72 hours before single-cells were isolated in 96-well plates via FACS (BD ARIAFusion), with the brightest 10% GFP positive cells being sorted. Testing for deletion was initially verified via genomic DNA extraction and PCR, followed by smFISH assay using flow cytometry.

#### B. Single-molecule RNA fluorescence in situ hybridization

**Probe sets.** Probe sets for HBB and HIV-1-*env* RNA were designed with the tool at www.biosearchtech.com/stellarisdesigner (see Table S31 for sequences). The probe sets were synthesized by LGC Biosearch Technologies as custom Stellaris® probe sets. AKT1 probes were ready-made and ordered from LGC Biosearch.

**smFISH.** smFISH staining followed the probe manufacturer's protocol. Briefly, cells were grown on poly-L-lysine treated glass coverslips overnight, fixed in 3.7% formaldehyde for 10 minutes and permeabilized in ethanol for > 1 hour. After overnight staining at 37°C in dextran sulphate and formamide buffered with SSC, cells were washed, followed by mounting onto a slide using Vectashield with DAPI as the mounting medium. Imageing was carried out

TABLE S31. Sequences and details of smFISH probe sets. Product Name: Stellaris® FISH Probes, Custom Assay with Quasar® 670 Dye.

1) Oligo Name: HIV

```
tcactaaacgagctcgtcga ggtcaaaacagcgtggatgg
taaacgctagagtccggagg gagctcggtaccaagcttaa
agaattccaccacactggac cagcagttgttgcagaatta
ctatgtcgacacccaattct tctgtcgagtaacgcctatt
agtctaggatctactggagg tggtacaagcagttttaggc
ggcaatgaaagcaacacttt cctaaggcttttgtcatgaa
gagtctgactgttctgatga gctgctttgatagagaagct
cttcttcttctattccttcg aggatccgttcactaatcga
cagatcgtcccagataagtg tagctgaagaggcacaggct
agagtaagtctctcaagcgg ttccacaatcctcgttacaa
aatatttgagggcttcccac ccaatactgtaggagattcc
gcactattctttagttcctg ctgtggcattgagcaagtta
cttctataaccctatctgtc agctctataagctgcttgta
tattcttctaggtatgtggc agcaaaatcctttccaagcc
tactttttgaccacttgcca ttacagcaggccatccaatc
ctcagctcgtctcattcttt cctctagactcgagatactg
gctgatcagcgggtttaaac ctggcaactagaaggcacag
accttccagggtcaaggaag taggaaaggacagtgggagt
```

2) Oligo Name: HBB

```
tcactaaacgagctcgtcga aaacagcgtggatggcgtct
cggtgtcttctatggaggtc tttaaacgctagagtccgga
gtcagaagcaaatgtaagct ggttgctagtgaacacagtt
tgcaccatggtgtctgtttg gcagtaacggcagacttctc
caacttcatccacgttcacc aaagaacctctgggtccaag
gagtggacagatccccaaag cttagggttgcccataacag
gagcactttcttgccatgag caggccatcactaaaggcac
cttgaggttgtccaggtgag cactcagtgtggcaaaggtg
acgtgcagcttgtcacagtg agcctgaagttctcaggatc
aaagtgatgggccagcacac ctggtggggtgaattctttg
caccactttctgataggcag cgcttagtgatacttgtggg
tggacagcaagaaagcgagc cttagggaacaaaggaacct
tagacccagtttggtagttg tcatgttttctacagctaga
tccagcagacatgggtgatc tcctcatgttttctacagtc
ctagacagcagacatgggtg gtgatcctcatgttttctac
tacagtcgtccagcagacat atgggtgatcctcatgtttt
ttctacagctagacagcaga agacatgggtgatcctcatg
ctcatgttttctacagtcgt tagacagcagacatgggtga
ttatctagatccggtggatc ttgtggtttgtccaaactca
gcatttttttcactgcattc tgcagcttataatggttaca
gcaattgttgttgttaactt
```

on a brightfield microscope.

**Flow cytometry.** DNA staining was carried out with FxCycle™ Violet Stain (ThermoFisher, F10347) at a concentration of 1 $\mu$g mL$^{-1}$. Fixed cells were analysed on a BD Fortessa. Processing and data analysis of raw flow cytometry data was carried out using the flowcore R package [1] (v1.48, R version 3.3).

**SmFISH spot counting.** Quantification of RNA was carried out using FISH-quant [30]. Images were imported with the following settings: XY 64.8 nm; Z 200 nm; Refractive index 1.515; NA 1.40; Em 592; Ex 546; Microscope widefield. Cell outlines were drawn manually. A single image was then processed and the settings used to batch-process the remaining set. The *threshold* and *quality score* parameters of FISH-quant were set to quantify as many spots as possible while reducing spurious detection through batch-specific selection of these parameters.

### C. RNA isolation and preparation, and degradation rate estimation

Total RNA was extracted from the respective cell lines following the RNeasy Mini Kit (Qiagen, 47104) protocol, using QIAshredder (Qiagen, 79654). RNA for RNA-seq analysis was treated with TURBO DNA-*free*™ kit (ThermoFisher, AM1907).

To estimate the mRNA degradation rate, RNA was reverse transcribed using random primers (Promega, C118A) and M-MLV reverse transcriptase (Promega, M170A) followed by qPCR using SensiMix™ SYBR® No-Rox (Bioline, QT650-02) on a Qiagen Rotor-Gene Q. Gene-specific primers were used (HIV Forward `TCTCCTACGGCAGGAAGAAG`; HIV Reverse `GGTAGCTGAAGAGGCACAGG`). Analysis was carried out by calculating the CT values using the qpcR R package [31] (v1.4-1) and from this $2^{-\Delta\Delta \mathrm{Ct}}$ were calculated using the mut time 0 concentration as the reference sample. A degradation time series was carried out by standard induction method at 250 ng mL$^{-1}$ tetracycline for 16 hours, followed by removal of media and washing with warm DMEM-10. Cells were then placed in fresh medium and samples were taken at different time points following on from this.

### D. Nanostring

Cells were seeded, induced and processed as indicated previously (subsection S10 A), with the following alteration: after trypsinisation cells were resuspended in 1 mL of PBS and kept on ice. Counting of cells was carried out via Countess (ThermoFisher) cell counter with 100 $\mu$L (50 : 50) PBS to trypan blue. Samples were spun down at 500 g for 5 minutes and were then resuspended in RLT buffer from RNeasy Mini Kit (Qiagen, 47104) with beta-mercaptoethanol to obtain a concentration of 6500 cells per $\mu$L. Samples were then vortexed for 1 minute and placed at $-80$°C. Cell lysis was verified under a microscope. Samples were shipped on dry ice to an external provider for processing. Custom probe sets, including probes targeting HIV-1-*env* along with GAPDH and AKT1 as house-keeping genes, were designed and shipped by NanoString Technologies.

### E. RNA-seq

**Library preparation.** RNA-seq libraries were prepared using 500 ng of total input RNA and the NEBNext® UltraTM II Directional RNA Library Prep Kit for Illumina (E7760L), along with the NEBNext® rRNA Depletion Kit (E6310L) and NEBNext® Multiplex Oligos for Illumina Set 1 and 2 (E7335, E7500). Ribo-depletion was carried out to capture transgene RNA regardless of the absence or presence of a poly(A) tail. The manufacturer's manual was followed, with the final PCR amplification using 9 cycles. Libraries were assessed via Bioanalyser, diluted and mixed before being sequenced on an Illumina® NextSeq 500, generating paired-end reads with read length 42.

**RNA-seq analysis.** Data quality control was performed with FastQC v0.11.5. Read and adapter trimming was carried out using TrimGalore! v0.4.3 with cutadapt v0.4.3 using default settings [32]. Indices for STAR to map to were constructed from the human genome (GRCh38.p12, Gencode primary annotation) and the respective (HBB WT/mut and HIV-1-*env* WT/mut) transgenic sequence. The GTF was modified to include these genes as a separate chromosome (chrHBB or chrHIV). To mask the existing HBB sequence, bedtools' (v2.25.0) `maskfasta` command was used [33]. RNA-seq reads were mapped to the genome using STAR software v2.5.3a with parameter `--outSAMattributes XS` [34]. Counts per gene were calculated using LiBiNorm [35] acting in an HTSeq-count [36] compatible mode with the following parameters: `--format=bam --minaqual=10 --stranded=reverse --mode=intersection-strict`. Coverage statistics were generated using deepTools' (v3.1.3) `bamCoverage` [37]. Fold changes for the HBB and HIV genes were calculated using DESeq2 v1.22.1 [38] from Bioconductor release 3.8 and R v3.5.1.

**Splicing analysis.** To analyse potential alternative splicing in the *env* transgene, BAM files were imported into R (v4.0.2) and analysed using SGSeq (v1.4) [39]. A table of counts relating to potential splice variants (Table S32) indicates that there are potential alternative splicing events, however the number of reads related to each variant indicates that they are present across all samples and appear related to the overall abundance of mRNA. To further analyse the difference between introns and exons in the HIV-1 env wild-type and mutant cell lines, the Transcripts Per Kilobase Million (TPM), a normalisation that takes account of both sequence depth and gene length, were calculated (Fig. S17). BAM files were imported into R (v4.0.2) and analysed using Rsubread (v2.2.6) [40], calling `FeatureCounts` with the following parameters: `minOverlap=20`, `isPairedEnd=TRUE`, `strandSpecific=2`. We do see a slightly higher fraction of intronic reads present in the *env* mutant at 250 ng $mL^{-1}$ Tet, although the principal difference between the mutant and wild-type appears to be overall mRNA abundance. In addition to this, ref. [22] quantified the levels of HBB pre-mRNA relative to the total HBB RNA and determined that ratios are the same within the first 2 hours of induction (splicing was slightly effected after 24 hours, albeit this phenotype appeared to arise subsequent to RNAPII depletion).

TABLE S32. Counts of potential splice variants in *env* transgene.

| variantID | mut 0 | mut 0 | mut 0 | mut 250 | mut 250 | mut 250 | WT 0 | WT 0 | WT 0 | WT 250 | WT 250 | WT 250 |
|---|---|---|---|---|---|---|---|---|---|---|---|---|
| 1 | 0 | 0 | 0 | 0 | 0 | 0 | 0 | 0 | 0 | 0 | 1 | 0 |
| 2 | 28 | 21 | 14 | 233 | 164 | 223 | 42 | 46 | 44 | 404 | 434 | 303 |
| 3 | 28 | 24 | 27 | 46 | 51 | 58 | 31 | 27 | 45 | 33 | 28 | 45 |
| 4 | 0 | 0 | 0 | 0 | 0 | 0 | 0 | 0 | 0 | 0 | 0 | 1 |
| 5 | 26 | 30 | 22 | 42 | 51 | 59 | 36 | 33 | 44 | 32 | 23 | 48 |
| 6 | 0 | 0 | 0 | 0 | 0 | 1 | 0 | 0 | 0 | 0 | 1 | 0 |
| 7 | 41 | 53 | 31 | 41 | 56 | 65 | 49 | 46 | 60 | 36 | 30 | 54 |
| 8 | 0 | 0 | 0 | 0 | 0 | 0 | 0 | 0 | 0 | 0 | 1 | 0 |
| 9 | 0 | 0 | 0 | 1 | 0 | 1 | 0 | 0 | 0 | 1 | 4 | 1 |
| 10 | 0 | 0 | 0 | 0 | 1 | 0 | 0 | 0 | 0 | 0 | 0 | 0 |

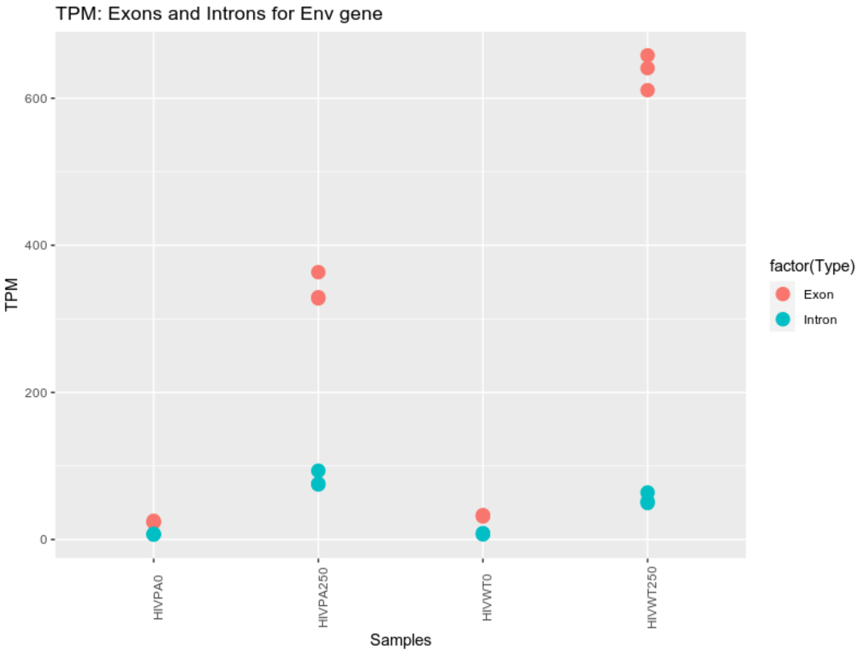


FIG. S17. TPMs of exonic and intronic regions in *env* transgene.

[1] F. Hahne, N. LeMeur, R. R. Brinkman, B. Ellis, P. Haaland, D. Sarkar, J. Spidlen, E. Strain, and R. Gentleman, BMC Bioinformatics **10**, 106 (2009).
[2] H. M. Shapiro, *Practical Flow Cytometry* (John Wiley & Sons, Inc., Hoboken, NJ, USA, 2003) pp. 1–733.
[3] K. Lo, R. Brinkman, and R. Gottardo, Cytometry A (2008).
[4] K. Lo, F. Hahne, R. Brinkman, and R. Gottardo, BMC Bioinformatics (2009), R package version 3.5.0.
[5] B. Munsky, G. Neuert, and A. van Oudenaarden, Science **336**, 183 (2012).
[6] J. Kim and J. C. Marioni, Genome Biology **14**, R7 (2013).
[7] S. Tiberi, M. Walsh, M. Cavallaro, D. Hebenstreit, and B. Finkenstädt, Bioinformatics **34**, i647 (2018).
[8] M. B. Elowitz, A. J. Levine, E. D. Siggia, and P. S. Swain, Science **297**, 1183 (2002).
[9] J. R. S. Newman, S. Ghaemmaghami, J. Ihmels, D. K. Breslow, M. Noble, J. L. DeRisi, and J. S. Weissman, Nature **441**, 840 (2006).
[10] A. Bar-Even, J. Paulsson, N. Maheshri, M. Carmi, E. O'Shea, Y. Pilpel, and N. Barkai, Nature Genetics **38**, 636 (2006).
[11] R. D. Dar, B. S. Razooky, L. S. Weinberger, C. D. Cox, and M. L. Simpson, PLoS ONE **10**, 1 (2015).
[12] M. Soltani, C. A. Vargas-Garcia, D. Antunes, and A. Singh, PLoS Computational Biology **12**, e1004972 (2016).
[13] M. Dobrzynski and F. J. Bruggeman, Proceedings of the National Academy of Sciences **106**, 2583 (2009).
[14] E. Meinelt, M. Reunanen, M. Edinger, M. Jaimes, A. Stall, D. Sasaki, and J. Trotter, *Standardizing Application Setup Across Multiple Flow Cytometers Using BD FACSDiva$^{TM}$ Version 6 Software* (2012).
[15] C. Zechner, J. Ruess, P. Krenn, S. Pelet, M. Peter, J. Lygeros, and H. Koeppl, Proceedings of the National Academy of Sciences **109**, 8340 (2012).
[16] D. Adler, "vioplot: Violin plot," (2005), https://cran.r-project.org/package=vioplot.
[17] D. J. Spiegelhalter, K. R. K. R. Abrams, and J. P. Myles, *Bayesian approaches to clinical trials and health-care evaluation* (John Wiley & Sons, 2004) p. 391.
[18] A. Patil, D. Huard, and C. Fonnesbeck, Journal of Statistical Software **35**, 1 (2010).
[19] H. Haario, E. Saksman, and J. Tamminen, Bernoulli **7**, 223 (2001).
[20] S. L. Scott, A. W. Blocker, F. V. Bonassi, H. A. Chipman, E. I. George, and R. E. McCulloch, International Journal of Management Science and Engineering Management **11**, 78 (2016).
[21] O. Tange, ;login: The USENIX Magazine **36**, 42 (2011).
[22] C. K. Mapendano, S. Lykke-Andersen, J. Kjems, E. Bertrand, and T. H. Jensen, Molecular Cell **40**, 410 (2010).
[23] A. M. Klein, L. Mazutis, I. Akartuna, N. Tallapragada, A. Veres, V. Li, L. Peshkin, D. A. Weitz, and M. W. Kirschner, Cell **161**, 1187 (2015).
[24] G. Li, X. Ruan, R. K. Auerbach, K. S. Sandhu, M. Zheng, P. Wang, H. M. Poh, Y. Goh, J. Lim, J. Zhang, H. S. Sim, S. Q. Peh, F. H. Mulawadi, C. T. Ong, Y. L. Orlov, S. Hong, Z. Zhang, S. Landt, D. Raha, G. Euskirchen, C.-L. Wei, W. Ge, H. Wang, C. Davis, K. I. Fisher-Aylor, A. Mortazavi, M. Gerstein, T. Gingeras, B. Wold, Y. Sun, M. J. Fullwood, E. Cheung, E. Liu, W.-K. Sung, M. Snyder, and Y. Ruan, Cell **148**, 84 (2012).
[25] A. J. Larsson, P. Johnsson, M. Hagemann-Jensen, L. Hartmanis, O. R. Faridani, B. Reinius, Å. Segerstolpe, C. M. Rivera, B. Ren, and R. Sandberg, Nature **565**, 251 (2019).
[26] W.-K. K. Cho, N. Jayanth, B. P. English, T. Inoue, J. O. Andrews, W. Conway, J. B. Grimm, J.-H. H. Spille, L. D. Lavis, T. Lionnet, and I. I. Cisse, eLife **5** (2016), 10.7554/eLife.13617.
[27] C. K. Damgaard, S. Kahns, S. Lykke-Andersen, A. L. Nielsen, T. H. Jensen, and J. Kjems, Molecular Cell **29**, 271 (2008).
[28] A. B. Eberle, S. Lykke-Andersen, O. Mühlemann, and T. H. Jensen, Nature Structural and Molecular Biology **16**, 49 (2009).
[29] F. A. Ran, P. D. Hsu, J. Wright, V. Agarwala, D. A. Scott, and F. Zhang, Nature Protocols **8**, 2281 (2013).
[30] F. Mueller, A. Senecal, K. Tantale, H. Marie-Nelly, N. Ly, O. Collin, E. Basyuk, E. Bertrand, X. Darzacq, and C. Zimmer, Nature Methods **10**, 277 (2013).
[31] A. Spiess, "qpcR: Modelling and Analysis of Real-Time PCR Data," (2013), https://CRAN.R-project.org/package=qpcR.
[32] M. Martin, EMBnet.journal **17**, 10 (2011).
[33] A. R. Quinlan and I. M. Hall, Bioinformatics **26**, 841 (2010).
[34] A. Dobin, C. A. Davis, F. Schlesinger, J. Drenkow, C. Zaleski, S. Jha, P. Batut, M. Chaisson, and T. R. Gingeras, Bioinformatics **29**, 15 (2013).
[35] N. P. Dyer, V. Shahrezaei, and D. Hebenstreit, PeerJ **7**, e6222 (2019).
[36] S. Anders, P. T. Pyl, and W. Huber, Bioinformatics **31**, 166 (2015).
[37] F. Ramírez, F. Dündar, S. Diehl, B. A. Grüning, and T. Manke, Nucleic Acids Research **42**, W187 (2014).
[38] M. I. Love, W. Huber, and S. Anders, Genome Biology **15**, 550 (2014).
[39] L. D. Goldstein, Y. Cao, G. Pau, M. Lawrence, T. D. Wu, S. Seshagiri, and R. Gentleman, PLOS ONE **11**, e0156132 (2016).
[40] Y. Liao, G. K. Smyth, and W. Shi, Nucleic Acids Research **47**, e47 (2019).
